# Blocking necroptosis reduces inflammation and tumor incidence in a mouse model of diet-induced hepatocellular carcinoma

**DOI:** 10.1101/2022.08.03.502666

**Authors:** Sabira Mohammed, Nidheesh Thadathil, Albert L Tran, Michael Van Der Veldt, Constantin Georgescu, Nair H Haritha, Phoebe Ohene-Marfo, Sangphil Oh, Evan H Nicklas, Dawei Wang, Wenyi Luo, Ralf Janknecht, Benjamin F Miller, Jonathan D. Wren, Willard Freeman, Sathyaseelan S Deepa

## Abstract

**Background & Aims:** Nonalcoholic fatty liver disease (NAFLD) is one of the etiologies that contribute to hepatocellular carcinoma (HCC), and chronic inflammation is one of the proposed mediators of HCC. As necroptosis is a cell death pathway that induces inflammation, we tested whether necroptosis-induced inflammation contributes to the progression of NAFLD to HCC in a mouse model of diet-induced HCC.

**Methods:** Male and female wild-type (WT) mice or mouse models where necroptosis is blocked (*Ripk3*^-/-^ or *Mlkl*^-/-^ mice) were fed a control diet or choline-deficient low fat diet (CD-LFD) or CD-high fat diet (CD-HFD) for 6 months. Changes in inflammation, immune cell infiltration, activation of oncogenic pathways, and tumor incidence were assessed by gene expression analysis, western blotting, and flow cytometry. RNA sequencing (RNA-seq) was performed to assess the changes in liver transcriptome.

**Results:** Blocking necroptosis by deleting either *Ripk3* or *Mlkl* reduced markers of inflammation [proinflammatory cytokines (TNFα, IL-6, and IL-1β), F4/80^+ve^ macrophages, CCR2^+ve^ infiltrating monocytes], inflammation associated oncogenic pathways (JNK, PD-L1/PD-1, β-catenin), and HCC in male mice. In female mice, blocking necroptosis reduced HCC independent of inflammation. Blocking necroptosis reduced cell senescence markers in males and females, suggesting a novel cross-talk between necroptosis and cell senescence.

**Conclusions:** Our data show that hepatic necroptosis promotes recruitment and activation of liver macrophages leading to chronic inflammation, which in turn trigger oncogenic pathways leading to the progression of NAFLD to HCC in male mice. In female mice necroptosis contributes to HCC independent of inflammation. Thus, our study suggests that necroptosis is a valid target for NAFLD-mediated HCC.

**Synopsis:** Necroptosis is a cell death pathway that mediate inflammation. Blocking necroptosis attenuated chronic inflammation by reducing recruitment and activation of liver macrophages, which in turn reduced activation of oncogenic pathways and progression of NAFLD to HCC in mice.

**Graphical Abstract:** 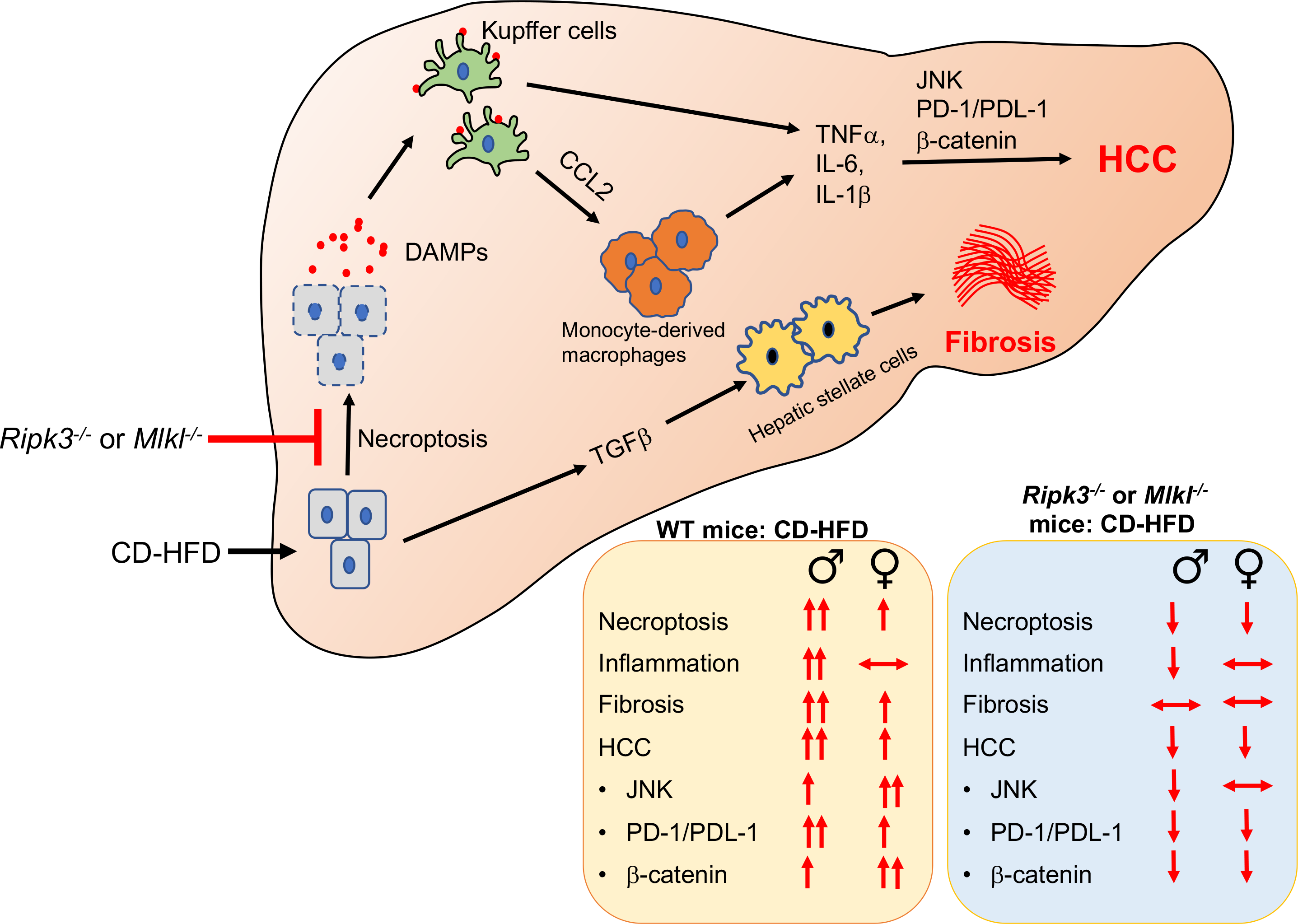

## Introduction

Hepatocellular carcinoma (HCC), the primary form of liver cancer, is a global health challenge that is predicted to affect more than 1 million people annually by the year 2025. HCC is the fourth cancer-related cause of death worldwide, and deaths from HCC are rising faster than any other cancer in the United States. Therapeutic options for HCC are limited, and survival after diagnosis is poor (<10%).^1^ Nonalcoholic fatty liver disease (NAFLD) is one of the etiologies that contribute to HCC and the prevalence of NAFLD-related HCC in the United States is expected to rise by over 100% by 2030 compared to 2016. NAFLD covers a spectrum of diseases ranging from fat deposition in the liver (non-alcoholic fatty liver, NAFL) to non-alcoholic steatohepatitis (NASH) that is characterized by steatosis, increased hepatic inflammation, fibrosis, and hepatocyte death. Nearly 30% of people with NAFL progress to NASH, and from these, 2-13% progress to HCC with cirrhosis and 8% progress to HCC without cirrhosis.^2^ Despite the strong association between NAFLD and HCC, the pathway(s) that cause the progression of NAFLD to HCC are not clearly understood.

Non-resolving chronic inflammation is believed to be a contributor to the development and progression of HCC. One of the pathways that triggers a persistent inflammatory response is the activation of innate immune cells by pathogen associated molecular patterns (PAMPs) or damage associated molecular patterns (DAMPs).^3^ Hepatic macrophages, the major innate immune cells in the liver, contain pattern recognition receptors (PRR, such as toll-like receptors) for DAMPs and PAMPs, and their binding to PRRs leads to increased production of proinflammatory cytokines.^4^ DAMPs such as high mobility group box protein 1 (HMGB1) and mitochondrial DNA released from hepatocytes are proposed to be mediators of chronic inflammation in NASH, and levels of these DAMPs are increased in mouse models and patients with NASH.^5^ Importantly, liver-specific HMGB1 deficiency reduces HCC development in mouse models of chronic liver injury.^6^ Thus, liver inflammation induced by DAMPs plays an important role in NASH and HCC.

One of the pathways that release DAMPs from cells is necroptosis, a regulated form of cell death that has been shown to lead to inflammation.^7^ Clinical data and studies using animal models suggest that hepatocyte death is the key trigger of liver disease progression, and hepatocytes are one of the major cell types that release DAMPs during hepatic injury.^8^ Necroptosis is initiated by necroptotic stimuli that sequentially phosphorylate and activate receptor-interacting protein kinase 1 (RIPK1), RIPK3, and the pseudokinase mixed lineage kinase domain-like (MLKL) protein. Phosphorylation of MLKL leads to its oligomerization and membrane attachment, which results in permeabilization of the cell membrane and DAMP release.^9^ Necroptosis has been reported to be increased in the livers of NAFLD and NASH patients and in mouse models of NASH.^10, 11^ Cell culture studies have shown that treating primary hepatocyte cultures with palmitic acid induces necroptosis, suggesting that lipotoxicity might be a trigger for necroptosis in NAFLD.^12, 13^

Previously, we reported that inhibiting necroptosis using necrostatin-1s (Nec-1s) in a mouse model of spontaneous HCC (mice deficient in the antioxidant enzyme Cu/Zn superoxide dismutase, *Sod1^-/-^* mice) reduced hepatic inflammation, fibrosis and pathways associated with HCC development.^14^ We also found that inhibiting necroptosis reduced hepatic inflammation and fibrosis in old wild type mice that exhibit NASH pathology.^15^ Based on these findings, we hypothesized that necroptosis mediated inflammation contributes to the progression of NAFLD to HCC. To test our hypothesis, we used a mouse model of NASH-induced HCC that develops robust fibrosis and HCC when mice are fed a choline deficient amino acid defined high fat diet (CD-HFD) for 6 months.^16^ Necroptosis was blocked using either *Ripk3^-/-^* or *Mlkl^-/-^*mice. Because the incidence of NAFLD and HCC is higher in men than in women, both male and female mice were used in our study.^17^ Our data show a sex-specific difference in the development of inflammation, fibrosis and HCC in WT mice. However, blocking necroptosis reduced HCC in both males and females without altering liver fibrosis.

## Results

### CD-HFD increased necroptosis in the liver and deleting either *Ripk3* or *Mlkl* reduced necroptosis

The gain in body weight for either *Ripk3^-/-^* or *Mlkl^-/-^*mice fed NC was significantly lower than WT male mice fed NC, however, body weight for the three groups of mice were similar for female mice fed NC (**Fig. S1A**). CD-LFD significantly reduced body weight gain for WT, *Ripk3^-/-^* and *Mlkl^-/-^* male mice relative to NC, and CD-HFD further exacerbated this effect (**Fig. S1A**). In contrast, CD-LFD or CD-HFD had no effect on body weight gain in female mice relative to NC, except for *Mlkl^-/-^*mice fed CD-HFD, which gained body weight (**Fig. S1A**). No significant difference in food intake was observed for male or female mice on the three diets. Feeding NC did not alter percentage liver weight (liver weight normalized to body weight) of all 3 groups of male or female mice whereas feeding CD-LFD resulted in a significant increase in percentage liver weight (∼1.5 to 2-fold for male mice and ∼1.5 fold for female mice), except for male *Mlkl^-/-^* mice. CD-HFD also increased percentage liver weight in all 3 groups of male and female mice (**Fig. S1B**). Raw liver weights also showed a similar trend (**Fig. S1C**). Thus, unlike the sex specific differences in body weight, no significant difference in liver weight was observed in male or female mice in response to the diets.

In both male and female WT mice, CD-LFD did not increase the levels of MLKL oligomers, a marker of necroptosis, over that observed with the NC diet, however, CD-HFD resulted in a significant increase in MLKL oligomers in both male (∼7.5-fold) and female (∼5-fold) WT mice relative to mice fed either NC or CD-LFD (**Fig. 1A**). Importantly, deleting either *Ripk3* or *Mlkl* significantly reduced MLKL oligomers in both male and female mice (**Fig. 1A**). MLKL protein levels were significantly increased in the livers of WT and *Ripk3^-/-^* mice in response to CD-LFD (∼5-fold in males and ∼4-fold in females) and CD-HFD (∼6-fold in males and ∼5-fold in females) (**Fig. 1A**), and was associated with an increase in *Mlkl* transcript levels in male and female mice in response to CD-LFD (∼3-fold for males and ∼2-fold for females) or CD-HFD (∼3.5-fold for males and ∼2-fold for females) (**Fig. S1D**). In WT male mice fed CD-HFD, RIPK3 protein levels were increased (∼2-fold), whereas in WT female mice both CD-LFD and CD-HFD increased RIPK3 expression (∼2-fold). Interestingly, absence of *Mlkl* blunted this effect in male mice (**Fig. 1A**). However, transcript levels of *Ripk3* were unaltered by the absence of *Mlkl* in male or female mice (**Fig. S1D**).

**Figure 1.**
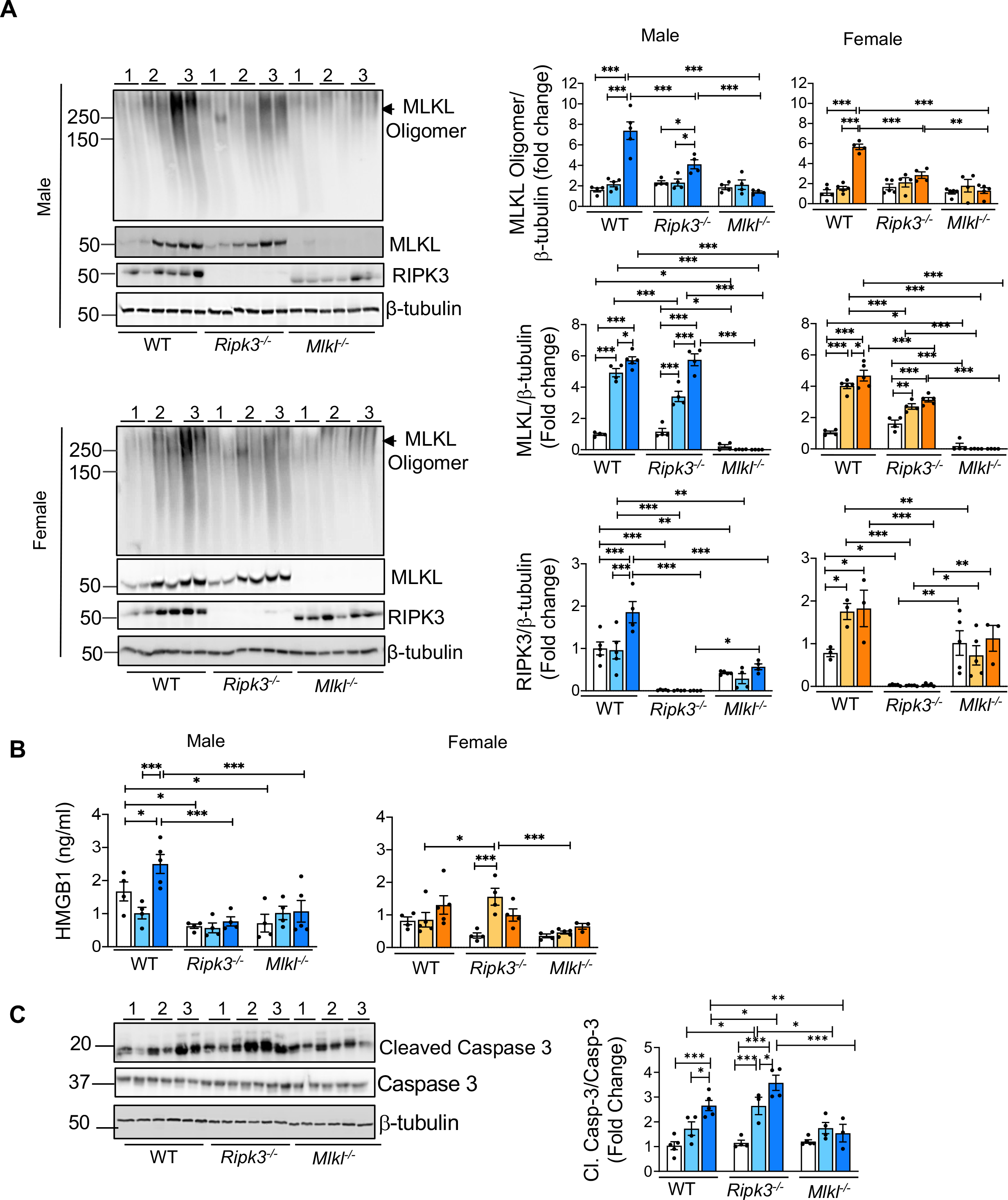
CD-HFD increased necroptosis in liver and deleting either *Ripk3* or *Mlkl* significantly reduced necroptosis. **(A)** *Left panel:* Immunoblots of liver tissue extracts for MLKL oligomers, MLKL, RIPK3, and β-tubulin from WT, *Ripk3^-/-^*, and *Mlkl^-/-^* male (top) and female (bottom) mice fed NC (1), CD-LFD (2) or CD-HFD (3). *Right panel:* Graphical representation of quantified blots normalized to β-tubulin. Males: NC (white bars), CD-LFD (light blue bars) or CD-HFD (dark blue bars). Females: NC (white bars), CD-LFD (light orange bars) or CD-HFD (dark orange bars). **(B)** Graphical representation of HMGB1 in plasma of WT, *Ripk3^-/-^*, and *Mlkl^- /-^* male (left panel) and female (right panel) mice fed NC or CD-LFD or CD-HFD. **(C)** *Left panel:* Immunoblots of liver tissue extracts for cleaved Caspase-3, Caspase-3, and β-tubulin from WT, *Ripk3^-/-^*, and *Mlkl^-/-^* male mice fed NC (1) or CD-LFD (2) or CD-HFD (3). *Right panel:* Graphical representation of quantified blots for cleaved Caspase-3 normalized to Caspase-3. Data are represented as mean±SEM, n=4-6 per group, *p<0.05, **p<0.01, ***p<0.001.

In WT male mice, CD-HFD but not CD-LFD resulted in a significant increase in circulating HMGB1, and blocking necroptosis (*Ripk3^-/-^* or *Mlkl^-/-^*mice) significantly reduced the levels of HMGB1 (**Fig. 1B**). In female mice, circulating HMGB1 showed a tendency to increase in response to CD-HFD; however, this increase did not reach statistical significance (**Fig. 1B**). Thus, increased MLKL oligomerization in response to CD-HFD in WT male mice parallels elevated HMGB1 levels.

CD-HFD resulted in a significant increase in cleaved Caspase-3, a marker of apoptosis, in the livers of WT male mice (∼3.5-fold) (**Fig. 1C**). Cleaved Caspase-3 levels were significantly elevated in response to CD-LFD (∼3-fold) and CD-HFD (∼4.3-fold) in *Ripk3^-/-^* male mice and were significantly higher than in WT mice. In contrast, cleaved Caspase-3 was not elevated in response to CD-LFD or CD-HFD in *Mlkl^-/-^* mice (**Fig. 1C**). Similar to male mice, CD-HFD increased the expression of cleaved Caspase-3 in WT female mice, however, it was ∼2-fold lower than in WT male mice. In female mice, knocking out neither *Ripk3* nor *Mlkl* had any effect on the expression of cleaved Caspase-3 (**Fig. S1E**).

### CD-HFD increased markers of inflammation in WT male mice and deleting either *Ripk3* or ***Mlkl* reduced inflammation in male mice.**

Increased expression of the proinflammatory cytokines IL6 and TNFα is reported to contribute to HCC development in response to a combination of chemical carcinogen and HFD in mice.^18^ Transcript levels of *IL6* (∼4-fold) in the liver, as well as levels of circulating IL6 (∼2-fold), were significantly increased in WT male mice in response to CD-HFD, whereas such an effect was not observed in WT female mice (**Figs. 2A, 2B**). Importantly, the transcript levels of hepatic *IL6* and circulating IL6 in male mice was significantly reduced in either *Ripk3^-/-^* or *Mlkl^-/-^* mice (**Figs. 2A, 2B**). Similarly, hepatic *TNF*α (∼2-fold) was elevated only in WT male mice in response to CD-HFD and this effect was blunted in either *Ripk3^-/-^* or *Mlkl^-/-^* mice (**Fig. 2C**). Circulating TNFα was elevated in male and female WT mice in response to CD-HFD and TNFα levels were reduced in either *Ripk3^-/-^* or *Mlkl^-/-^* male mice, but not in female mice (**Fig. 2D**). Similar to IL6 and TNFα, transcript levels of hepatic *IL-1β* were significantly elevated in liver in response to CD-HFD in males, but not females, and *IL-1β* levels were significantly reduced in either *Ripk3^-/-^* or *Mlkl^-/-^* mice (**Fig. S2A)**.

**Figure 2.**
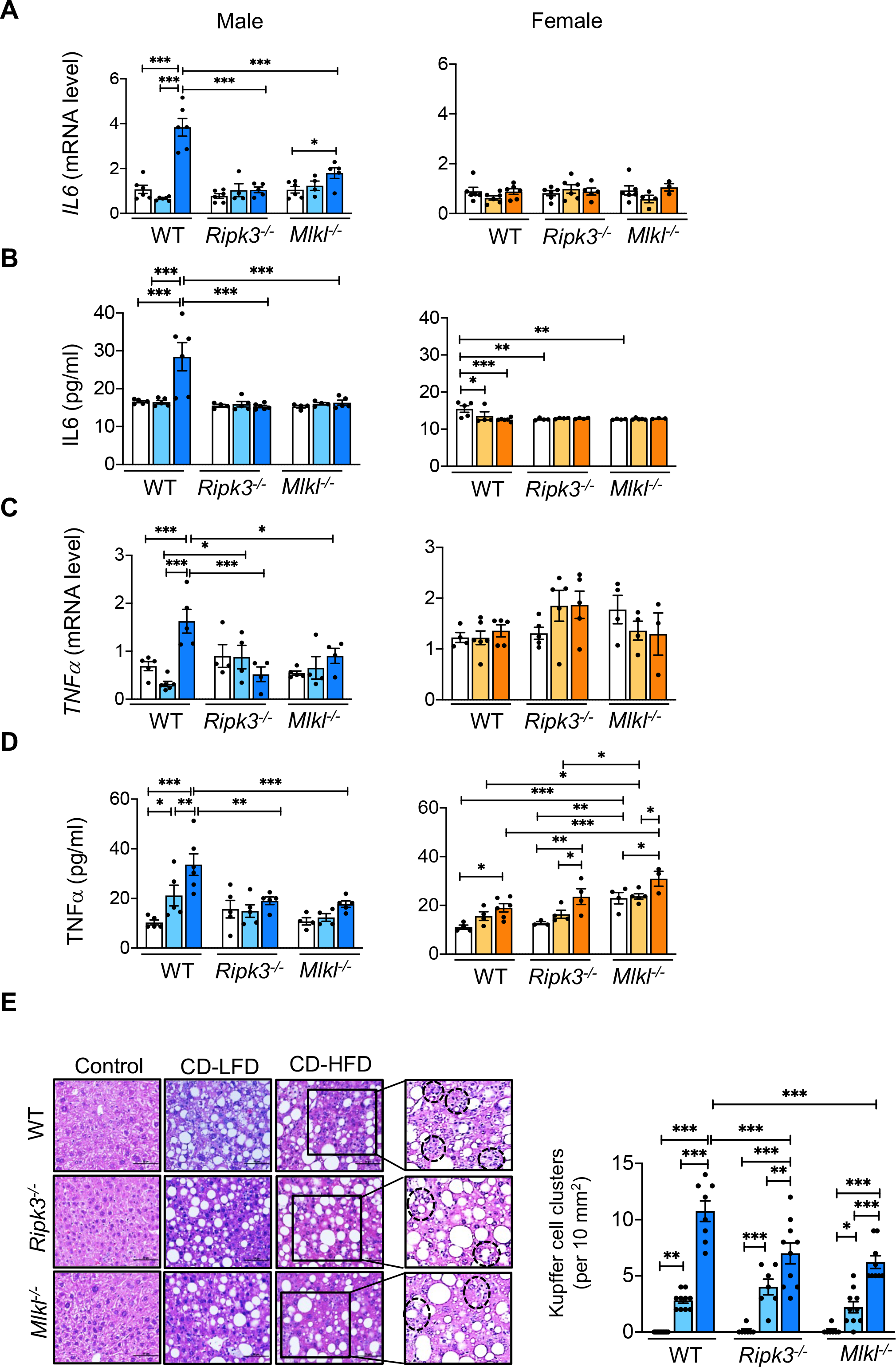
CD-HFD increased proinflammatory cytokine expression in WT mice in a sex- specific manner and deleting either *Ripk3* or *Mlkl* reduced expression of proinflammatory cytokines. Transcript levels of hepatic IL6 (A), circulating IL6 (B), transcript levels of hepatic TNFα (C), and circulating TNFα (D) in WT, *Ripk3^-/-^*, and *Mlkl^-/-^* male (*left panel*) and female (*right panel*) mice fed NC or CD-LFD, or CD-HFD. (E) *Left panel:* Images of H&E stained sections from WT, *Ripk3^-/-^*, and *Mlkl^-/-^* male mice fed NC or CD-LFD or CD-HFD. Black dotted circles represent KC clusters. Scale bar: 50 μM. *Right panel:* Graphical representation of the number of KC clusters in each group. Data are represented as mean±SEM, n=5-7 per group, *p<0.05, **p<0.01, ***p<0.001. Males: NC (white bars), CD-LFD (light blue bars) or CD-HFD (dark blue bars). Females: NC (white bars), CD-LFD (light orange bars) or CD-HFD (dark orange bars).

Histological comparison of H&E stained liver sections showed a significant increase in KC clusters, a measure of the proinflammatory status of the liver, in response to CD-LFD (∼3- to 5- fold) and CD-HFD (∼9- to 10-fold) in WT male mice (**Fig. 2E**).^19^ Blocking necroptosis using either *Ripk3^-/-^*or *Mlkl^-/-^* male mice did not reduce KC clusters in response to CD-LFD, whereas KC clusters were significantly reduced in response to CD-HFD (**Fig. 2E**). KC clusters were significantly elevated in WT female mice in response to CD-HFD (∼7.5-fold), however, this was ∼1.5-fold lower than in male mice, and KC clusters showed a significant reduction in *Ripk3^-/-^* female mice, but not in *Mlkl^-/-^* female mice (**Fig. S2B**).

Analysis of total immune cell population (percentage of CD45^+ve^ cells) showed that percentage of CD45^+^ cells are significantly elevated in the livers of WT male mice fed CD-HFD (∼3-fold), and this effect was significantly reduced in either *Ripk3^-/-^* or *Mlkl^-/-^* mice (**Fig. S3A**). In contrast to male mice, WT female mice showed a significant reduction in the CD45^+ve^ population (∼3-fold) in response to CD-HFD (**Fig. S3A**). Whereas the CD45^+ve^ population in CD-HFD fed *Ripk3^-/-^* female mice was similar to that in CD-HFD fed WT female mice, CD-HFD fed *Mlkl^-/-^*female mice showed a significant increase (∼2-fold) in CD45^+ve^ population (**Fig. S3A**). The gating strategy that was followed for the analysis is shown in **Fig S3B**. Analysis of total macrophage population, the major immune cell mediating hepatic inflammation in response to DAMPs, by staining cells with F4/80 showed a significant increase in the F4/80^+ve^ population in WT male mice in response to a CD-HFD (∼2-fold) relative to control diet. F4/80^+ve^ cell population was significantly reduced in either *Ripk3^-/-^* or *Mlkl^-/-^* male mice fed CD-HFD (**Figs. 3A, 3B**). In contrast, no significant change in F4/80^+ve^ population was observed in female mice in response to the diets (**Figs. 3A, 3B**). Because levels of macrophages derived from infiltrating monocytes (CCR2^+ve^ cells) increase during liver injury and are highly proinflammatory in nature, levels of CCR2^+ve^ cells were also assessed.^20^ A significant increase in CCR2^+ve^ cells was observed in WT male mice in response to CD-HFD (∼2.5-fold) and CCR2^+ve^ cells were significantly reduced in either *Ripk3^-/-^* or *Mlkl^-/-^* male mice (**Figs. 3C, 3D**). Similar to WT male mice, CD-HFD fed WT female mice also had a ∼2.5-fold increase in CCR2^+ve^ cells relative to control diet, however, their levels were 2-fold lower than in male mice. Importantly, blocking necroptosis using neither *Ripk3^-/-^* nor *Mlkl^-/-^* female had any effect on CCR2^+ve^ cells (**Figs. 3C, 3D**).

**Figure 3.**
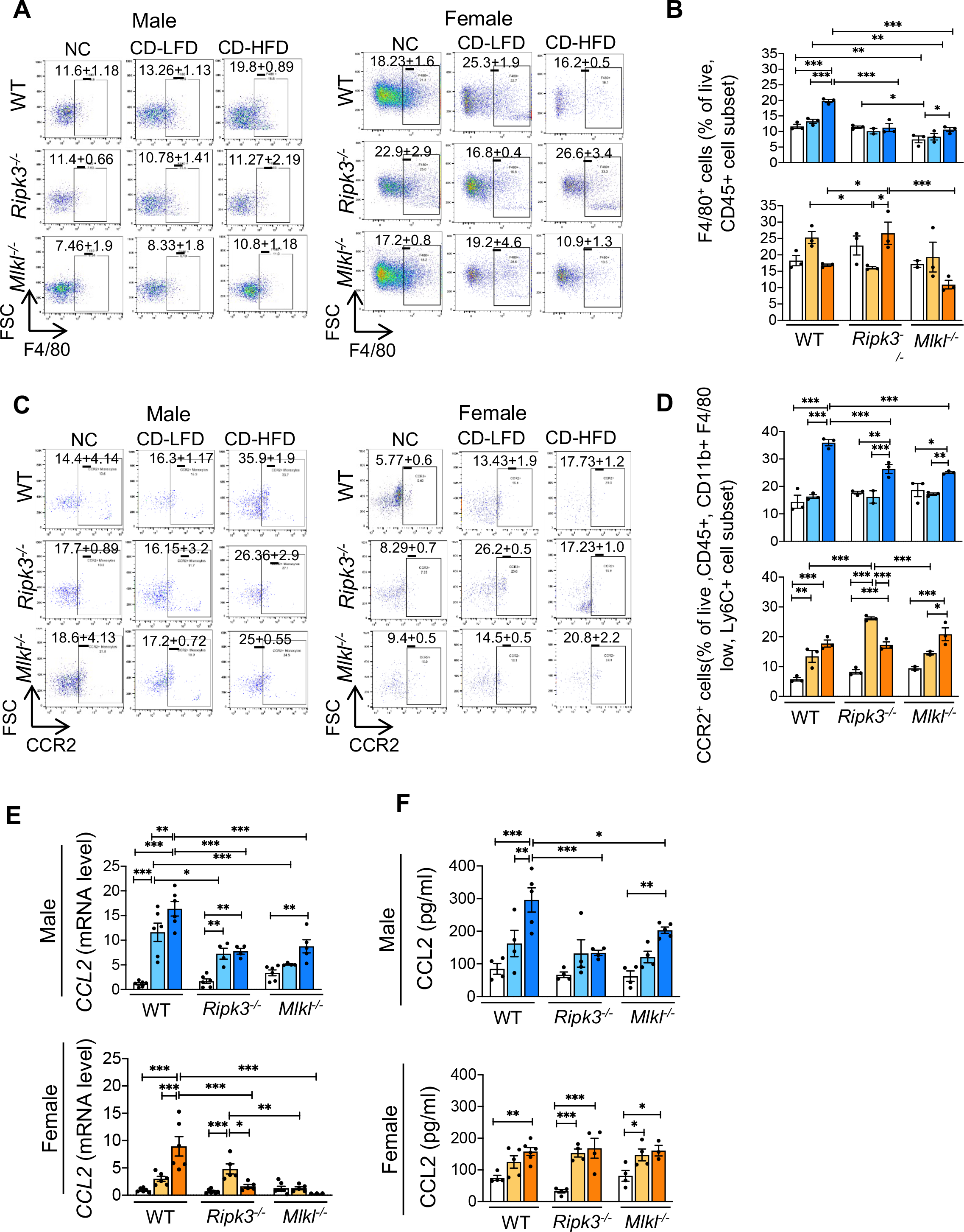
CD-HFD increased levels of inflammatory cell types and CCL2 in WT mice in a sex-specific manner and deleting either *Ripk3* or *Mlkl* reduced their levels. *Left panel:* Flow cytometric analysis of percentage of F4/80^+ve^ cell population (A) and CCR2^+ve^ cell population (C) in the livers of WT, *Ripk3^-/-^*, and *Mlkl^-/-^* male (left) and female (right) mice fed NC or CD-LFD or CD-HFD. *Right panel:* Graphical representation of the percentage population of F4/80^+ve^ cells gated on liver CD45^+ve^ cell subset (B) and CCR2^+ve^ cells, gated on liver CD45^+^, CD11b^+^, F4/80^low^, Ly6C^+^ cell subsets (D) in male (top) and female (bottom) mice. (E) Transcript levels of hepatic CCL2 and circulating CCL2 (F) in WT, *Ripk3^-/-^*, and *Mlkl^-/-^* male (top) and female (bottom) mice fed NC or CD-LFD or CD-HFD. Data are represented as mean±SEM, n=4-6 per group, *p<0.05, **p<0.01, ***p<0.001. Males: NC (white bars), CD-LFD (light blue bars) or CD-HFD (dark blue bars). Females: NC (white bars), CD-LFD (light orange bars) or CD-HFD (dark orange bars).

Transcript levels of chemokine (C-C motif) ligand 2 (*CCL2*), a chemokine that recruits monocytes to the sites of injury, were significantly increased in the livers of WT male (∼16-fold) and female (∼9-fold) mice in response to CD-HFD and these levels were significantly reduced in male and female *Ripk3^-/-^* or *Mlkl^-/-^* mice (**Fig. 3E**).^20^ Similarly, circulating CCL2 was elevated in response to CD-HFD in WT male and female mice, however, CCL2 levels were significantly reduced in either *Ripk3^-/-^* or *Mlkl^-/-^* male mice, but not in female mice (**Fig. 3F**). We also assessed transcript levels of markers of proinflammatory M1 macrophages. *CD68* (16-fold) and *TLR4* (5- fold) were significantly elevated in WT male mice in response to CD-HFD and these markers of M1 macrophages were significantly reduced in either *Ripk3^-/-^*or *Mlkl^-/-^* male mice (**Fig. S3C**).

### CD-HFD increased markers of liver fibrosis in WT mice and deleting either *Ripk3* or *Mlkl* **did not alter liver fibrosis.**

Assessment of fibrosis by picrosirius red (PSR) staining of liver tissue sections showed higher (∼2-fold) staining intensity in response to CD-HFD relative to CD-LFD in both males and females. PSR staining was higher in CD-HFD fed WT male mice (∼3-fold) than in females and blocking necroptosis using neither *Ripk3^-/-^* nor *Mlkl^-/-^* mice had any effect on PSR staining in male or female mice (**Fig. 4A**). Feeding WT mice CD-HFD increased the transcript levels of *Col3*α1 (∼28-fold for males and ∼12-fold for females), *Acta2* (∼3-fold for males and ∼4-fold for females), and *Col1α1* (∼46-fold for males and ∼30-fold for females), and compared to control diet (**Figs. 4B, 4C, S4A**). In male mice, blocking necroptosis by deleting *Ripk3* significantly reduced transcript levels of *Col3α1*, *Col1α1*, and *Acta2*, whereas only *Col1α1* showed a significant reduction in *Mlkl^-/-^* mice in response to CD-HFD. In female mice, absence of *Ripk3* or *Mlkl* had no effect on transcript levels of *Col3α1* or *Col1α1* or *Acta2* (**Figs. 4B, 4C, S4A**). Assessment of hepatic stellate cell (HSC) markers showed that desmin was elevated in WT mice in response to CD-LFD (∼4-fold for males and ∼2-fold for females) and CD-HFD (∼5-fold for males and ∼2-fold for females) (**Figs. 4D, S4B**). However, α-smooth muscle actin (α-SMA) levels were induced in WT mice only in response to CD-HFD (∼2-fold for males and females). Desmin and α-SMA levels in response to CD-HFD were reduced significantly in male *Ripk3^-/-^* mice, but not in female *Ripk3^-/-^* mice relative to WT mice, however, such a reduction was not observed for *Mlkl^-/-^* mice (**Figs. 4D, S4B**). *TGF-β* transcript levels were significantly upregulated in WT male and female mice (∼2-fold) in response to CD-HFD and blocking necroptosis using either *Ripk3*^-/-^ or *Mlkl*^-/-^ mice had no effect on *TGF-β* levels in male or female mice (**Fig. 4E, S4C**).

**Figure 4.**
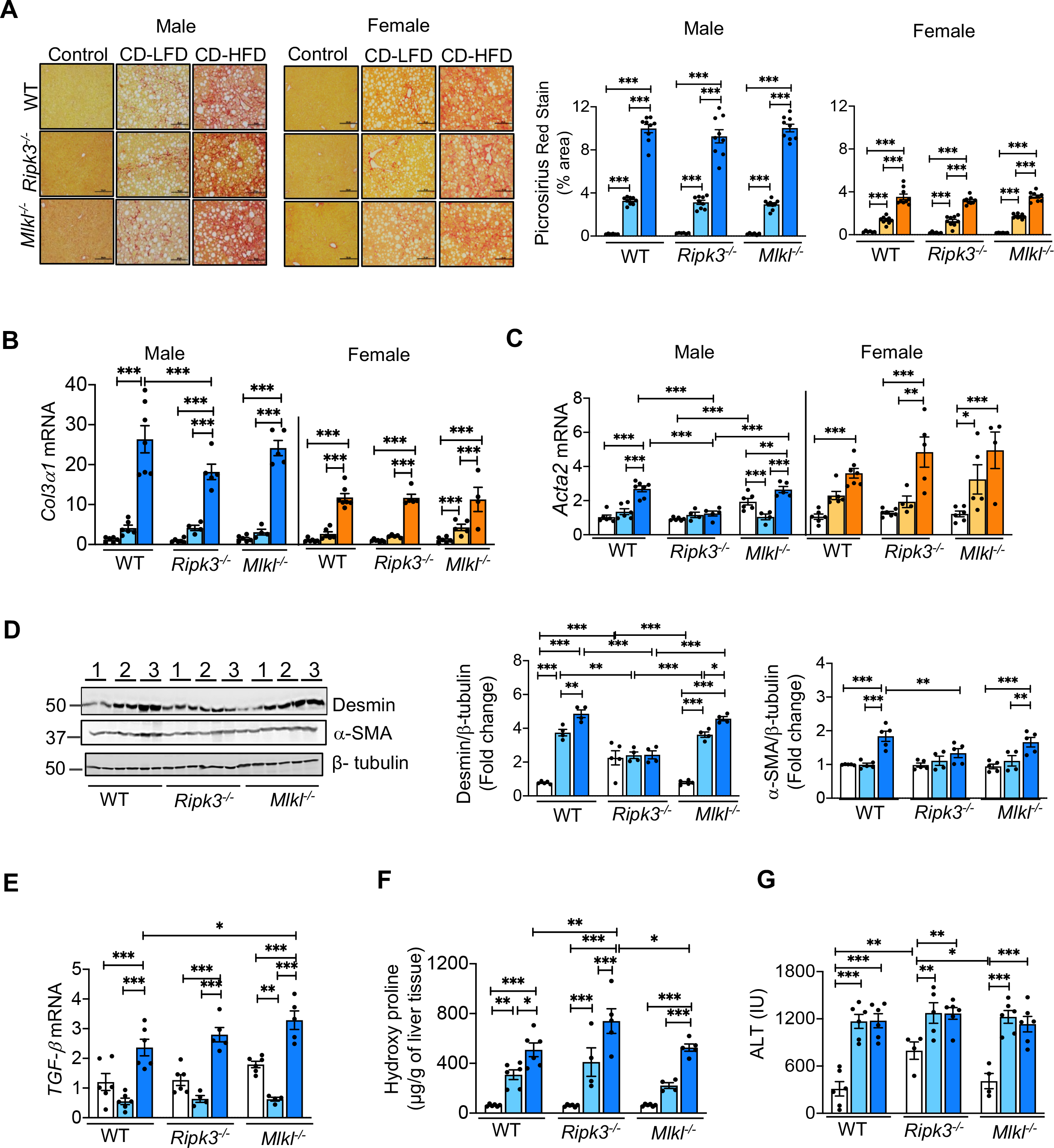
CD-HFD increased markers of liver fibrosis in WT mice and deleting either *Ripk3* or *Mlkl* did not alter liver fibrosis. **(A)** *Left panel:* PSR staining of liver sections of WT, *Ripk3^-/-^*, and *Mlkl^-/-^* male (left) and female (right) mice fed NC or CD-LFD or CD-HFD. Scale bar: 50 μM. *Right panel:* Graphical representation of PSR quantification. Transcript levels of *Col3α1* **(B)** and *Acta2* **(C)** in WT, *Ripk3^-/^*,*^-^* and *Mlkl^-/-^* male mice (left) and female mice (right) fed NC or CD- LFD or CD-HFD. **(D)** *Left panel:* Immunoblots of liver tissue extracts for desmin, α-SMA, and β- tubulin from WT, *Ripk3^-/-^*, and *Mlkl^-/-^* male mice fed NC (1) or CD-LFD (2) or CD-HFD (3). *Right panel:* Graphical representation of quantified blots normalized to β-tubulin. Transcript levels of hepatic *TGF-β* normalized to β-microglobulin. **(E)**, liver hydroxyproline content **(F)**, and serum ALT levels **(G)** in WT, *Ripk3^-/-^*, and *Mlkl^-/-^* male mice fed NC or CD-LFD or CD-HFD. Data are represented as mean±SEM, n=4-6 per group, *p<0.05, **p<0.01, ***p<0.001. Males: NC (white bars), CD-LFD (light blue bars) or CD-HFD (dark blue bars). Females: NC (white bars), CD-LFD (light orange bars) or CD-HFD (dark orange bars).

Assessment of total collagen content by measuring concentration of hydroxyproline (OHP) showed that OHP content was significantly elevated in the livers of WT male mice in response to CD-LFD (∼5-fold) and CD-HFD (∼8-fold) (**Fig. 4F**). OHP levels were not reduced in either *Ripk3*^-/-^ or *Mlkl*^-/-^ male mice fed a CD-HFD compared to WT mice. Circulating levels of ALT, a biomarker of hepatocellular damage, were also significantly elevated in WT male mice in response to CD-LFD and CD-HFD (∼4-fold) and ALT levels in response to CD-LFD or CD-HFD did not differ significantly between WT and either *Ripk3*^-/-^ or *Mlkl*^-/-^ male mice (**Fig. 4G**).

### Deleting either *Ripk3* or *Mlkl* significantly reduced HCC development in response to a CD- HFD

WT male mice fed the CD-HFD developed liver nodules (2-25 nodules/mouse) while no or 1-2 liver nodules were detected in WT mice fed the control or CD-LFD diets (**Fig. 5A, S5A**). A similar effect was observed in female mice; however, the number of liver nodules was lower (∼2- to 3-fold) in female mice relative to male mice (**Fig. 5A**). Importantly, the number of liver nodules was significantly reduced in both *Ripk3*^-/-^ and *Mlkl*^-/-^ male and female mice fed the CD-HFD (**Fig. 5A). Fig. 5B** shows that the liver nodule size is also reduced in the *Ripk3*^-/-^ or *Mlkl*^-/-^ male mice compared to WT mice fed CD-HFD. For example, 12% of the nodules were large (>5 mm) in WT male mice fed CD-HFD compared to 0 and 5% for *Ripk3^-/-^* and *Mlkl^-/-^*mice, respectively. Large size nodules in the liver of female mice decreased from 2% in WT to no large tumor nodules being detected in either *Ripk3^-/-^* or *Mlkl^-/-^* female mice (**Table 1).** The tumor (T) region of liver sections from CD-HFD fed WT mice showed strong positive staining for glypican- 3, a marker for HCC, compared to non-tumor (NT) region and glypican-3 staining was reduced in both *Ripk3^-/-^* and *Mlkl^-/-^*mice (**Fig. 5C**). Tumors were further characterized using the cell proliferation marker, Ki-67, which showed that Ki-67 staining was increased in the tumor region of WT male mice in response to CD-HFD and blocking necroptosis by targeting *Ripk3^-/-^* and *Mlkl^-/-^* reduced Ki-67 levels (**Fig. 5D**). Next, levels of alpha-fetoprotein (AFP), a biomarker of HCC in serum and liver tissue, was measured. Serum AFP levels were significantly elevated (∼6-fold in males and ∼2-fold in females) in WT mice in response to CD-HFD, and serum AFP levels were significantly reduced in both *Ripk3^-/-^* and *Mlkl^-/-^* male and female mice (**Fig. 5E**). Transcript levels of liver AFP also showed a similar trend for male mice, but not for female mice (**Fig. S5B**).

**Figure 5.**
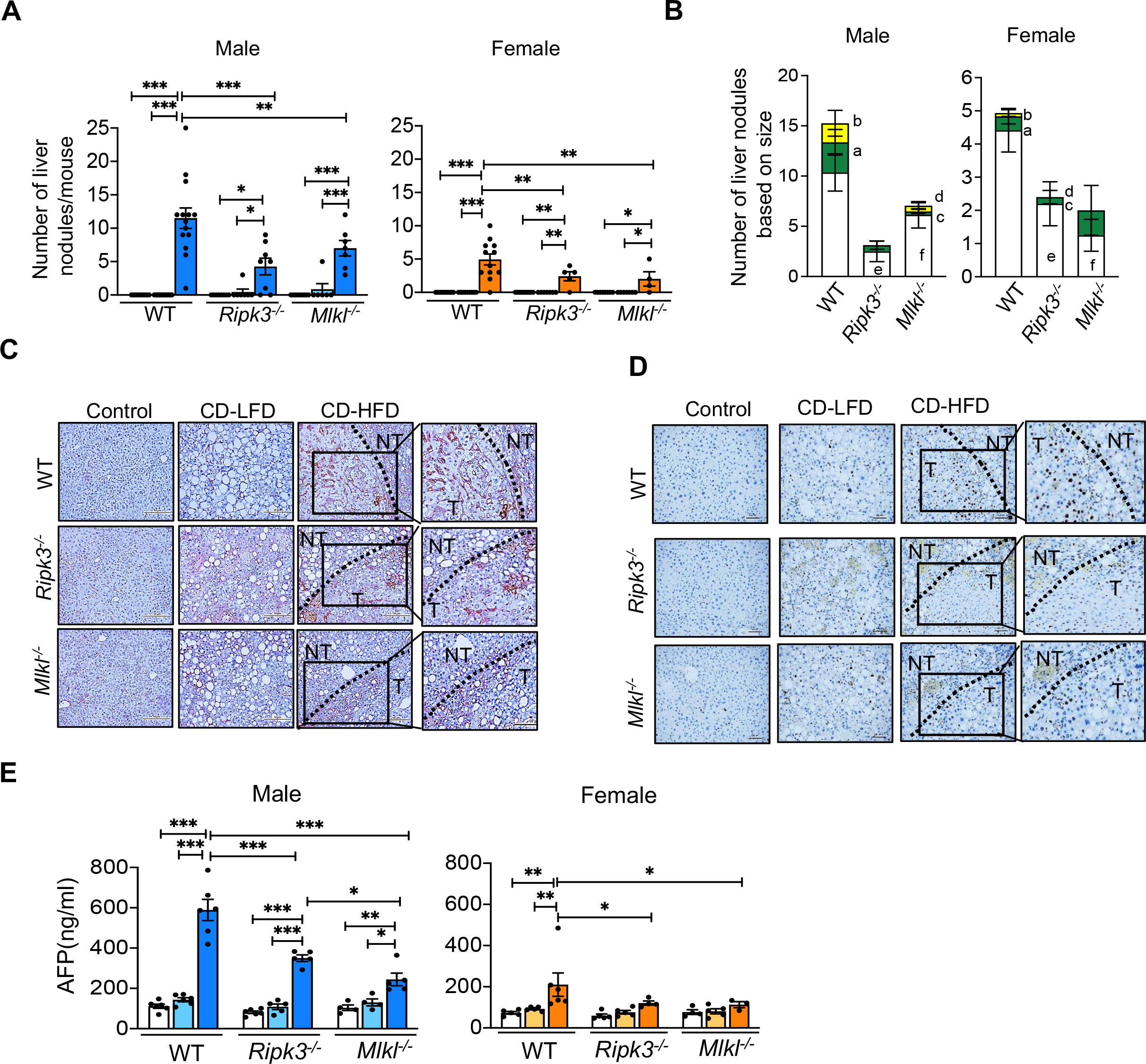
Deleting either *Ripk3* or *Mlkl* significantly reduced HCC development in response to a CD-HFD. (**A)** Tumor nodules in the livers of WT, *Ripk3^-/-^*, and *Mlkl^-/-^* male mice (left) and female mice (right) fed NC or CD-LFD or CD-HFD. **(B)** Graphical representation of large (>5mm, yellow), medium (2-4mm, green), and small (<1 mm, white) liver nodules in the livers of WT, *Ripk3^-/-^*, and *Mlkl^-/-^* male (left) and female (right) mice fed CD-HFD (^a^WT small tumor vs WT medium tumor, p<0.0001; ^b^WT small tumor vs WT large tumor, p<0.0001; ^c^*Mlkl^-/-^* small tumor vs *Mlkl^-/-^* medium tumor, p<0.01; ^d^*Mlkl^-/-^* small tumor vs *Mlkl^-/-^* large tumor, p<0.01; ^e^WT small tumor vs *Ripk3^-/-^*small tumor, p<0.0001; ^f^ WT small tumor vs *Mlkl^-/-^*small tumor, p<0.03). Immunohistochemical staining for glypican-3 **(C)** and Ki-67 **(D)** in the non-tumor (NT) and tumor (T) of liver sections from WT, *Ripk3^-/-^*, and *Mlkl^-/-^*male mice, fed NC or CD-LFD and CD-HFD. Dark red color in tumor (T) region indicate positive staining for glypican-3 and dark brown spots in tumor region indicate positive staining for Ki-67. Scale bar: 50 μM. **(E)** Circulating AFP levels in WT, *Ripk3^-/-^*, and *Mlkl^-/-^* male (left) and female (right) mice fed NC or CD-LFD or CD-HFD. Data are represented as mean±SEM, n=4-6 per group, *p<0.05, **p<0.01, ***p<0.001. Males: NC (white bars), CD-LFD (light blue bars) or CD-HFD (dark blue bars). Females: NC (white bars), CD-LFD (light orange bars) or CD-HFD (dark orange bars).

**Table 1:**
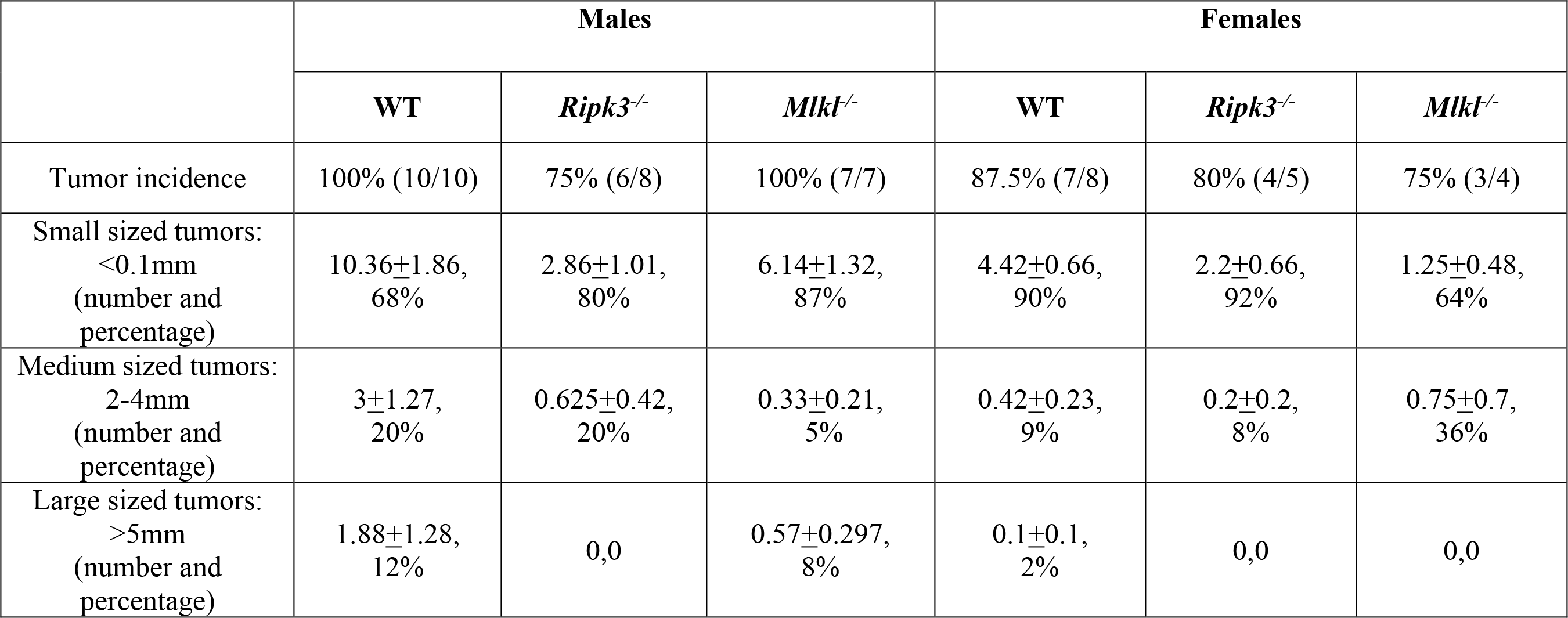
Comparison of percentage incidence of liver tumor with difference in sizes in WT, Ripk3^-/-^, and Mlkl^-/-^ male and female mice fed NC or CD-LFD, or CD-HFD.

### Gene up- or downregulation in response to CD-LFD or CD-HFD in WT mice was attenuated when either *Ripk3* or *Mlkl* was deleted

To determine the effect of blocking necroptosis on the liver transcriptome of male mice, we measured the levels of gene transcripts that change in response to the three diets in WT, *Ripk3^-/-^* and *Mlkl^-/-^* male mice by RNA-seq. Feeding mice the CD-LFD and CD-HFD resulted in a major change in the transcriptome of the liver, which was largely prevented in the *Ripk3^-/-^* and *Mlkl^-/-^* mice (**Fig. 6A)**. For WT mice fed CD-LFD, 1655 genes were altered (951-upregulated and 704-downregulated), and for WT mice fed CD-HFD, 3697 genes were altered (2212-upregulated, 1485-downregulated) relative to WT mice fed NC. A list of the transcripts that significantly changed is given in **Table S1**. To confirm the RNA-seq data, we selected genes from WT mice that were altered in response to CD-HFD, which were associated with tumorigenesis: *Tonsl, Dppa2*, and *Pogz.*^21–23^ qRT-PCR analysis showed that the mRNA levels of *Tonsl, Dppa2,* and *Pogz* were significantly increased in WT mice in response to CD-HFD and blocking necroptosis significantly reduced their expression in response to CD-HFD **(Fig. 6D)**.

**Figure 6.**
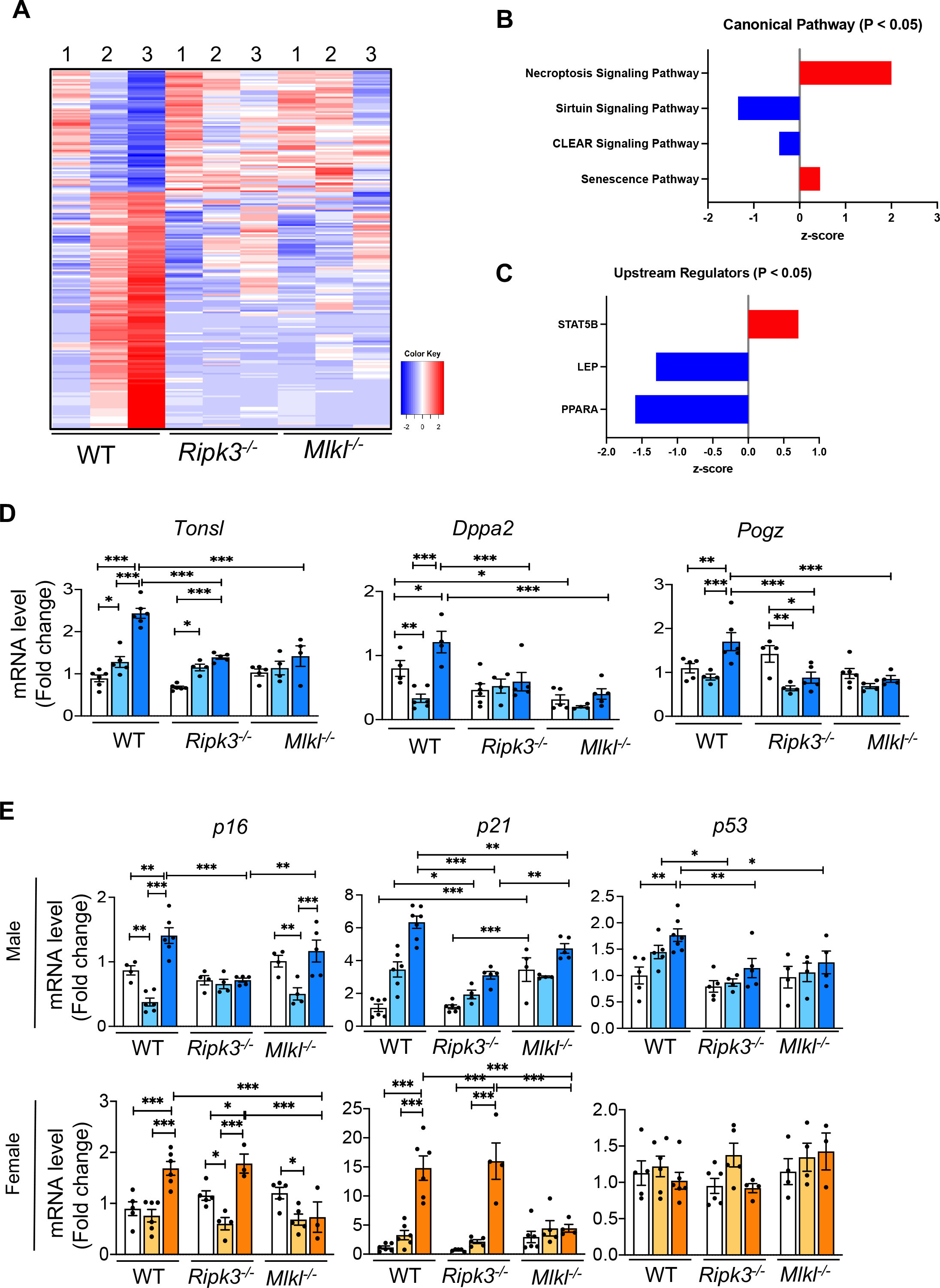
Genes upregulated or downregulated in response to CD-LFD or CD-HFD in WT mice were reversed when either *Ripk3* or *Mlkl* was deleted. (A) Schematic representation of transcriptome analysis of liver tissue extracts from WT, *Ripk3^-/^*, and *Mlkl^-/-^* male mice fed NC or CD-LFD or CD-HFD. Red color represents genes that are significantly upregulated and blue color represents genes that are significantly down regulated (1: NC, 2: CD-LFD, 3: CD-HFD). Graphical representation of canonical pathways which are significantly up-regulated (red) and down- regulated (blue) (B) upstream regulators that are significantly up-regulated (red) and down- regulated (blue) (C) in WT, *Ripk3^-/-^*, and *Mlkl^-/-^* male mice fed NC or CD-LFD or CD-HFD. (D) Transcript levels of *Tonsl*, *Dppa2*, and *Pogz* normalized to β-microglobulin in WT, *Ripk3^-/-^*, and *Mlkl^-/-^* male mice fed NC or CD-LFD or CD-HFD. (E) Transcript levels of *p16*, *p21,* and *p53* in male (top panel) and female (bottom panel) in WT, *Ripk3^-/-^*, and *Mlkl^-/-^* mice fed NC or CD-LFD or CD-HFD. Data are represented as mean±SEM, n=5-7 per group, *p<0.05, **p<0.01, ***p<0.001. Males: NC (white bars), CD-LFD (light blue bars) or CD-HFD (dark blue bars). Females: NC (white bars), CD-LFD (light orange bars) or CD-HFD (dark orange bars).

We performed further analysis of the transcriptome by conducting Ingenuity pathway analysis, which allows us to determine if genes in a pathway are altered. The ingenuity pathway analysis identified 24 pathways that were significantly (p<0.05) altered (**Fig. S6A**) and of these, two pathways showed positive z-score (necroptosis signaling pathway and senescence pathway), and two pathways showed negative z-score (sirtuin signaling pathway and CLEAR signaling pathway) (**Fig. 6B**). Assessment of markers of senescence showed that WT male mice fed CD-HFD showed a significant increase in *p16* (∼1.5-fold), *p21* (∼6-fold), and *p53* (∼2-fold) compared to WT mice fed NC. The transcript levels for p21 and p53 were significantly reduced in *Ripk3^-/-^* and *Mlkl^-/-^* male mice fed CD-HFD, and transcript levels for p16 were significantly reduced in the *Ripk3^-/-^*mice (**Fig. 6E**). In WT female mice, *p16* (∼2-fold) and *p21* (∼15-fold), but not *p53*, were significantly elevated in response to CD-HFD and blocking necroptosis reduced their expression in *Mlkl^-/-^* mice, but not in *Ripk3^-/-^* mice (**Fig. 6E**). Ingenuity pathway analysis further identified 15 upstream regulators of the pathways (**Fig. S6B**), of which one showed a positive z-score (STAT5) and two showed negative z-score (LEP and PPARα) (**Fig. 6C**).

### Oncogenic pathways involved in HCC promotion are induced in liver by the CD-HFD and deleting either *Ripk3* or *Mlkl* attenuated their induction

To identify pathways mediating HCC development in response to CD-HFD, we assessed activation of several HCC-associated pathways. Male mice were used for the initial studies and the pathways that were found to be activated in males were then validated in females. Because we used a high-fat diet, we first determined whether the Akt/mTOR pathway is activated. As shown in **Fig. S7A**, CD-HFD did not increase Akt phosphorylation (active form of Akt) and mTOR activation (as assessed by the phosphorylation of S6 kinase) in WT male mice. Next, we assessed MAPK pathways that are modulated by inflammation and are involved in HCC, e.g., JNK, ERK and p38 pathways, by assessing phosphorylation levels of these proteins which represent the active form of these proteins. Phosphorylation of JNK was increased ∼2-fold in the livers of CD-HFD fed WT male mice, and was significantly reduced in both *Ripk3^-/-^* and *Mlkl^-/-^* male mice (**Fig. 7A**). Similarly, JNK was activated in WT female mice livers in response to CD-HFD (∼3.5-fold), however, absence of *Ripk3* or *Mlkl* had no effect on JNK phosphorylation in female mice (**Fig. 7A**). P38 phosphorylation was unaffected by CD-HFD in WT male mice, whereas ERK phosphorylation was reduced in response to CD-HFD in WT male mice and their phosphorylation levels were unaltered in either *Ripk3^-/-^* or *Mlkl^-/-^* male mice (**Fig. S7B**).

**Figure 7.**
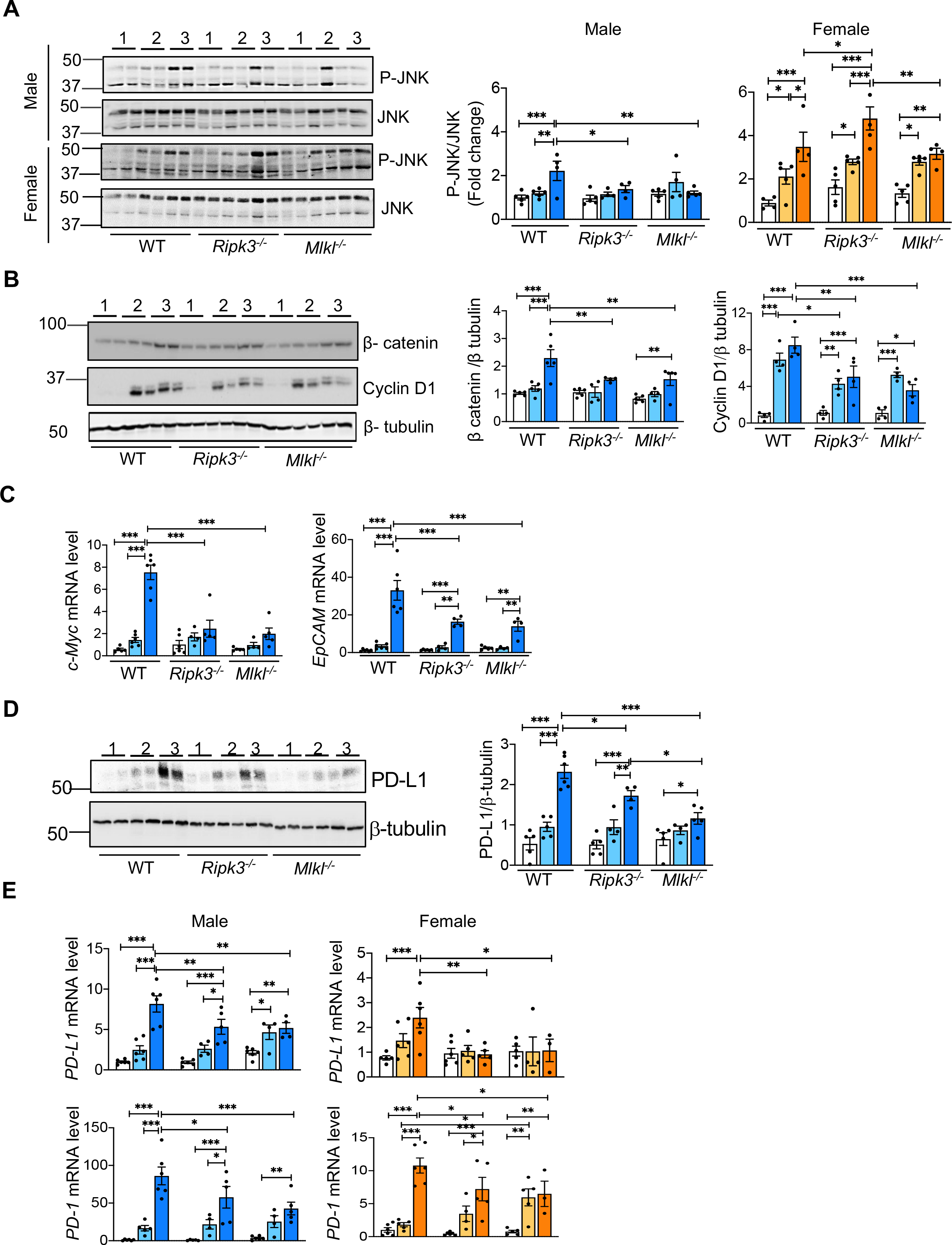
Several pathways involved in HCC are induced in liver by the CD-HFD and deleting either *Ripk3* or *Mlkl* attenuated their induction. (A) *Left panel:* Immunoblots of liver tissue extracts for P-JNK and JNK from WT, *Ripk3^-/-^*, and *Mlkl^-/-^* male (top) and female (bottom) mice fed with NC, CD-LFD or CD-HFD. *Right panel:* Graphical representation of quantified phospho-protein normalized to total protein. (B) Immunoblots of liver tissue extracts for β-catenin, cyclin-D1 and β-tubulin (left panel) from WT, *Ripk3^-/-^*, and *Mlkl^-/-^* male mice fed NC or CD-LFD or CD-HFD. *Right panel:* Graphical representation of quantified blots normalized to β-tubulin. (C) Transcript levels of *c-Myc* (left) and *EpCAM* (right) in WT, *Ripk3^-/-^* or *Mlkl^-/-^* male mice fed NC, CD-LFD or CD-HFD. (D) *Left panel:* Immunoblots of liver tissue extracts from WT, *Ripk3^-/-^*, and *Mlkl^-/-^* male mice fed NC or CD-LFD or CD-HFD, for PD-L1 and β-tubulin. *Right panel*: Graphical representation of quantified blots normalized to β-tubulin. (E) Transcript levels of hepatic *PD-L1* (top panel) and *PD-1* (bottom panel) normalized to β-microglobulin in WT, *Ripk3^-/-^*, and *Mlkl^-/-^* male (left) and female (right) mice fed NC or CD-LFD or CD-HFD. Data are represented as mean±SEM, n=4-6 per group, *p<0.05, **p<0.01, ***p<0.001. Males: NC (white bars), CD-LFD (light blue bars) or CD-HFD (dark blue bars). Females: NC (white bars), CD-LFD (light orange bars) or CD-HFD (dark orange bars).

The levels of β-catenin (∼2.5-fold) and its downstream target, cyclin D1 (∼5-fold) were significantly elevated in CD-HFD fed WT male mice and the levels of these proteins were significantly reduced in both *Ripk3^-/-^* and *Mlkl^-/-^* male mice (**Fig. 7B**). In WT female mice, CD- HFD increased β-catenin levels (∼4-fold) in the liver and was significantly reduced in *Ripk3^-/-^* and *Mlkl^-/-^* female mice fed CD-HFD (**Fig. S7C**). Transcript levels of *c-Myc* (∼8-fold) and *EpCAM* (∼35-fold), the downstream targets of the JNK and β-catenin pathways, were significantly elevated in WT male mice in response to CD-HFD, and their expression was significantly reduced in both *Ripk3^-/-^* and *Mlkl^-/-^*male mice fed CD-HFD (**Fig. 7C**). Expression of PD-L1 protein, the immune checkpoint inhibitor, was significantly upregulated (4-fold) in the livers of WT male mice in response to CD-HFD, and PD-L1 expression was significantly reduced in both *Ripk3^-/-^* and *Mlkl^-/-^* male mice fed CD-HFD with a greater reduction in *Mlkl*^-/-^ mice (**Fig. 7D**). Transcript levels of PD-L1 in the liver were also increased in male (∼8-fold) and female (∼2.5-fold) WT mice in response to CD-HFD, and PD-L1 transcript levels were significantly reduced in *Ripk3^-/-^* and *Mlkl^-/-^* male and female mice fed CD-HFD (**Fig. 7E**). Similarly, PD-1 transcript levels were significantly elevated in the livers of CD-HFD fed male (∼90-fold) and female WT (∼11-fold) mice, and PD-1 transcript levels were significantly reduced in both *Ripk3^-/-^* and *Mlkl^-/-^*male and female mice fed CD-HFD (**Fig. 7E)**. The levels of p62, a marker of autophagy that is elevated in HCC, was significantly up-regulated in the livers of WT male mice fed CD-HFD (4-fold); however, p62 levels in the livers of both *Ripk3^-/-^* and *Mlkl^-/-^* male mice fed CD-HFD were not different from WT mice (**Fig. S7D**). Interestingly, basal p62 expression in both *Ripk3^-/-^* and *Mlkl^-/-^* male mice fed normal chow diet were significantly higher (∼3-fold) than WT mice fed a normal chow.

## Discussion

This study addressed the role of two key proteins in necroptosis pathway, RIPK3 and MLKL, on hepatic inflammation and HCC development in a NASH-induced HCC mouse model; and the sex-specific effects of deleting *Ripk3* or *Mlkl* on inflammation, liver fibrosis, and HCC. The major findings of the study are that deleting either *Ripk3* or *Mlkl* ameliorated hepatic inflammation and HCC in male mice and HCC only in female mice. Liver fibrosis was however not improved in either male or female mice.

We used two approaches to genetically block necroptosis by targeting either *Ripk3* or *Mlkl*, which allowed us to rigorously test the role of necroptosis in HCC. While both RIPK3 and MLKL are key proteins involved in necroptosis, they also have an impact on other processes. For example, RIPK3 is involved in apoptosis, NLRP3 activation, and lipid metabolism,^24–26^ whereas MLKL is involved in autophagy and endosomal trafficking.^13, 27^ By showing that both *Ripk3* and *Mlkl* knockout mice had similar effects on reducing inflammation and/or HCC in response to CD-HFD, we have strong evidence that necroptosis plays a role in HCC induction in the CD-HFD mouse model.

Our study showing that necroptosis is elevated in the livers of WT male mice in response to CD-HFD agrees with previous studies that have reported increased necroptosis in the livers of male mice fed other NAFLD or NASH inducing diets such as CD diet or HFD or western diet.^10, 12, 26^ We found that blocking necroptosis by targeting either *Ripk3* or *Mlkl* reduced hepatic inflammation in male mice assessed by the reduced levels of proinflammatory cytokines, KC clusters, total macrophages (F4/80^+ve^ cells), and infiltrating CCR2^+ve^ monocytes. Liver macrophages, in particular the infiltrating monocytes derived macrophages, are the major and most important mediators of the inflammatory response in NASH and HCC.^28^ Thus, our finding that blocking necroptosis reduced total liver macrophages and infiltrating monocytes suggests a role of liver macrophages in necroptosis mediated inflammation in male WT mice. In contrast, WT female mice did not show an increase in the levels of proinflammatory cytokines (TNFα, IL6, and IL-β) or total macrophage numbers in the liver. Even though infiltrating CCR2^+ve^ monocytes were elevated in response to CD-HFD in WT female mice, this was 50% lower than the response in male mice. A recent study has shown that male mice accumulated more macrophages in adipose tissue than female mice leading to elevated levels of inflammatory cytokines.^29^ Therefore, it is possible that reduced macrophage accumulation in the livers of female mice could contribute to the reduction in hepatic inflammation relative to male mice. Yet another possibility is the presence of the female sex hormone, estrogen, which has been shown to inhibit production of proinflammatory cytokines by KC.^30^ Even though female mice did not exhibit an increase in the levels of proinflammatory cytokines, we did observe an increase in the level of necroptosis markers in the livers of WT female mice in response to CD-HFD, suggesting the possibility that plasma membrane rapture, the terminal step in necroptosis, is inhibited in female mice. Consistent with this observation, levels of HMGB1, which are released during necroptosis, were elevated in male mice, but not in female mice. A recent study has shown that ubiquitination of MLKL oligomers blocks the cytotoxic potential of MLKL.^31^ Therefore, future studies need to determine if MLKL oligomers in the livers of female mice undergo ubiquitination resulting in necroptosis inhibition.

Consistent with the increase in proinflammatory cytokines, pathways regulated by inflammatory cytokines in HCC promotion were also increased in the livers of WT male mice fed a CD-HFD, i.e. JNK, PD-L1/PD-1 signaling, and β-catenin pathways. JNK promotes hepatocyte proliferation through the activation of oncogenes, PD-1/PD-L1 immune checkpoint inhibitors have emerged as a promising strategy for the treatment of HCC, and the β-catenin pathway regulates several cellular processes that are involved in HCC.^32–34^ Deleting either *Ripk3* or *Mlkl* reduced JNK, PD-1/PD-L1 signaling, and β-catenin pathways in male mice, supporting a role of necroptosis-mediated inflammation in HCC in male mice. Afonso et al. (2021) have shown that blocking necroptosis by targeting *Ripk3* reduced β-catenin levels and HCC in response to CD diet.^26^ Whereas JNK, PD-1/PD-L1 signaling, and β-catenin pathways were also elevated in WT female mice in response to CD-HFD, blocking necroptosis reduced only PD-1/PD-L1 signaling and β-catenin pathways, suggesting that other factors or pathways in addition to inflammation can also modulate these pathways.

Our finding that deleting either *Ripk3* or *Mlkl* reduced hepatic inflammation and oncogenic pathways in male mice that paralleled the reduction in number and size of liver tumor nodules supports a role of necroptosis-mediated inflammation in HCC in male mice. Afonso et al. (2021) also reported reduced inflammation and HCC in *Ripk3^-/-^* mice fed a CD diet. Even though WT female mice developed HCC in response to CD-HFD, the number of tumor nodules were ∼2.5- lower than in WT male mice.^26^ This observation is consistent with the well documented fact that incidence of HCC in humans is >2-fold higher in males compared to females.^35^ A similar observation was reported in a mouse model of diethylnitrosamine (DEN)-induced hepatocarcinogenesis where males had higher incidence of HCC relative to females. In their study, Naugler et al. (2007) showed that DEN induced the production of IL-6 by KC, and ablation of IL- 6 in male mice eliminated the gender differences in hepatocarcinogenesis in mice, further supporting the role of proinflammatory cytokines in HCC promotion.^30^ Thus, high levels of proinflammatory cytokines in WT male mice observed in our study could contribute to increased HCC incidence in male mice. Surprisingly, similar to male mice, blocking necroptosis using either *Ripk3* or *Mlkl* reduced liver tumor nodules in females; however, the reduction occurred in the absence of any significant reduction in inflammation. This finding would suggest the existence of alternate pathways regulated by both *Ripk3* and *Mlkl* in HCC development in females. For example, our RNA seq data for male mice identified peroxisome proliferator-activated receptor α (PPARα), which plays a crucial role in the regulation of fatty acid oxidation in the liver, as an upstream regulator of pathways modulated by necroptosis. Expression of PPARα is lower in HCC in humans, and no significant difference in PPARα expression was observed in HCC between males and females.^36^ This finding suggests the possibility that blocking necroptosis could activate PPARα signaling and thereby reducing HCC in both males and females.

Chronic liver inflammation is a well-known mediator of liver fibrosis.^37^ Even though the increase in hepatic inflammation was paralleled by increased liver fibrosis in male mice in response to CD-HFD, deleting neither *Ripk3* nor *Mlkl* had any effect on liver fibrosis suggesting that necroptosis-mediated inflammation does not play a role in liver fibrosis. Contrary to our findings, deleting *Ripk3* reduced hepatic inflammation and liver fibrosis in male mice fed methionine- and choline-deficient diet or choline deficient diet.^10, 26^ However, Roychowdhury et al. (2016) reported that feeding *Ripk3^-/-^* mice a HFD exacerbated hepatic inflammation and liver fibrosis, suggesting that the response of *Ripk3* deficiency to liver fibrosis is dependent on the nature of diet, and in our study the diet contained HFD.^38^ One of the most important fibrogenic factors involved in the activation of HSCs is transforming growth factor β (TGF-β), which induces HSC growth and differentiation and the production of extracellular membrane proteins.^39^ Our data show that deleting neither *Ripk3* nor *Mlkl* had any effect on elevated TGF-β levels in response to CD-HFD, suggesting a role of TGF-β in fibrosis in both *Ripk3^-/-^* and *Mlkl^-/-^* mice. Liver fibrosis was also observed to be higher in male mice relative to females, similar to the sex-specific difference in HCC incidence, and these results are consistent with a report in humans.^40^ A role for the female sex hormone, estrogen, has been proposed for the gender differences in liver fibrosis and HCC based on the finding that the degree of liver fibrosis and HCC are higher in postmenopausal women than in premenopausal women.^17^

Our study also identified a novel cross-talk between necroptosis and other pathways that regulate tumorigenesis, e.g., apoptosis and cell senescence. Inducing apoptosis is a chemotherapeutic strategy for treating HCC, and sorafenib, the first line therapy for advanced stages of liver cancer, is a strong inducer of apoptosis.^41^ In our study, CD-HFD fed *Ripk3^-/-^* and *Mlkl^-/-^* male mice showed a disparity in apoptosis response: whereas absence of *Ripk3* elevated apoptosis, deficiency of *Mlkl* reduced apoptosis in male mice fed a CD-HFD. Afonso et al., (2021) reported increased apoptosis in the livers of *Ripk3^-/-^* mice in response to CD diet. In our study, reduction in HCC was similar in both *Ripk3^-/-^* and *Mlkl^-/-^* mice suggesting that necroptosis-mediated inhibition of HCC is independent of apoptosis.^26^ Even though cell senescence is a tumor suppressor mechanism, growing evidence suggests that senescent cells can contribute to cancer including HCC due to the secretion of senescence-associated secretory phenotype that promotes tumor progression.^42^ Our study showing that senescence markers were elevated in response to CD-HFD and this effect was reduced in both *Ripk3^-/-^* and *Mlkl^-/-^* mice suggests a possible crosstalk between necroptosis and cell senescence. *Ripk3^-/-^* mice showed a sex-specific response in cell senescence inhibition whereas *Mlkl^-/-^* mice did not show such an effect. Recently, we have shown that inhibiting necroptosis using necrostatin-1s reduced markers of cell senescence in the livers of *Sod1^-/-^* mice, a mouse model of spontaneous HCC, as well as in the livers of old WT mice, further supporting a role of necroptosis in cell senescence.^15, 43^ As cell senescence also regulates inflammation, it is possible that the observed outcome of HCC inhibition in both *Ripk3^-/-^* and *Mlkl^-/-^* mice could also be due to inhibition of both necroptosis and cellular senescence.

In summary, our data provide strong support for necroptosis playing a role in hepatic inflammation and HCC in a mouse model of NASH-induced HCC. While necroptosis mediated inflammation appears to play an important role in HCC promotion in male mice, in females necroptosis impacts HCC by mechanism other than inflammation. Our study offers insight into sex-specific differences in the progression of NASH to HCC, demonstrating the need of using both males and females for studies, especially for developing therapeutic strategies for treating HCC. Because HCC incidence is higher in postmenopausal women relative to premenopausal women, the study also highlights the importance of using estrogen-deficient female mice for studying HCC.^44^

Currently, therapeutic applications of necroptosis inhibitors for the treatment of a variety of human diseases are being tested in clinical trials.^45^ In addition, several FDA-approved anti-cancer drugs have been identified as anti-necroptotic agents.^46^ For example, sorafenib, the multi-kinase inhibitor that is clinically used to treat advanced HCC, has been shown to inhibit necroptosis and protect against inflammation and tissue injury in mouse models of TNF-induced systemic inflammatory response syndrome and renal ischemia–reperfusion injury.^47^ Thus, inhibiting necroptosis appears to be an effective strategy to prevent NASH-mediated HCC. Whether anti-necroptotic agents can also be used to treat HCC needs to be tested. In our study, we have used whole body knockout mouse models to block necroptosis, therefore, the reduction in hepatic inflammation could be due to a direct effect of blocking necroptosis in the liver tissue or a secondary effect of reduced inflammation or necroptosis in other tissues. Future studies will explore the effect of necroptosis in specific liver cell types on hepatic inflammation to identify whether inflammation in the microenvironment or at the systemic level contributes to HCC development.

## Limitations and conclusions

One of the limitations of the study is that we were unable to separate tumors from non-tumor tissue to study the effect of blocking necroptosis specifically in tumor tissues due to the small tumor size in *Ripk3^-/-^*and *Mlkl^-/-^* mice. Secondly, we focused our studies only on liver tissue and have not explored the role of adipose tissue necroptosis on inflammation, as adipose tissue is a major producer of circulating inflammatory cytokines.^48^

In addition to identifying a role of necroptosis in NASH-mediated HCC, our study also identified a sex-specific effect on inflammation and HCC. Our study offers insight into sex-specific differences in the progression of NASH to HCC, demonstrating the need of using both males and females for studies, especially for developing therapeutic strategies for treating HCC. Because HCC incidence is higher in postmenopausal women relative to premenopausal women, our study also highlights the importance of using estrogen-deficient female mice for studying HCC^.44, 49^

## Methods

### Animals and diet feeding

All procedures were approved by the Institutional Animal Care and Use Committee at the University of Oklahoma Health Sciences Center (OUHSC). *Ripk3^-/-^*mice generated by Vishva M. Dixit (Genentech, San Francisco, CA)^50^ and *Mlkl^-^*^/-^ mice generated by Warren Alexander (The Walter and Eliza Hall Institute of Medical Research, Australia)^51^ were used in these experiments. Colonies of *Ripk3^-^*^/-^ and *Ripk3^+/+^* mice and *Mlkl^-^*^/-^ and *Mlkl^+/+^* mice were generated by breeding male and female *Ripk3^+/-^* or *Mlk^+^*^/-^ mice. The mice were group housed in ventilated cages at 20 ± 2 °C, 12-h/12-h dark/light cycle. Starting at 2 months of age, male and female mice (n=7-10/group) were fed a choline deficient amino acid defined diet containing 60% fat by calories (CD-HFD) (A06071302i, Research Diets) for a period of 6 months at the OUHSC animal care facility. Normal chow diet (NC) (5053 Pico Lab, Purina Mills, Richmond, IN) or choline deficient amino acid defined diet containing 10% fat by calories (CD-LFD) (A06071302i, Research diets, New Brunswick, NJ) were used as controls. After 6 months of feeding the diets, the mice were euthanized, and liver tissue and plasma were collected for analyses. The livers were examined for the presence of visible tumors and classified based on the size and the numbers of tumors per mouse. The body weight and liver weight were also recorded for each mouse.

### Western Blotting

Western blotting was performed as described previously.^15^ Briefly, the tissues collected upon euthanasia were immediately frozen in liquid nitrogen and stored at −80 °C until further use. For western blotting, approximately 20 mg of tissues were homogenized in tissue lysis buffer [(50 mM 4-(2-hydroxyethyl)-1-piperazineethanesulfonic acid (HEPES), pH 7.6; 150 mM sodium chloride; 20 mM sodium pyrophosphate; 20 mM β-glycerophosphate; 2 mM ethylenediaminetetraacetic acid (EDTA); 1% Nonidet P-40; 10% glycerol; 2 mM phenylmethylsulfonyl fluoride; and protease inhibitor cocktail (GoldBio, St Louis, MO)]. Protein quantification was performed by Bradford assay (Bio-Rad Protein Assay Dye Reagent Concentrate, BioRad, Hercules, CA) and 40 µg of protein/well was used for performing western blotting. The samples were boiled with Laemmli buffer for 5 minutes and western blotting was done after SDS-PAGE under reducing conditions, except for the detection of MLKL oligomers. Images were taken using a Chemidoc imager (Bio-Rad) and quantified using ImageJ software (U.S. National Institutes of Health, Bethesda, MD, USA). Primary antibodies against the following proteins were used: MLKL (Cat. No. MABC604), α-smooth muscle actin (α-SMA; Cat. No. A5228) from Millipore Sigma (Burlington, MA), RIPK3 (Cat. No. NBP1-77299) from Novus Biologicals (Centennial, CO), β-tubulin (Cat. No. T5201) from Sigma-Aldrich (St. Louis, MO), Desmin (Cat. No. PA5-16705) from Invitrogen (Waltham, MA), P-SAPK/JNK (T183/Y185) (Cat. No. 4668P), SAPK/JNK (Cat. No. 9252), P-p38 MAPK (T180/Y182) (Cat. No. 9211S), p38 MAPK (Cat. No. 9212), P-p44/42 MAPK (Erk1/2) (Cat. No. 9101S), p44/42 MAPK (Erk1/2) (Cat. No. 4695S), β-catenin (Cat. No. 8480), Cyclin D1 (Cat. No. 2978), P-Akt (T308) (Cat. No. 4056), Akt2 (Cat. No. 3063S), P-S6 ribosomal protein (S235/236) (Cat. No. 4857), S6 ribosomal Protein (Cat. No. 2217), cleaved Caspase-3 (Cat. No. 9664), and Caspase-3 (Cat. No. 9662) from Cell Signaling Technology (Danvers, MA); p62 (Cat. No. MAB8028-SP), and PD-L1 (Cat. No. AF 1019) from R&D systems (Minneapolis, MN). HRP-linked anti-rabbit IgG, HRP-linked anti-mouse IgG and HRP-linked anti-rat IgG from Cell Signaling Technology (Danvers, MA) were used as secondary antibodies.

### Detection of MLKL oligomers by western blotting

MLKL oligomers were detected as described by Miyata et al. (2021).^52^ Briefly, 20 mg of liver tissue was homogenized using tissue lysis buffer. 40 µg of protein was used per sample and was prepared by heating with Laemmli buffer at 37°C for 15 minutes. The samples were loaded into 8% polyacrylamide gel without SDS, and electrophoresis was performed in running buffer containing 25 mM of Tris base and 40 mM glycine. MLKL oligomers were detected as bands above 250 kDa using MLKL antibody and HRP-linked anti-rat IgG.

### Quantitative real-time PCR (RT-PCR)

The transcript levels of different genes were analyzed as described.^15^ Briefly, 20 mg of liver tissue was used for isolating the total RNA using RNeasy kit (Qiagen, Valencia, CA). RNA was quantified using Qubit RNA BR Assay Kit (Thermo Fisher Scientific, Waltham, MA) and 1 µg equivalent of RNA was used for first-strand cDNA synthesis using High capacity cDNA reverse transcription kit [Thermo Fisher Scientific (Applied Biosystems), Waltham, MA]. This was followed by quantitative real-time PCR using Power SYBR Green PCR Master Mix [Thermo Fisher Scientific (Applied Biosystems), Waltham, MA] in a Quantstudio 12K Flex real time PCR system (Applied Biosystems). Calculations were performed by a comparative method (2^−ΔΔCt^) using β-microglobulin and hypoxanthine phosphoribosyl transferase 1 (HPRT) as controls, as described previously (Mohammed et al, 2021a). The list of primers used are given in **Table S2**.

### Characterization of immune cells by flow cytometry

Immune cell populations were analyzed by flow cytometry as described by Mohar et al (2015) with modifications.^53^ Briefly, mice were euthanized and liver was perfused through the portal vein with perfusion buffer (5 mM HEPES (Sigma-Aldrich), 0.6 mM EDTA (BioRad) in Hank’s balanced salt solution (HBSS) (Gibco, Thermo Fisher Scientific). The liver was resected out and gall bladder was removed. The lobes were cut into small pieces and incubated in digestion buffer (5 mM HEPES, 0.5 mM calcium chloride (Sigma-Aldrich), 0.075 mg/ml Liberase (Sigma-Aldrich) for 30 minutes at 37°C with gentle shaking. The reaction was stopped by adding cold blocking buffer (5 mM EDTA, 2% fetal bovine serum (FBS) (Gibco) in HBSS). The suspension was filtered through 70 µM cell strainer and centrifuged at 50 *g* for 3 minutes. The pellet containing hepatocytes was discarded and the supernatant containing the non-parenchymal cells (NPC) was collected and centrifuged at 600 *g* for 10 minutes. The pelleted NPC was then subjected to differential centrifugation using OptiPrep density gradient media (Cosmo Bio USA, Carlsbad, CA) at 1500 *g* with no brake and acceleration for 30 minutes. The interphase containing NPC was collected and pelleted at 800 *g* for 5 minutes. The cells were incubated with Live/Dead fixable violet dead cell stain [Thermo Fisher Scientific (Waltham, MA)] for the live cell gating and with the following antibodies from Biolegend (San Diego, CA): CD16/32, CD45- APC/Cy7, CD11b- PE/Cy7, Ly6C-APC, F4/80-PE, CCR2-BV 605. Data was collected using Stratedigm 4-Laser flow cytometer (San Jose, CA), and analyzed using Flow Jo software (BD Biosciences, NJ, USA) software.

### Histological analysis of liver sections

Liver tissues were fixed in 10% buffered formalin (LabChem, Zelienople, PA) and embedded in paraffin. Sections (4 µm) were stained with Hematoxylin & Eosin (H&E) as per standardized protocol at the Stephenson Cancer Center Tissue Pathology core. Images were acquired using a Nikon Ti Eclipse microscope (Nikon, Melville, NY) at 200x magnification for 3 random non overlapping fields per sample. Kupffer cell (KC) clusters were identified as groups of 10-12 cells clustered together and counted manually in a blinded fashion. The number of KC clusters per 10 mm^2^ was quantified and represented graphically.

### Immunohistochemistry

Immunohistochemistry was done as described by Lankadasari et al. (2018) with some modifications.^54^ Briefly, paraffin embedded liver sections (4 µm) were deparaffinized and hydrated by a xylene, ethanol gradient. Antigen retrieval was performed by incubating the sections with Proteinase K (20 μg/ml in Tris-EDTA buffer) (Goldbio, St.Louis, MO) for 20 minutes at 37°C. This was followed by washing with 1X PBS and permeabilization with 0.1% Triton-X 100 (Sigma-Aldrich) for 10 minutes at room temperature. Blocking was done with normal goat serum (Sigma-Aldrich) for 1 hour at room temperature. The sections were incubated with primary antibody against Glypican 3 (Thermo Fisher Scientific, Cat. No. MA5-17083) and Ki-67 (Abcam, Cat. No. ab15580, Cambridge, MA) overnight at 4°C. The next day, the sections were washed with 1X PBS and incubated with the corresponding HRP conjugated secondary antibody. Diaminobenzidine based colorimetric method was used for the detection of target proteins in the sections. Nuclei were counter stained with Mayer’s Hematoxylin (Sigma-Aldrich). Sections were dehydrated with ethanol, xylene gradient and mounted with DPX mountant (Hatfield, PA). Images were taken using a Nikon Ti Eclipse microscope (Nikon, Melville, NY) for 3 random fields per sample.

### Picrosirius red staining

Picrosirius red staining was performed with paraffin embedded liver sections (4 µM) by following a standardized protocol at the Imaging Core facility at the Oklahoma Medical Research Foundation. Images were acquired using Nikon Ti Eclipse microscope (Nikon, Melville, NY) for 3 random non-overlapping fields per sample at 200x magnification and quantified using Image J software (U.S. National Institutes of Health, Bethesda, MD).

### Enzyme Linked Immunosorbent Assay (ELISA)

Plasma separated from anticoagulated blood was used for measuring the levels of pro-inflammatory cytokines, chemokine CCL2, and HMGB1 using commercially available sandwich ELISA kits as per manufacturer’s instructions. TNF alpha mouse high sensitivity ELISA kit and IL-6 mouse high sensitivity ELISA kit were from Invitrogen (Thermo Fisher Scientific), mouse CCL2/JE/MCP-1 quantikine ELISA kit, mouse alpha-fetoprotein/AFP quantikine ELISA kit from R&D Systems, mouse HMGB1(high mobility group protein B1) ELISA Kit from Elabscience (Houston, TX). The optical density readings obtained were used to generate a four-parameter logistic curve and the concentration of the analytes in the samples were calculated by comparing to the standard curve generated.

### Alanine transaminase (ALT) colorimetric activity assay

The levels of ALT in plasma were measured using ALT colorimetric activity assay kit from Cayman Chemical Company (Ann Arbor, MI) as per manufacturer’s instructions.

### Hydroxyproline assay

The collagen content in the liver was measured by hydroxyproline content as described by Smith et al. (2016).^55^ Briefly, approximately 250 mg of liver tissue was pulverized using liquid nitrogen and digested in hydrochloric acid (Sigma-Aldrich) overnight at 110°C. From this digest, 10 ml was mixed with 150 µl of isopropanol (Sigma-Aldrich), 75 μl of Solution A (1:4 mix of 7% Chloramine T (Sigma-Aldrich (St. Louis, MO), and acetate citrate buffer [containing 57 g sodium acetate anhydrous, 33.4 g citric acid monohydrate, 435 ml 1M sodium hydroxide, 385 ml isopropanol in 1L]. The mixture was vortexed and incubated at room temperature for 10 minutes. To this, 1 ml of Solution B [3:13 mix of Ehrlich’s reagent (3 g p-dimethyl amino benzaldehyde, 10 ml absolute ethanol, 675 µl sulfuric acid) and isopropanol] was added, and incubated at 58°C for 30 minutes. The reaction was stopped by placing on ice for 10 minutes. The absorbance at 558 nm was measured in a Spectra Max M2 spectrophotometer (Molecular Devices, San Jose, CA). The absorbance values were converted into µg units using the 4-parameter standard curve generated using the standards and expressed as µg hydroxyproline/g of tissue.

### RNA sequencing and data processing

Total RNA was isolated from liver tissue using RNeasy kit (Qiagen, Valencia, CA) and library preparation was done using NEBNext Ultra II Directional RNA Library Prep Kit for Illumina (New England Biolabs, Ipswich, MA).^56^ Paired-end 150 bp read sequencing was performed, in four to six biological replicates per diet by mouse type condition, on an Illumina NextSeq 500 sequencing platform. Raw reads, in a FASTQ format, were trimmed of residual adaptor sequences using the Scythe software. Low quality bases at the beginning and end of reads were removed using Sickle, then the quality of remaining sequences was confirmed with FastQC. Trimmed quality reads were aligned to the *Mus musculus* genome reference (GRCm39/mm39) using STAR v2.4.0h.^57^ Gene-level read counts were determined using HTSeq v0.5.3p9 with the GENCODE Release M29 (GRCm39) annotation.^58^ Read-count normalization and differentially expression analyses was performed using the edgeR package from Bioconductor, following the widely used limma/voom workflow.^59^ Differentially expressed genes were organized in expression profile sets following similar phenotype variation patterns observed in tumor incidence. Functional analysis, identifying sets of genes sharing the same functionality (GO, KEGG pathways), overrepresented among the differentially expressed genes, was performed with specialized packages from Bioconductor. Ingenuity Pathway Analysis (IPA, QIAGEN, Redwood City, CA) https://www.qiagenbioinformatics.com/products/ingenuitypathway-analysis) was used further for discovery and interactive exploration of significantly impacted static and causal gene networks, pathways, disease, upstream regulators, and regulatory effects. Complete sequence data are available as GSE200923 on Gene Expression Omnibus.

### Statistical analysis

All data are represented as mean ± SEM. Two-way ANOVA with Tukey’s post hoc test was used to analyze data using GraphPad Prism (La Jolla, CA, USA). P< 0.05 is considered statistically significant. * P<0.05, ** P<0.005, *** P<0.005.

## Acknowledgements

The authors would like to thank Stephenson Cancer Center Tissue Pathology Core for performing H & E staining, the Imaging Core facility at the Oklahoma Medical Research Foundation for performing Picrosirius red staining, and the Laboratory for Molecular Biology and Cytometry Research at the University of Oklahoma Health Sciences Center for providing the facilities for the flow cytometry experiments.

## AUTHOR CONTRIBUTIONS

Sabira Mohammed, Nidheesh Thadathil, Albert L Tran, Michael Van Der Veldt, Evan H Nicklas, and Dawei Wang performed the experiments and analyzed data; Constantin Georgescu, Haritha H Nair, Phoebe Ohene-Marfo, Sangphil Oh, Wenyi Luo, Ralf Janknecht, Benjamin F Miller, Jonathan Wren, Willard M Freeman, and Sathyaseelan S Deepa analyzed the data and edited the manuscript; S.S.D provided study concept and design, and wrote the manuscript.

## Grant Support

The efforts of authors were supported by NIH grants R01AG059718 (SSD), R03 CA262044 (SSD), P30AG050911(WMF, JDW, & BFM), R01AG064951 (BFM), R56 AG067754 (BFM), R21 AR077387 (BFM), P20 GM139763 (BFM), as well as a GeroOncology Pilot Grant, (RJ & SSD) from the University of Oklahoma Health Sciences Center.

## Abbreviations

Acta2, actin alpha 2, smooth muscle; ALT, alanine transaminase; CCL2, C-C motif chemokine ligand 2; CCR2, C-C motif chemokine receptor 2; CD-LFD; choline-deficient L-amino acid defined low fat diet; CD-HFD; choline-deficient L-amino acid defined high fat diet; Col3α1, collagen 3 alpha 1; Col1α1, collagen 1 alpha 1; DAMPs, damage associated molecular patterns; Dppa2, Developmental Pluripotency Associated 2; ELISA, enzyme linked immunosorbent assay; EpCAM, epithelial cellular adhesion molecule; ERK, extracellular-signal-regulated kinase; HCC, hepatocellular carcinoma; HMGB1, high mobility group box protein 1; HSC, hepatic stellate cell; IL-1β-interleukin-1beta; IL-6, interleukin-6; JNK, c-Jun N-terminal kinase; KC, Kupffer cell; LEP, leptin; MAPK, mitogen-activated protein kinase; MLKL, pseudokinase mixed lineage kinase domain-like; mTOR, mammalian target of rapamycin; NC, Normal chow diet; NAFL, non- alcoholic fatty liver; NAFLD, nonalcoholic fatty liver disease; NASH, non-alcoholic steatohepatitis; OHP, hydroxyproline; NLRP3, NOD-, LRR- and pyrin domain-containing protein 3; PAMPs, pathogen associated molecular patterns; PD-1, programmed cell death protein; PD- L1, programmed cell death-ligand 1; PPARα, Peroxisome Proliferator Activated Receptor Alpha; PRR, pattern recognition receptor; PSR, picrosirius red; RIPK1, receptor-interacting protein kinase 1; RIPK3, receptor-interacting protein kinase 3; RNA-seq, RNA sequencing; RT-PCR, real time polymerase chain reaction; α-SMA, alpha smooth muscle actin; Sod1, Cu/Zn superoxide dismutase; STAT5, Signal transducer and activator of transcription 5; TGF-β, transforming growth factor β; TLR4, toll-like receptor 4; TNFα, tumor necrosis factor alpha; Tonsl, Tonsoku Like, DNA Repair Protein; WT, wild-type.

## Disclosures

No potential conflicts.

## Transcript Profiling

Complete RNA-sequencing data are available as GSE200923 on Gene Expression Omnibus.

## Data Transparency

Analytic methods, and study materials will be made available to other researchers on request.

## Supplementary figure legends

**Figure S1.**
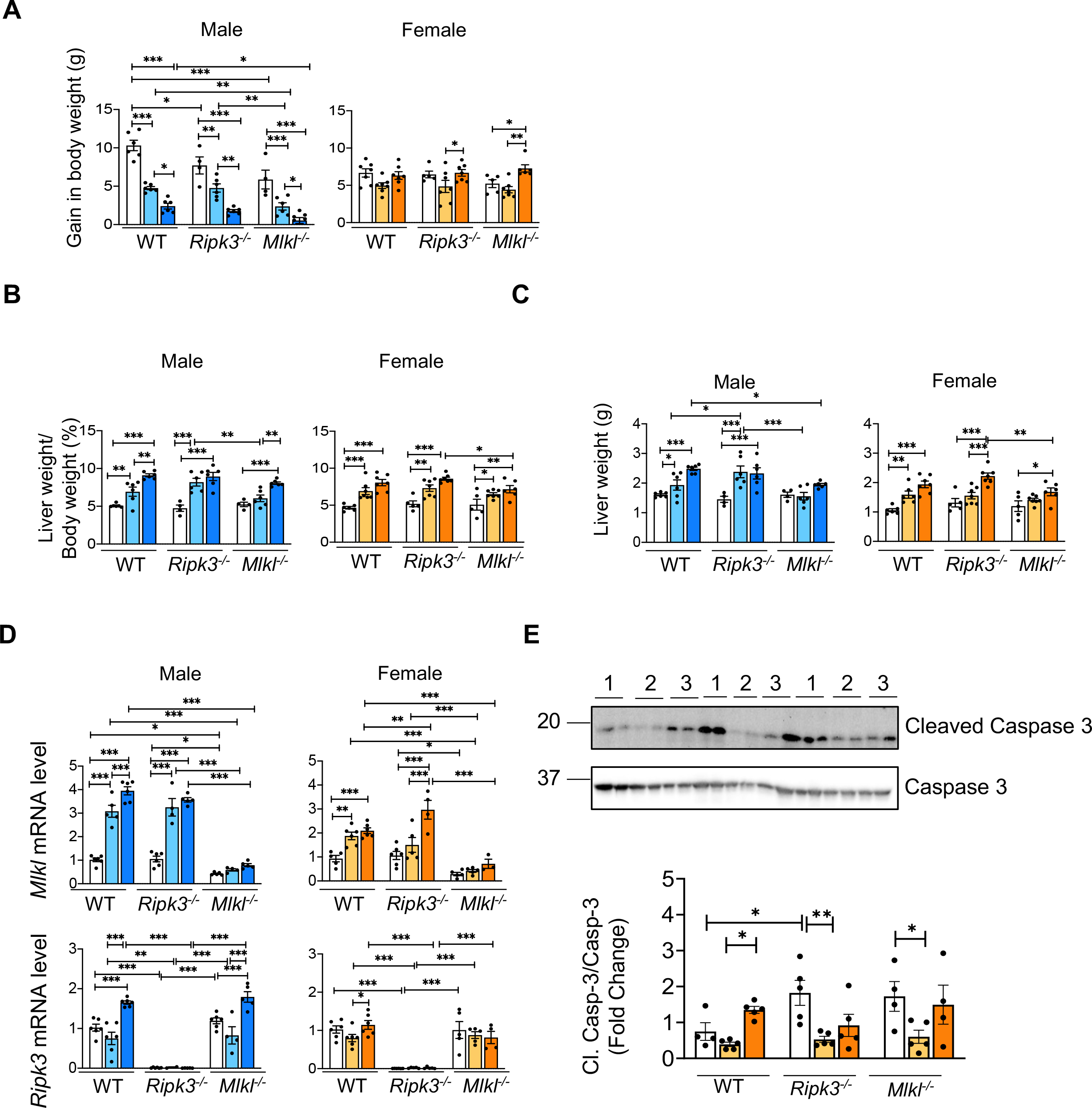
**(A)** Graphical representation of comparison of body weight gain by WT, *Ripk3^-/-^*, and *Mlkl^-/-^* male (left) and female (right) mice fed NC or CD-LFD or CD-HFD. Males: NC (white bars), CD-LFD (light blue bars) or CD-HFD (dark blue bars). Females: NC (white bars), CD-LFD (light orange bars) or CD-HFD (dark orange bars). Graphical representation of liver weight normalized to body weight **(B)** and liver weight **(C)** for WT, *Ripk3^-/-^*, and *Mlkl^-/-^* male (left) and female (right) mice fed NC or CD-LFD or CD-HFD. **(D)** Transcript levels of *Mlkl* (top panel) and *Ripk3* (bottom panel) in WT, *Ripk3^-/-^*, and *Mlkl^-/-^* male (left) and female (right) mice fed NC or CD-LFD or CD-HFD. **(E)** *Top panel:* Immunoblots of liver tissue extracts for cleaved Caspase-3 and Caspase-3 from WT, *Ripk3^-/-^*, and *Mlkl^-/-^* female mice fed NC (1) or CD-LFD (2) or CD-HFD (3). *Bottom panel:* Graphical representation of quantified blots for cleaved Caspase-3 normalized to Caspase-3. Data are represented as mean±SEM, n=4-6 per group, *p<0.05, **p<0.01, ***p<0.001.

**Figure S2.**
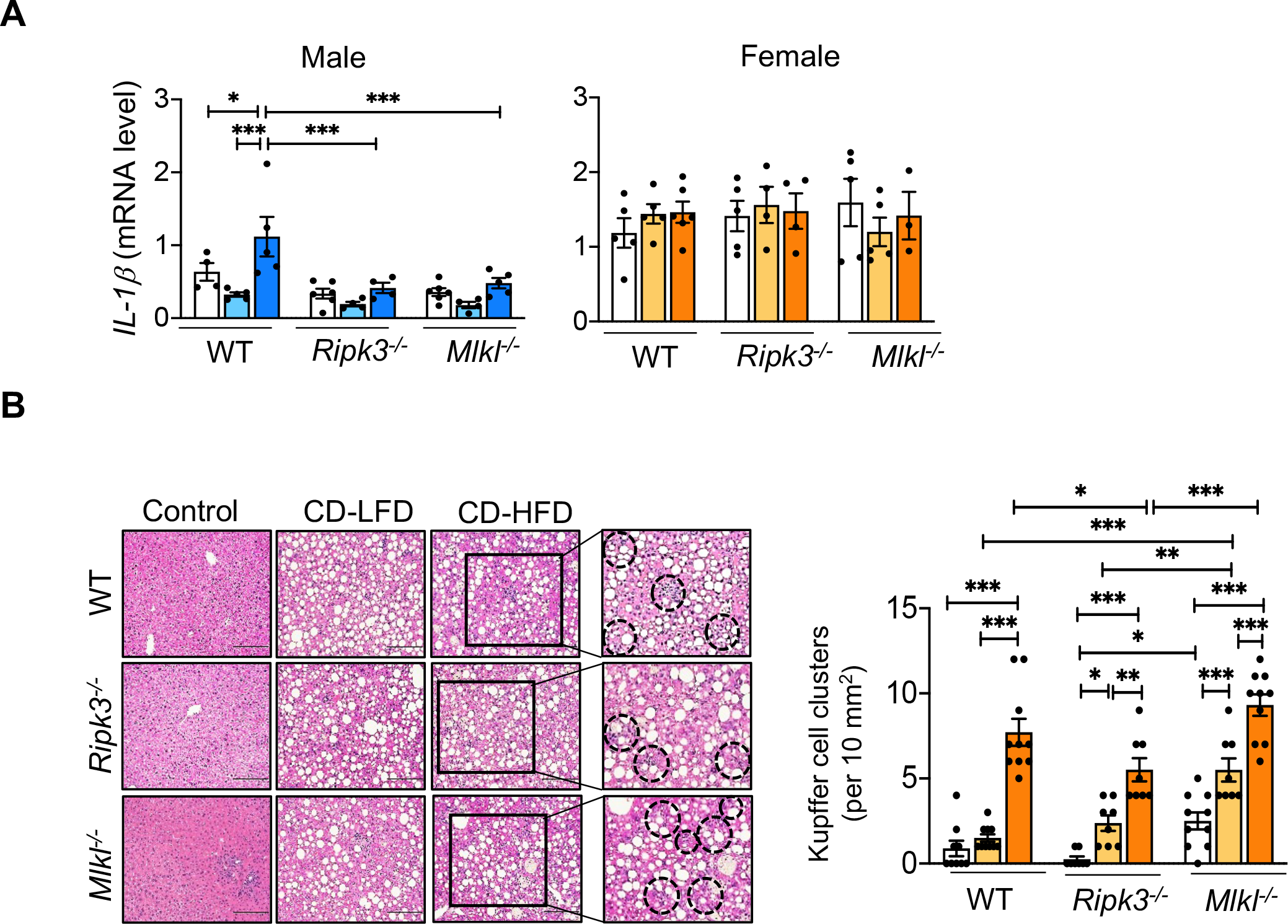
**(A)** Transcript levels of *IL-1β* in WT, *Ripk3^-/-^*, and *Mlkl^-/-^* male (left) and female (right) mice fed NC or CD-LFD or CD-HFD. **(B)** *Left panel:* Images of H&E stained sections from WT, *Ripk3^-/-^*, and *Mlkl^-/-^* female mice fed NC or CD-LFD or CD-HFD. Black dotted circles represent KC clusters. Scale bar: 50 μM. *Right panel:* Graphical representation of the number of KC clusters in each group. Data are represented as mean±SEM, n=4-6 per group, *p<0.05, **p<0.01, ***p<0.001. Males: NC (white bars), CD-LFD (light blue bars) or CD-HFD (dark blue bars). Females: NC (white bars), CD-LFD (light orange bars) or CD-HFD (dark orange bars).

**Figure S3.**
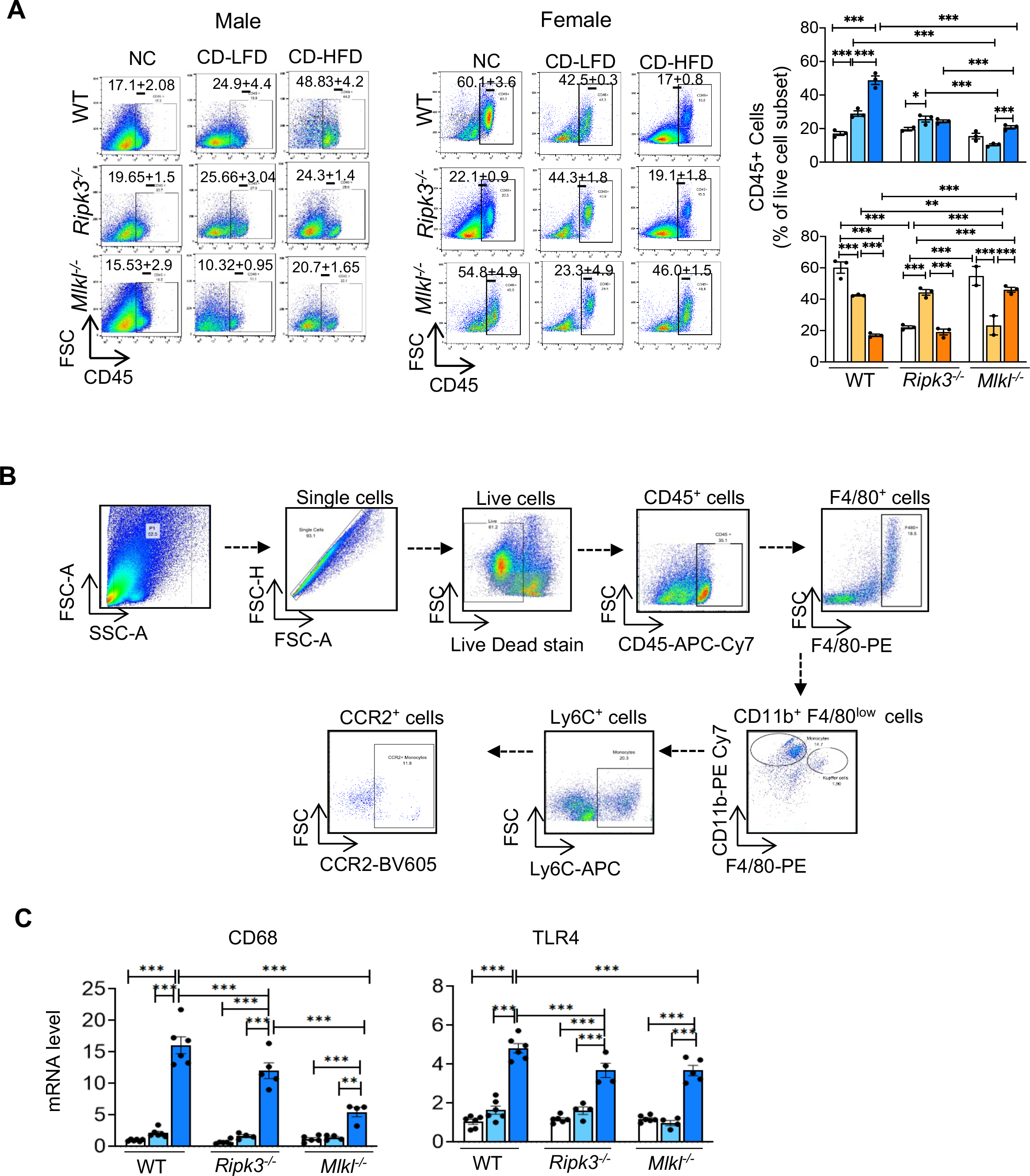
**(A)** *Left panel:* Flow cytometric analysis of percentage of CD45^+ve^ cell population in the livers of WT, *Ripk3^-/-^*, and *Mlkl^-/-^* male (left) and female (right) mice fed NC or CD-LFD or CD-HFD. *Right panel:* Graphical representation of the percentage population of gated CD45^+ve^ cells in male mice (top) and female (bottom) mice. **(B)** Gating strategy that was followed for flow cytometry analysis. **(C)** Transcript levels of *CD68* (left) and *TLR4* (right) in the livers of WT, *Ripk3^-/-^*, and *Mlkl^-/-^*male mice fed NC or CD-LFD or CD-HFD. Data are represented as mean±SEM, n=4-6 per group, *p<0.05, **p<0.01, ***p<0.001. Males: NC (white bars), CD-LFD (light blue bars) or CD-HFD (dark blue bars). Females: NC (white bars), CD-LFD (light orange bars) or CD-HFD (dark orange bars).

**Figure S4.**
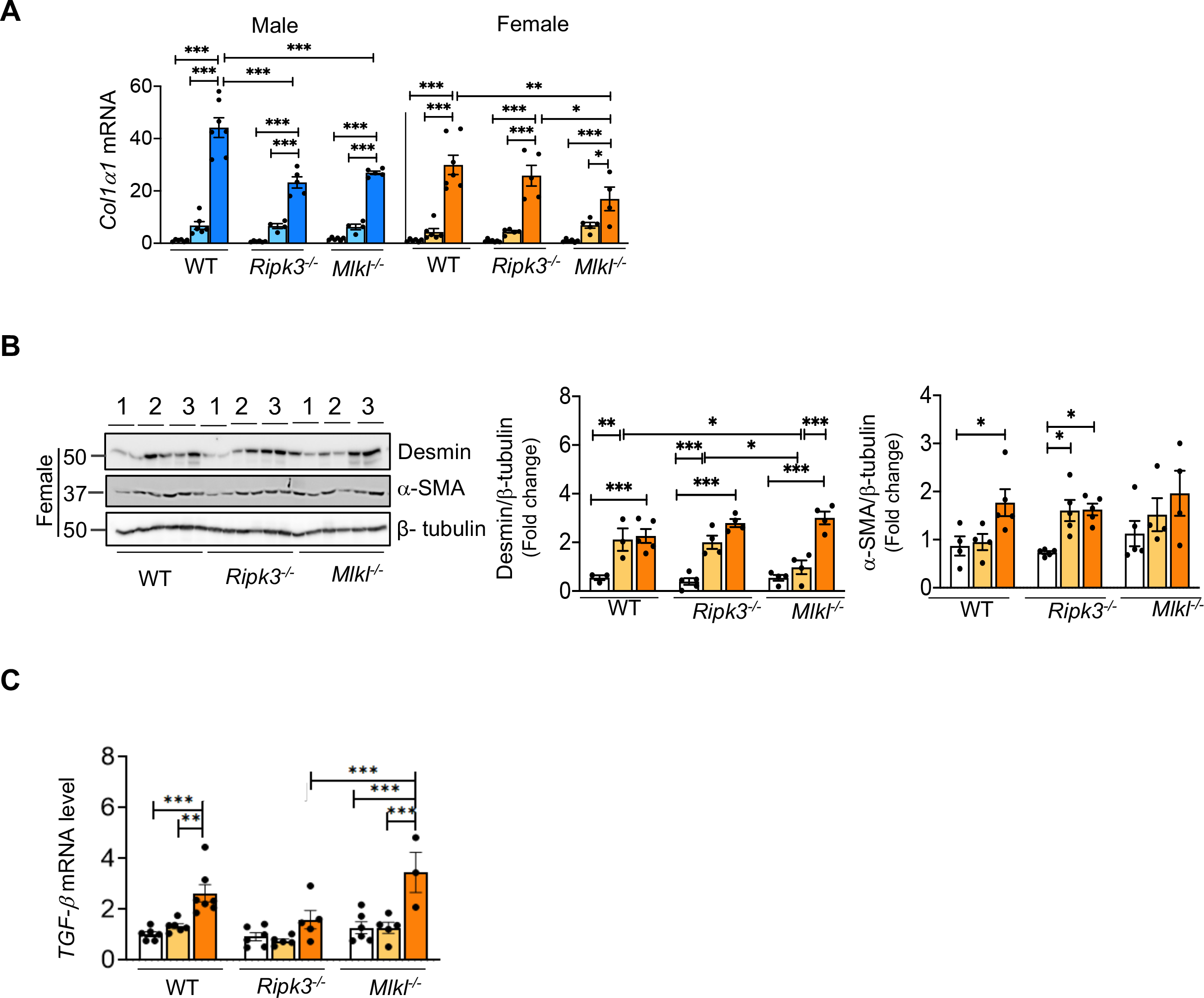
**(A)** Transcript levels of *Col1a1* in the livers of WT, *Ripk3^-/-^*, and *Mlkl^-/-^* male (left) and female (right) mice fed NC or CD-LFD or CD-HFD. **(B)** *Left panel*: Immunoblots of liver tissue extracts from WT, *Ripk3^-/-^*, and *Mlkl^-/-^* female mice fed NC or CD-LFD or CD-HFD for desmin,α- SMA and β-tubulin. *Right panel:* Graphical representation of quantified blots normalized to β- tubulin. **(C)** Transcript levels of *TGF-β* in the livers of WT, *Ripk3^-/-^*, and *Mlkl^-/-^* female mice fed NC or CD-LFD or CD-HFD. Data are represented as mean±SEM, n=4-6 per group, *p<0.05, **p<0.01, ***p<0.001. Males: NC (white bars), CD-LFD (light blue bars) or CD-HFD (dark blue bars). Females: NC (white bars), CD-LFD (light orange bars) or CD-HFD (dark orange bars).

**Figure S5.**
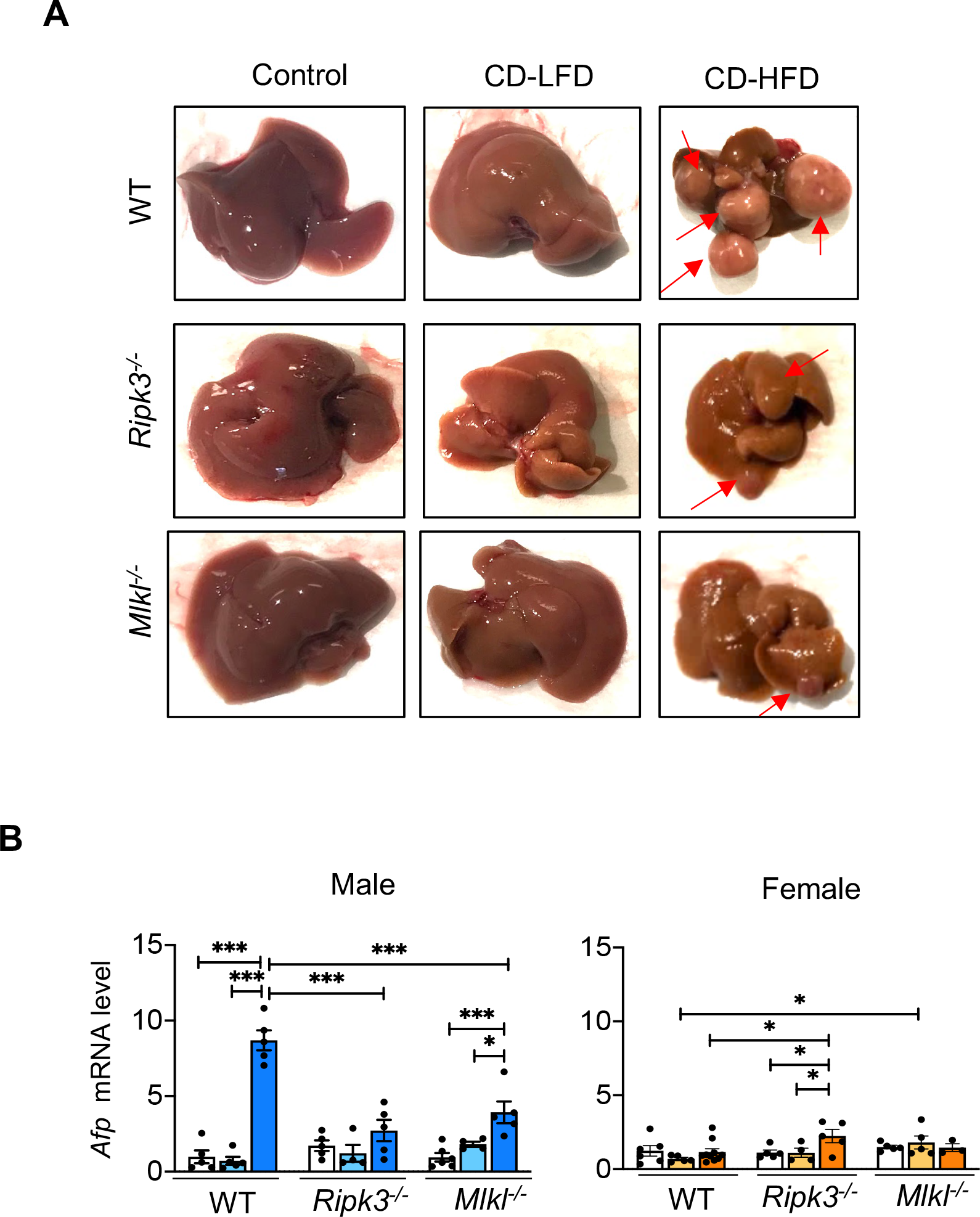
**(A)** Representative images of liver nodules in WT, *Ripk3^-/-^*, and *Mlkl^-/-^* male mice fed NC or CD-LFD or CD-HFD. **(B)** Transcript levels of *AFP* normalized to *β-microglobulin* in WT, *Ripk3^-/-^*, and *Mlkl^-/-^* male (left) and female (right) mice fed NC or CD-LFD and CD-HFD. Data are represented as mean±SEM, n=5-7 per group, *p<0.05, **p<0.01, ***p<0.001. Males: NC (white bars), CD-LFD (light blue bars) or CD-HFD (dark blue bars). Females: NC (white bars), CD-LFD (light orange bars) or CD-HFD (dark orange bars).

**Figure S6.**
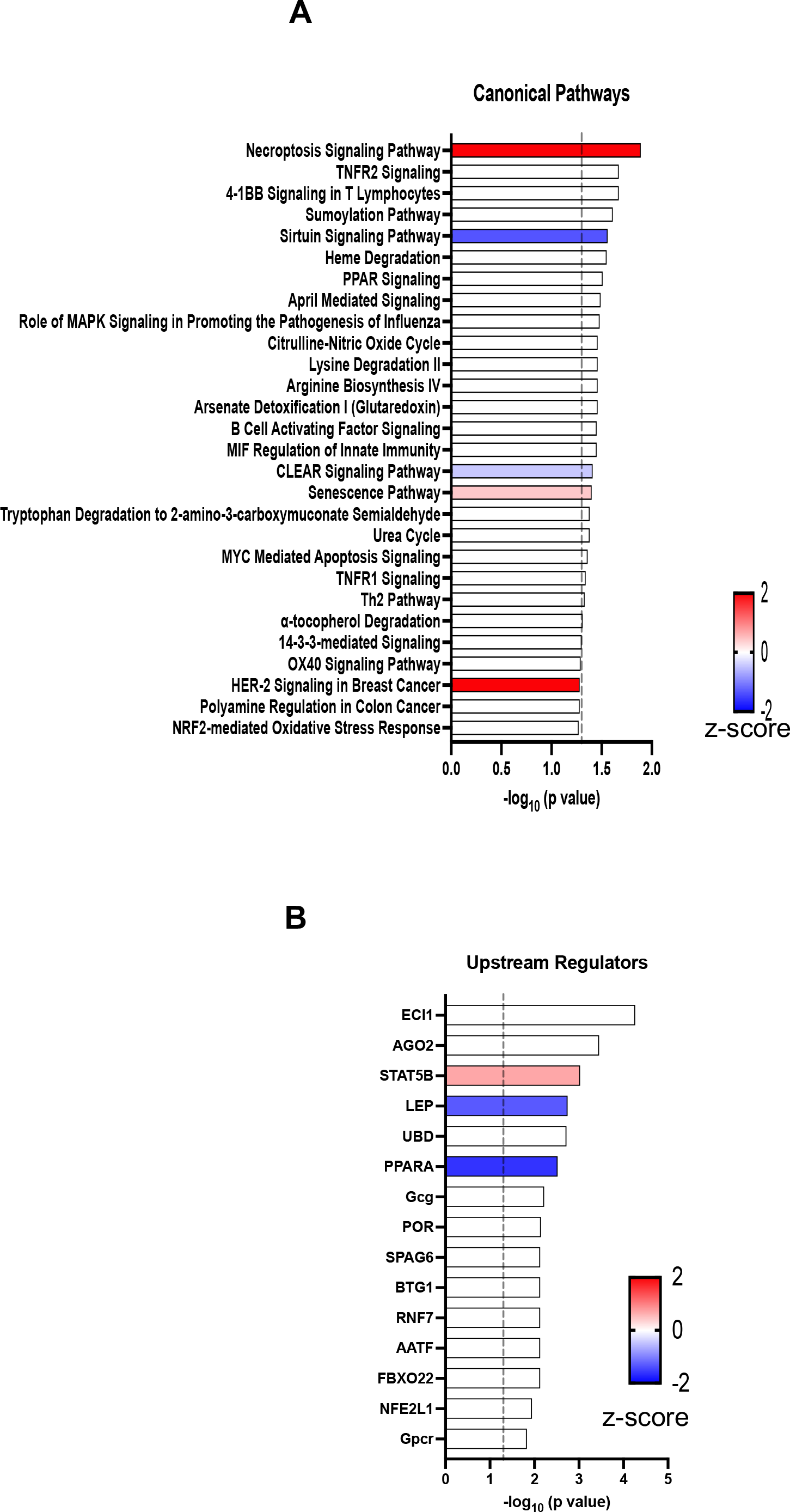
**(A)** The ingenuity pathway analysis for canonical pathways altered in the livers of WT, *Ripk3^-/-^*, and *Mlkl^-/-^* male mice fed by NC, CD-LFD or CD-HFD. **(B)** The ingenuity pathway analysis for upstream regulatory molecules and pathways affected in WT, *Ripk3^-/-^*, and *Mlkl^-/-^* male mice fed by NC or CD-LFD or CD-HFD.

**Figure S7.**
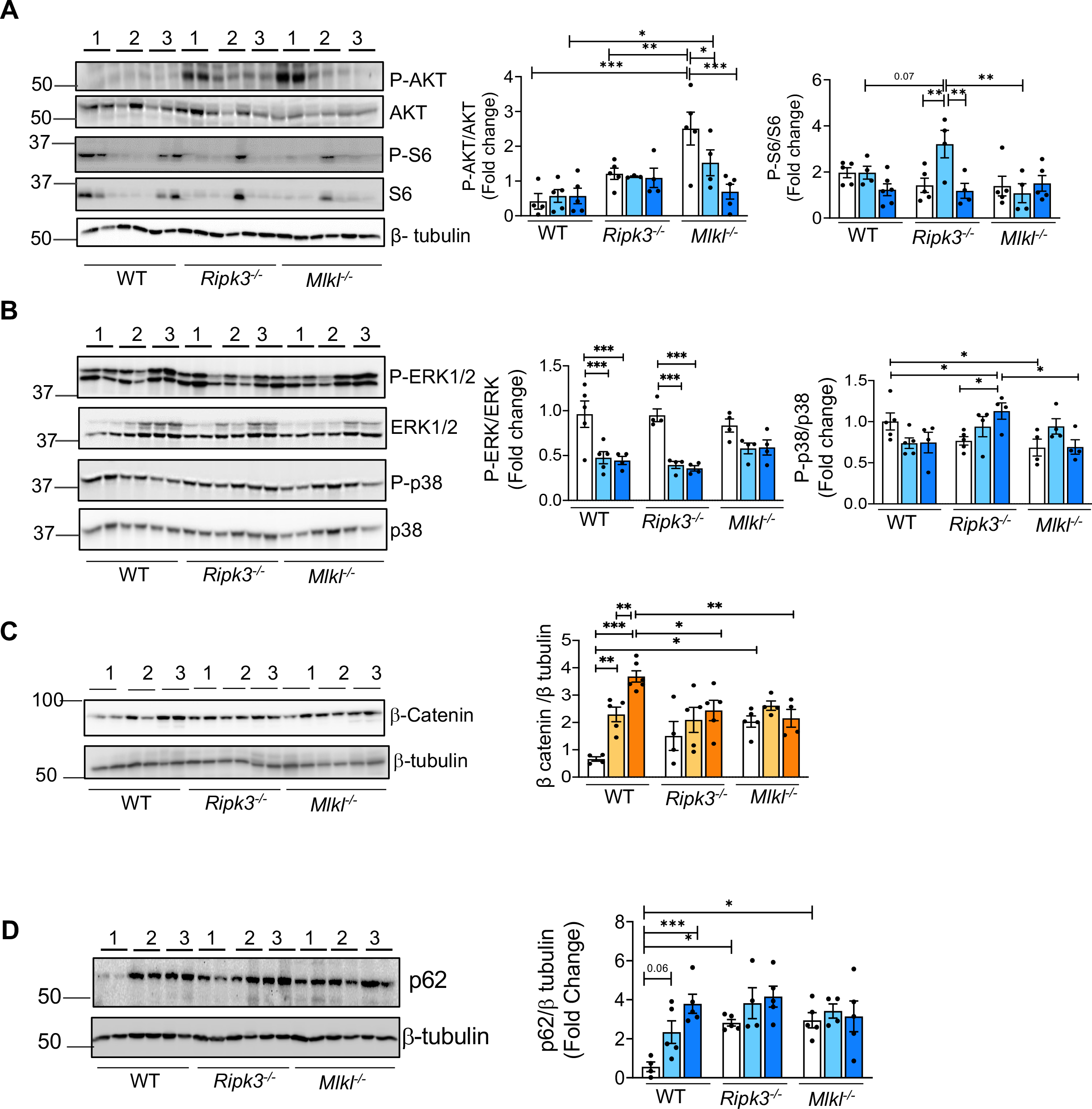
*Left panel:* Immunoblots of liver tissue extracts from WT, *Ripk3^-/-^*, and *Mlkl^-/-^* male mice fed NC or CD-LFD or CD-HFD for p-AKT, AKT, p-S6, S6, and β-tubulin **(A)**; p-ERK1/2, ERK1/2, p-p38, p38, and β-tubulin **(B)**. *Right panel:* Graphical representation of fold change of phosphorylated proteins to respective unphosphorylated proteins. **(C)** *Left panel:* Immunoblots of liver tissue extracts for β-catenin and β-tubulin from WT, *Ripk3^-/-^*, and *Mlkl^-/-^*female mice fed NC or CD-LFD or CD-HFD (left panel). *Right panel:* Graphical representation of β-catenin normalized to β-tubulin. **(D)** *Left panel:* Immunoblots of liver tissue extracts for p62 and β-tubulin from WT, *Ripk3^-/-^*, and *Mlkl^-/-^* male mice fed NC or CD-LFD or CD-HFD. Right panel: Graphical representation of quantified blots normalized to β-tubulin. Data are represented as mean±SEM, n=4-6 per group, *p<0.05, **p<0.01, ***p<0.001. Males: NC (white bars), CD-LFD (light blue bars) or CD-HFD (dark blue bars). Females: NC (white bars), CD-LFD (light orange bars) or CD-HFD (dark orange bars).

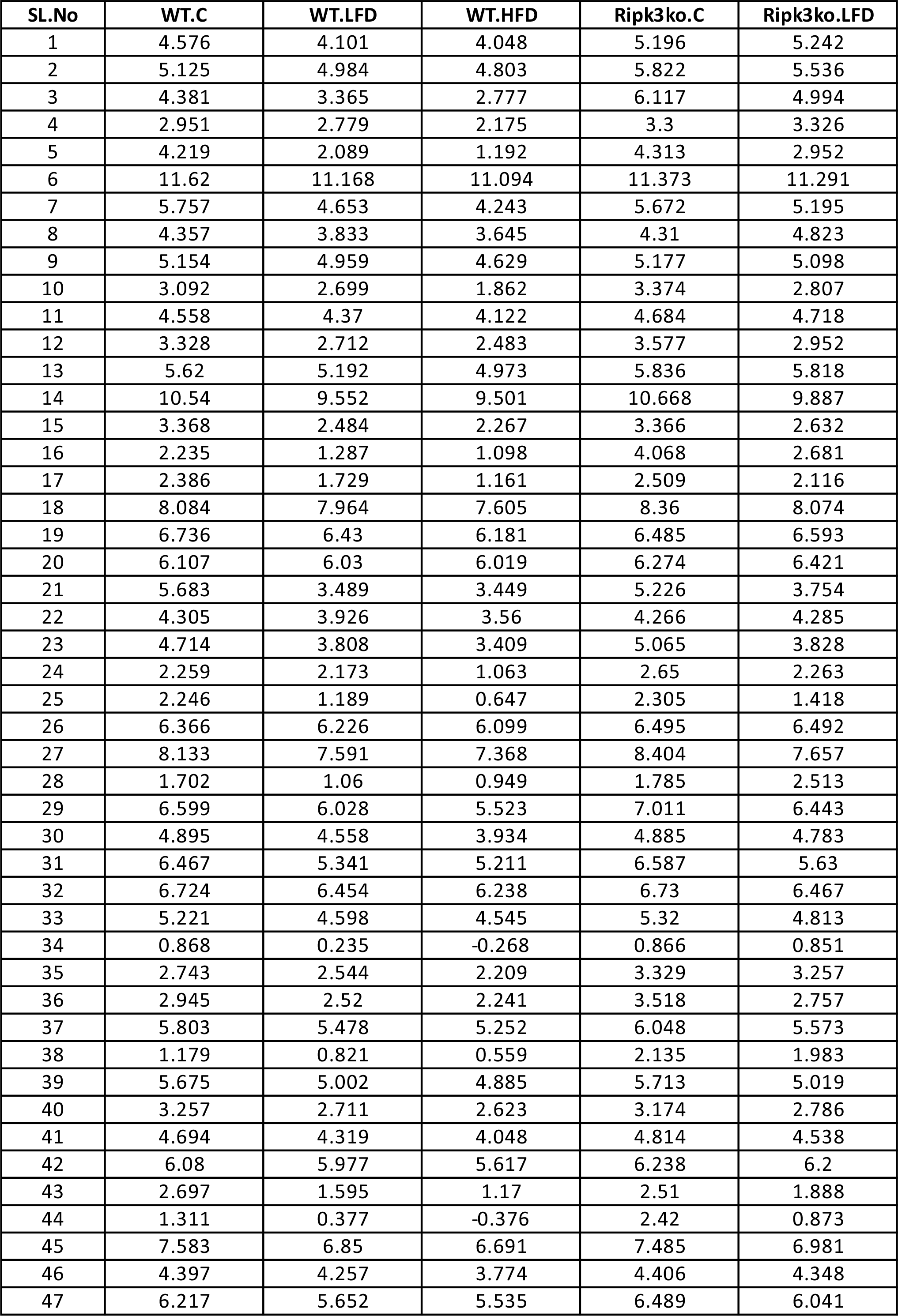
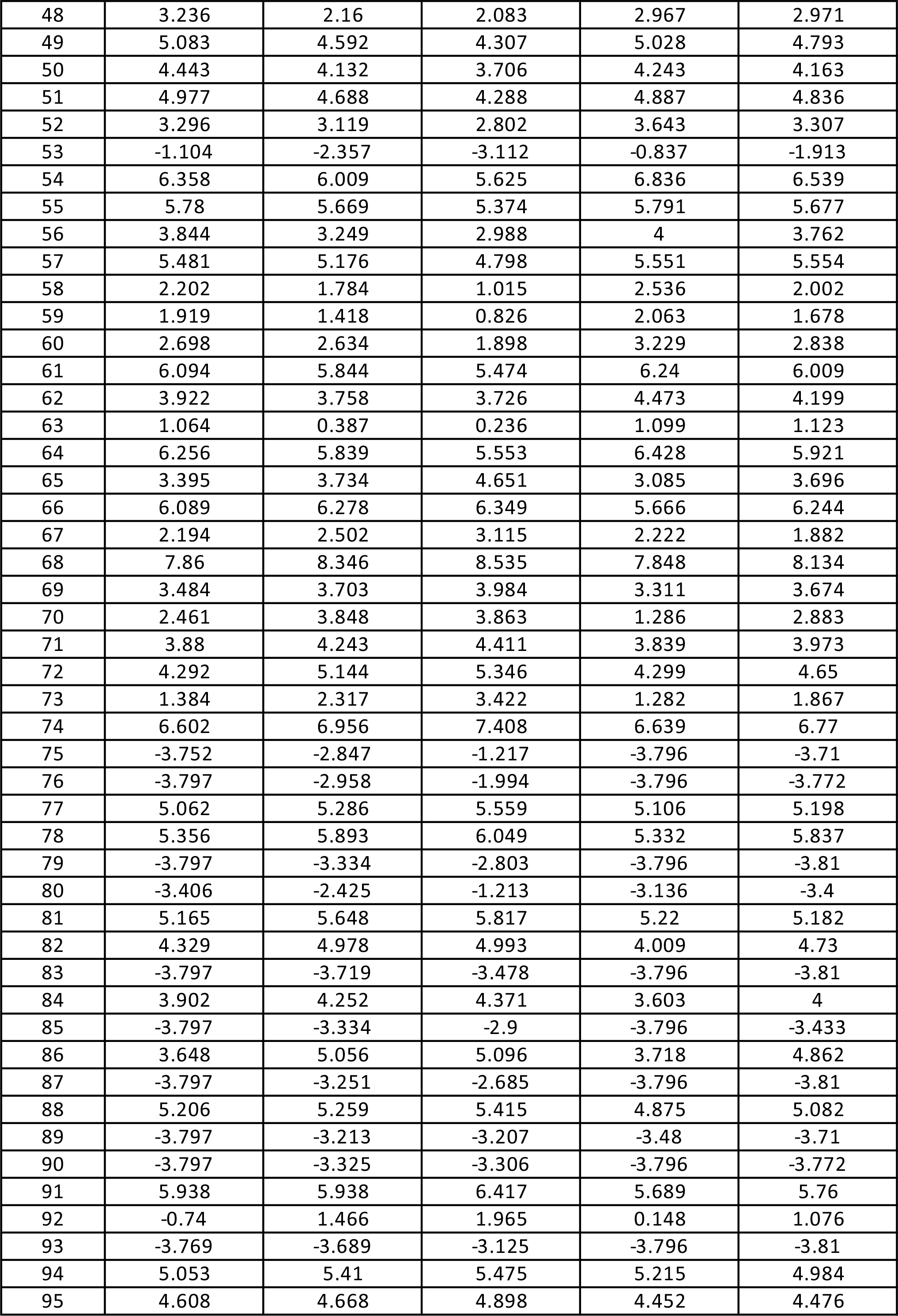
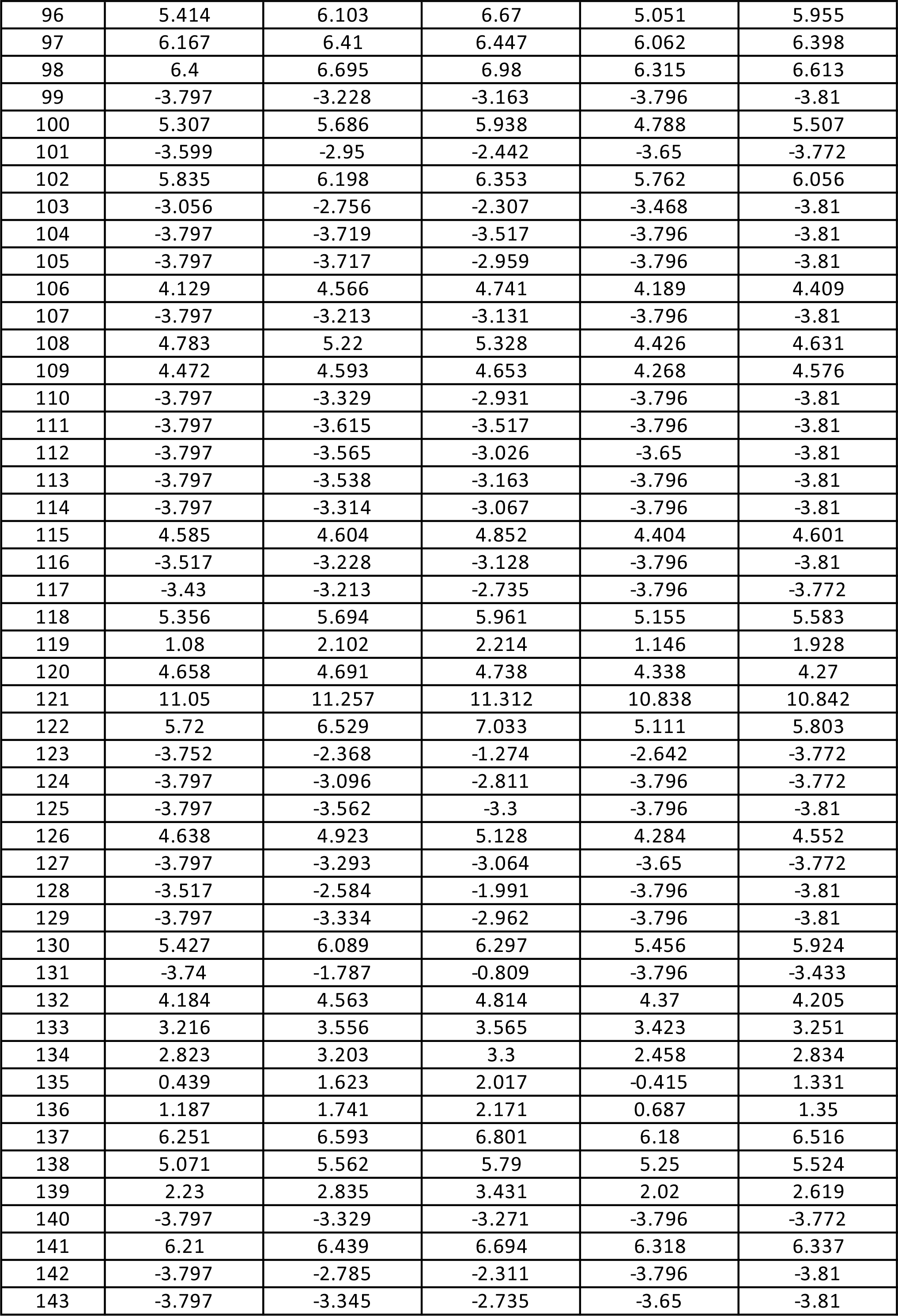
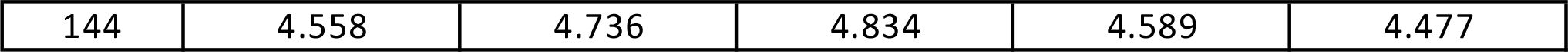
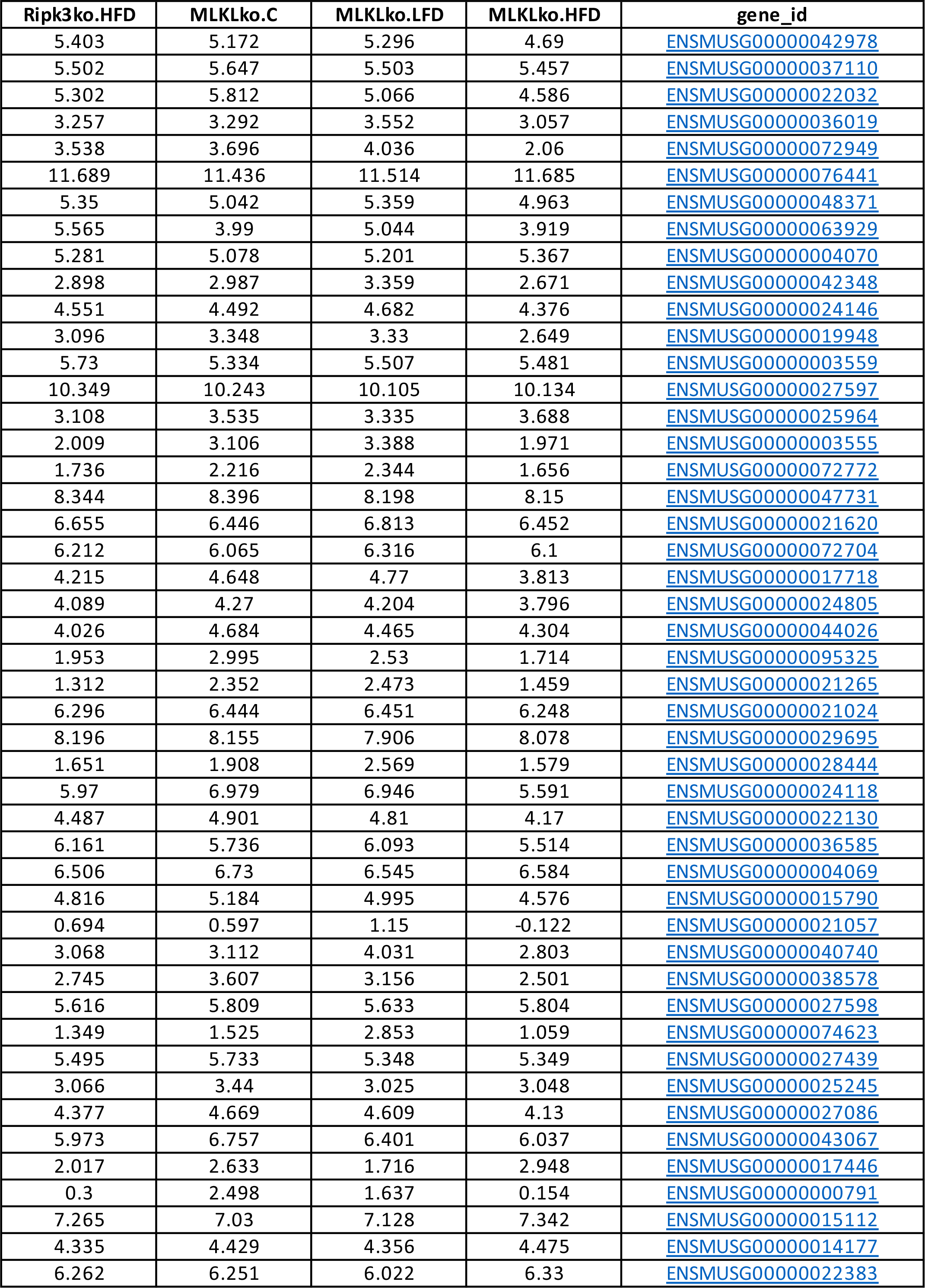
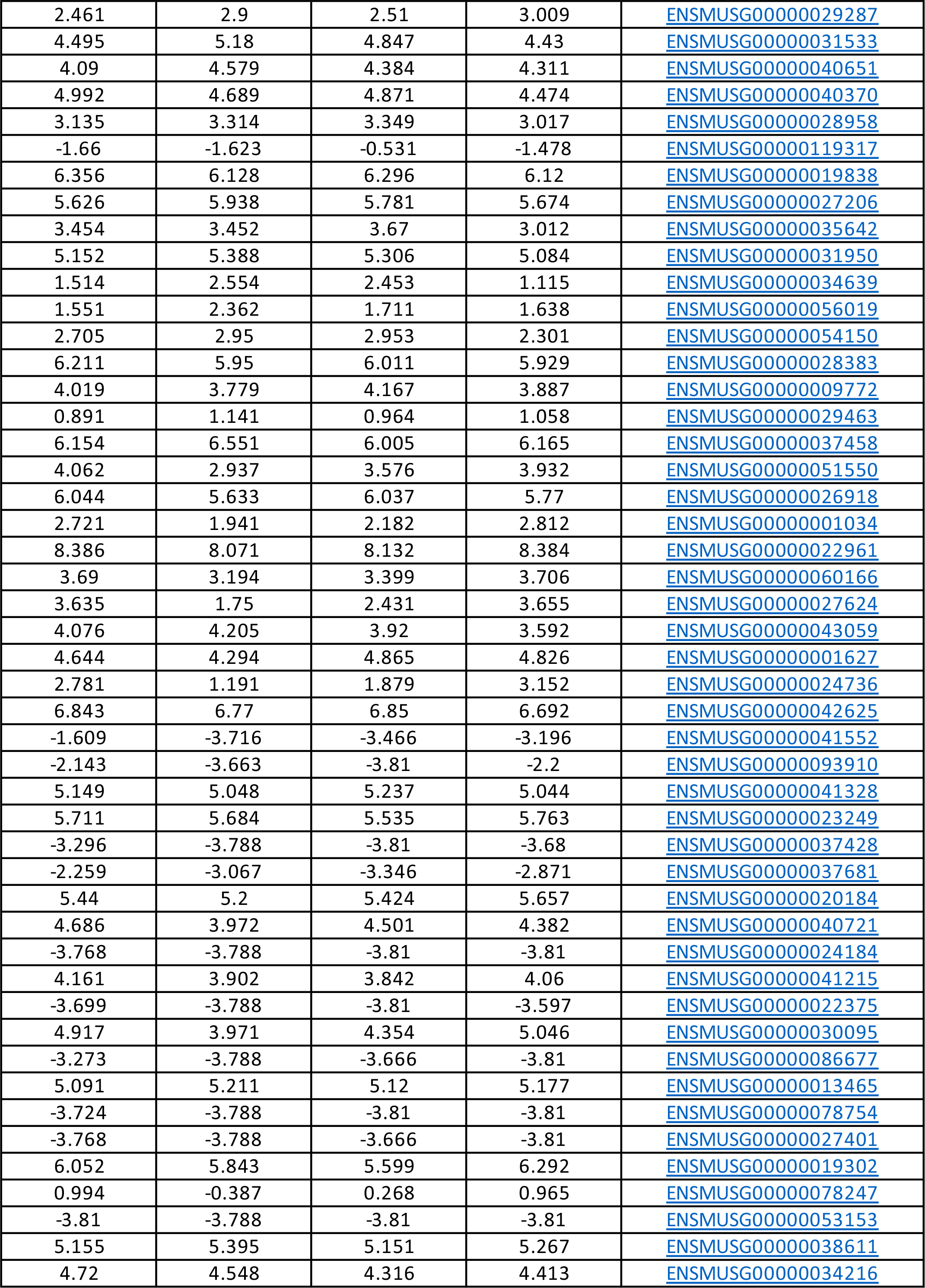
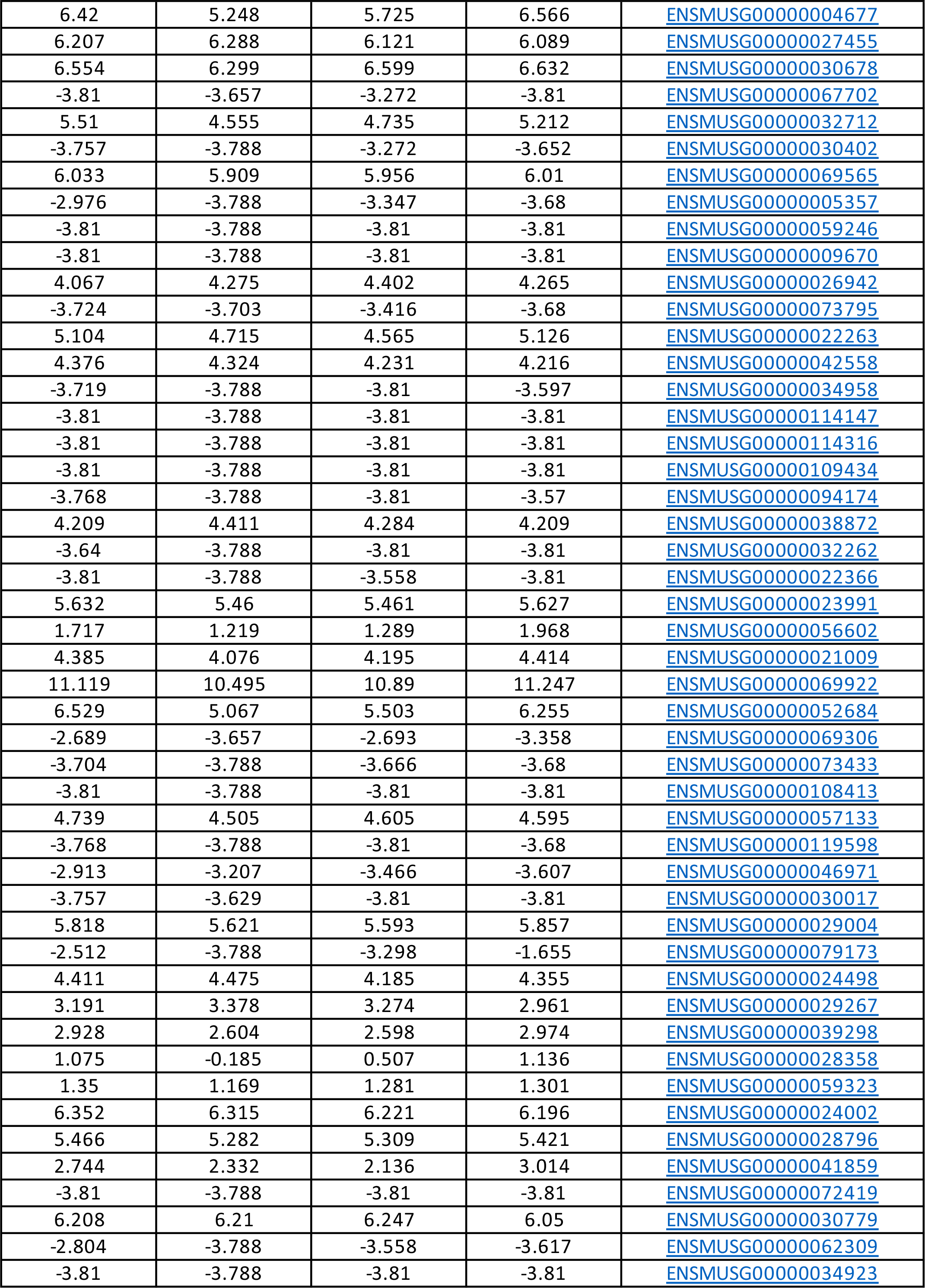
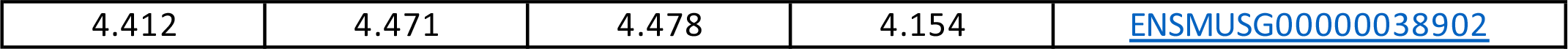
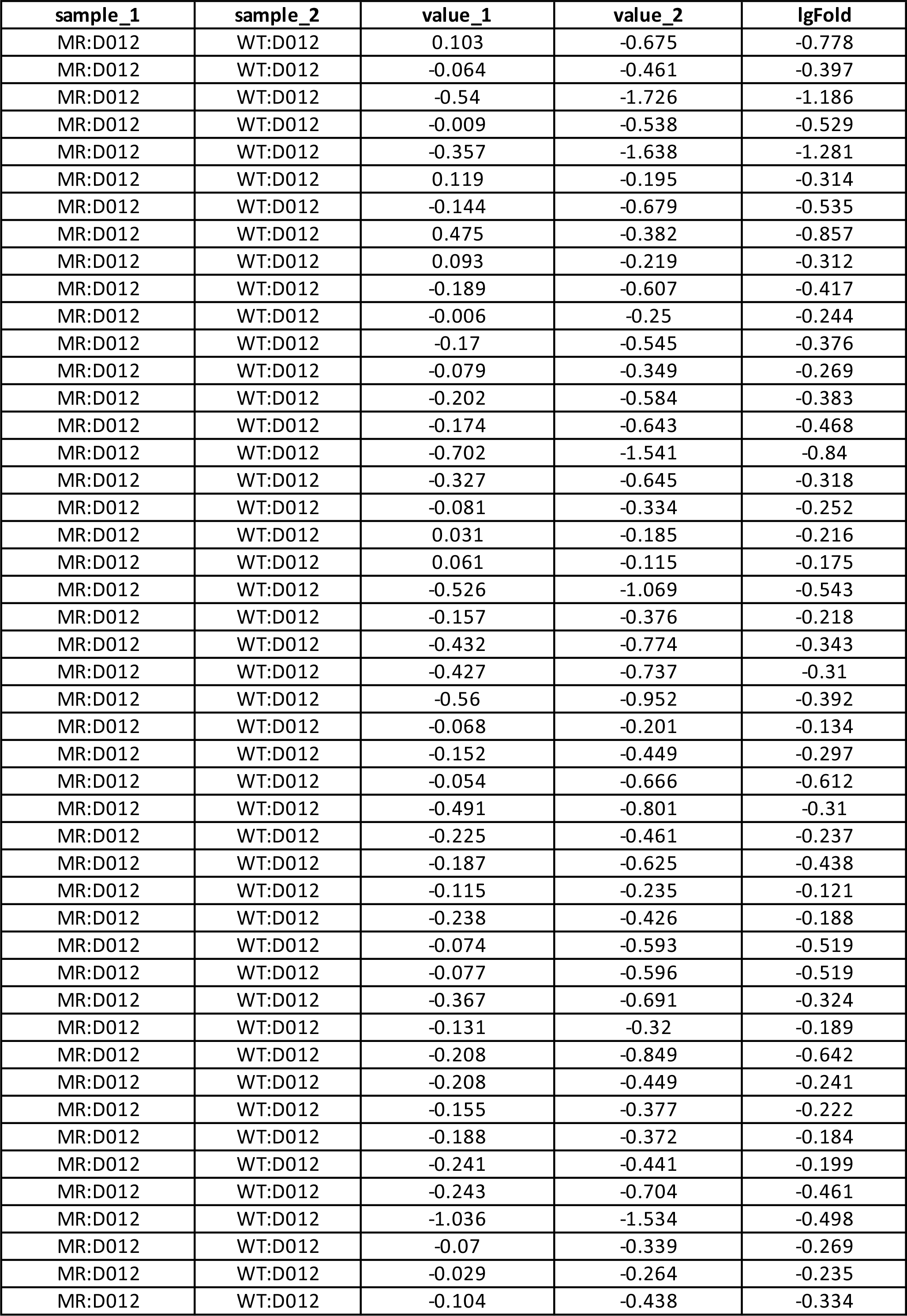
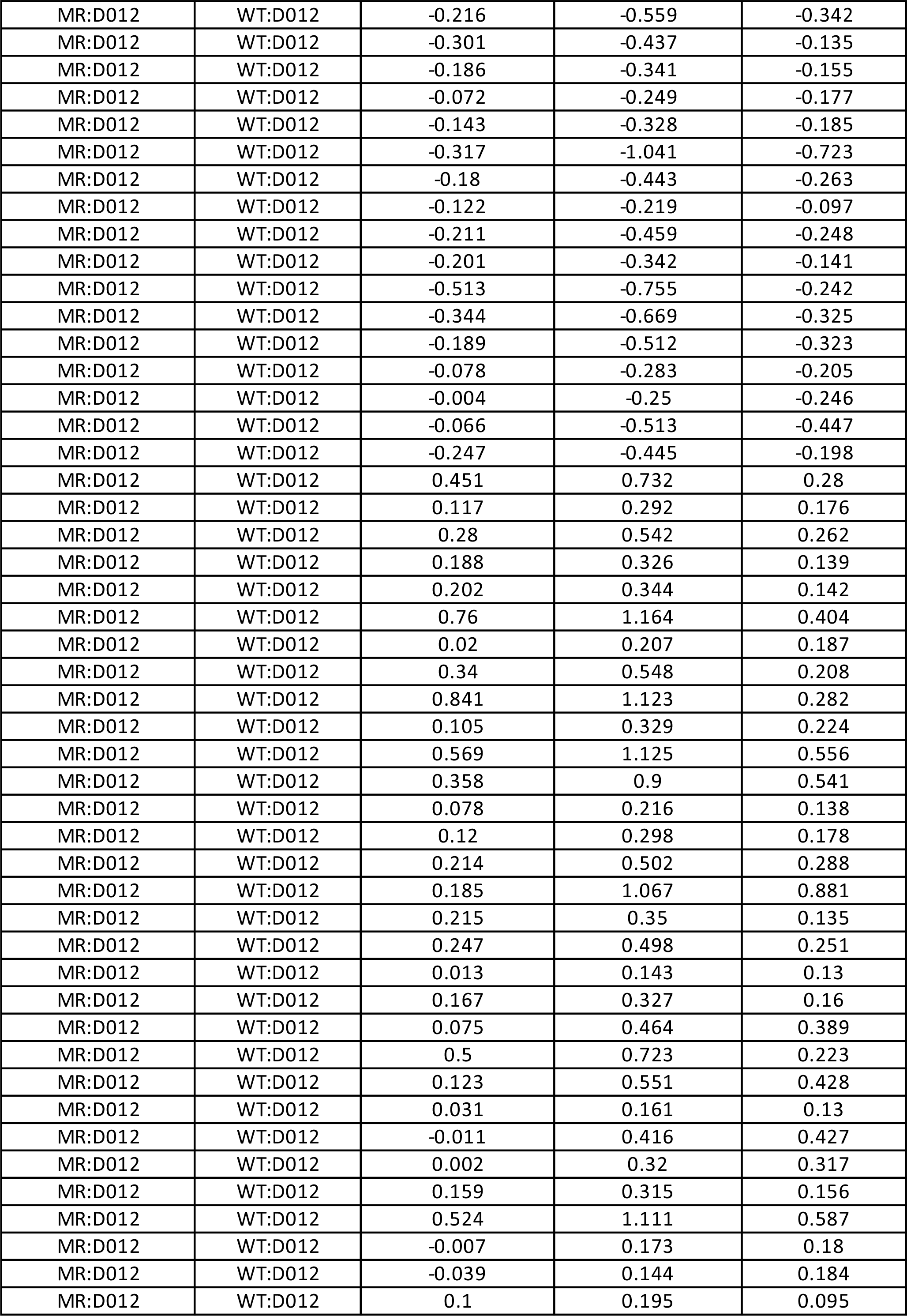
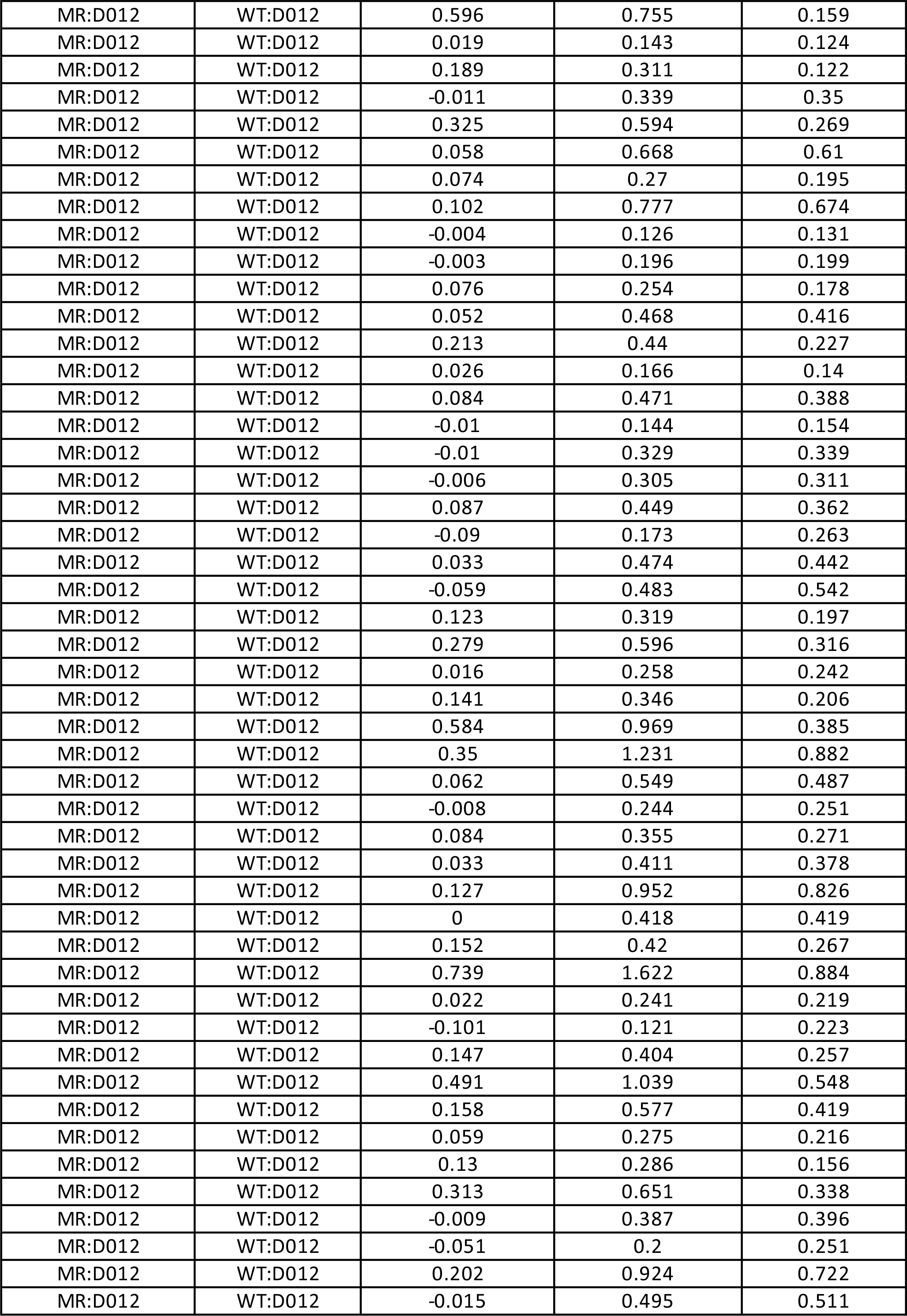
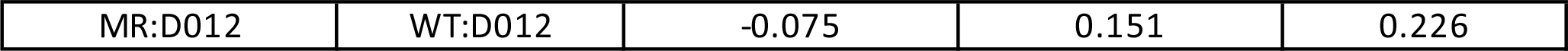
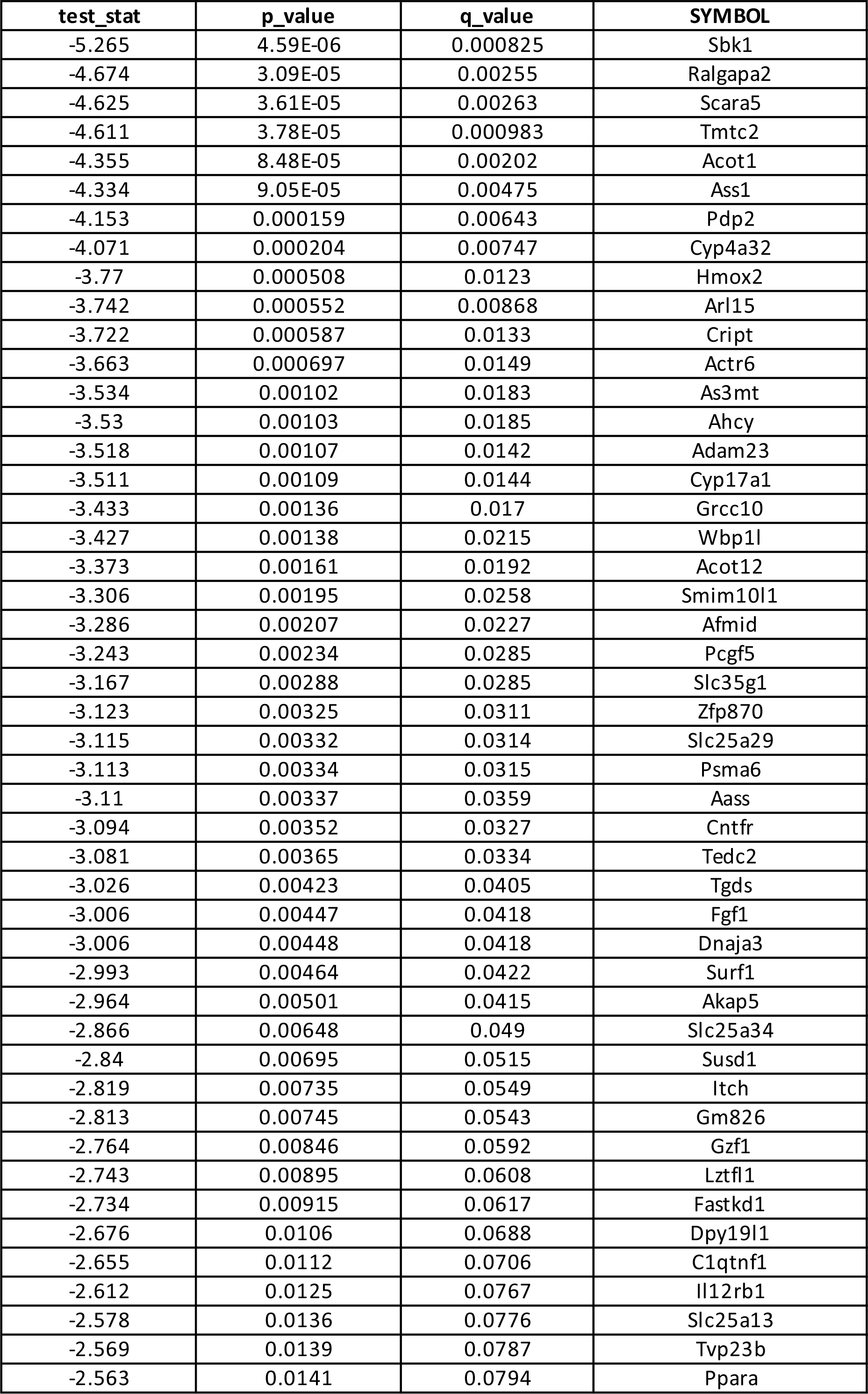
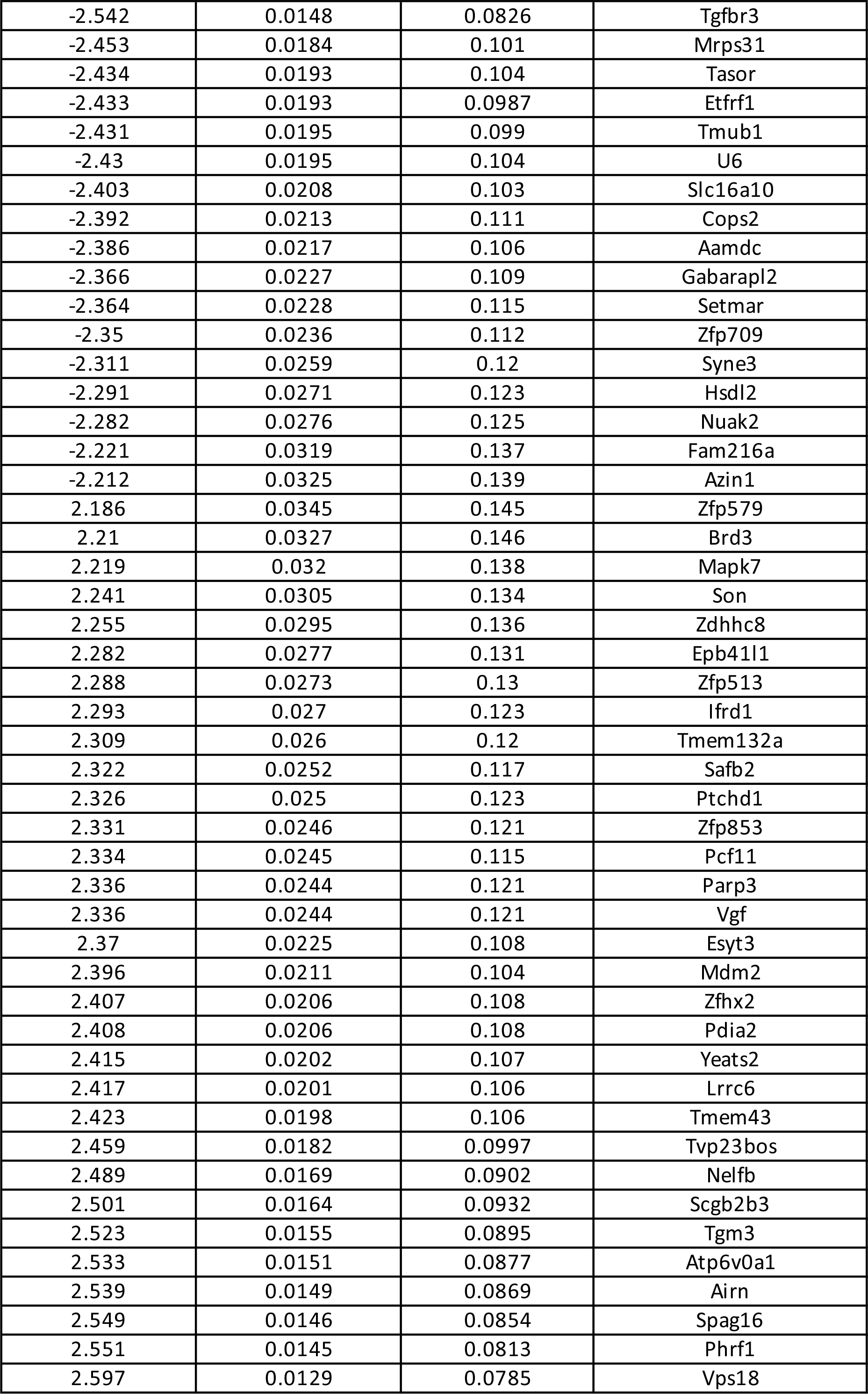
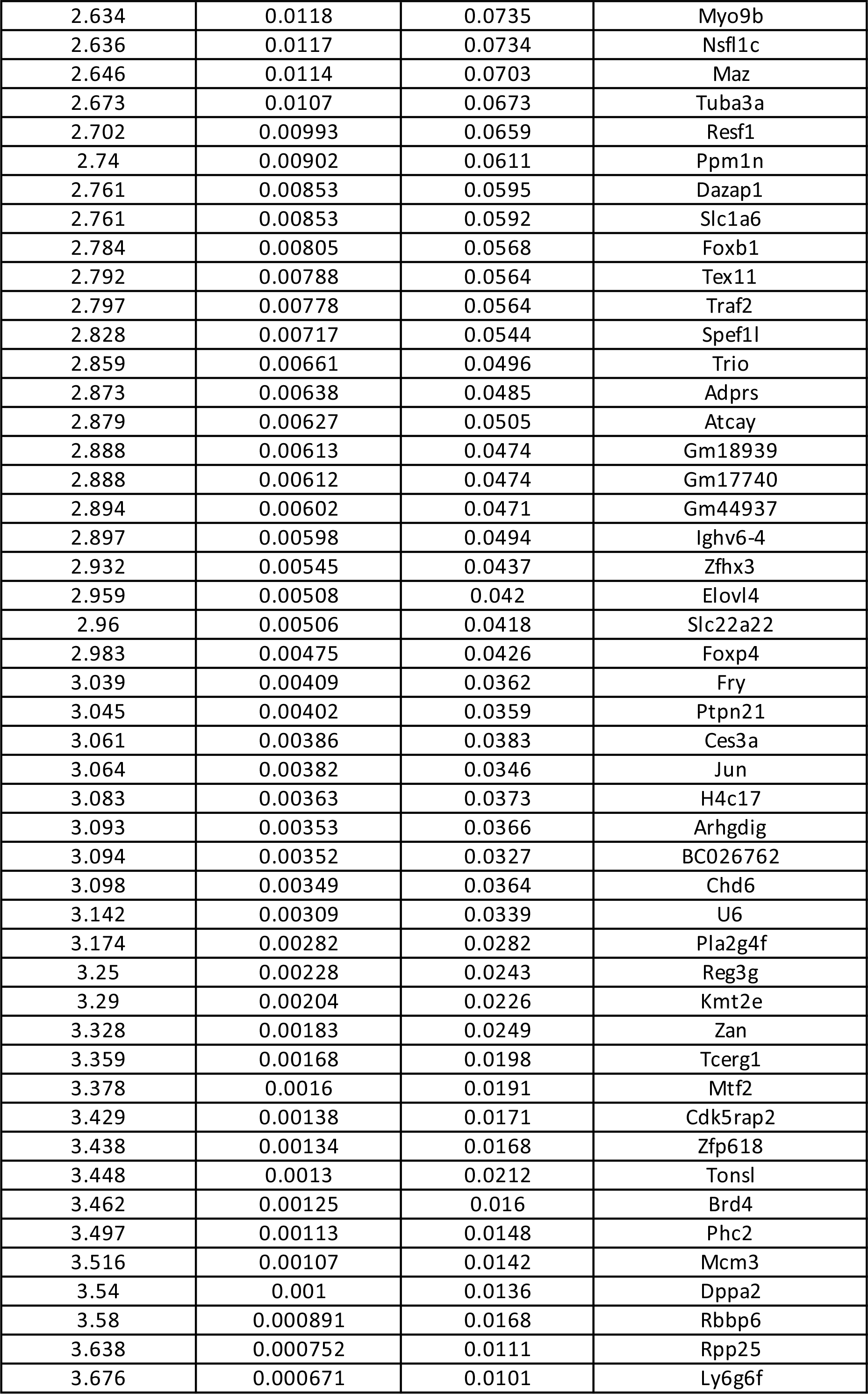
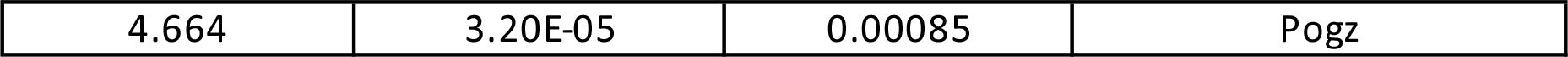
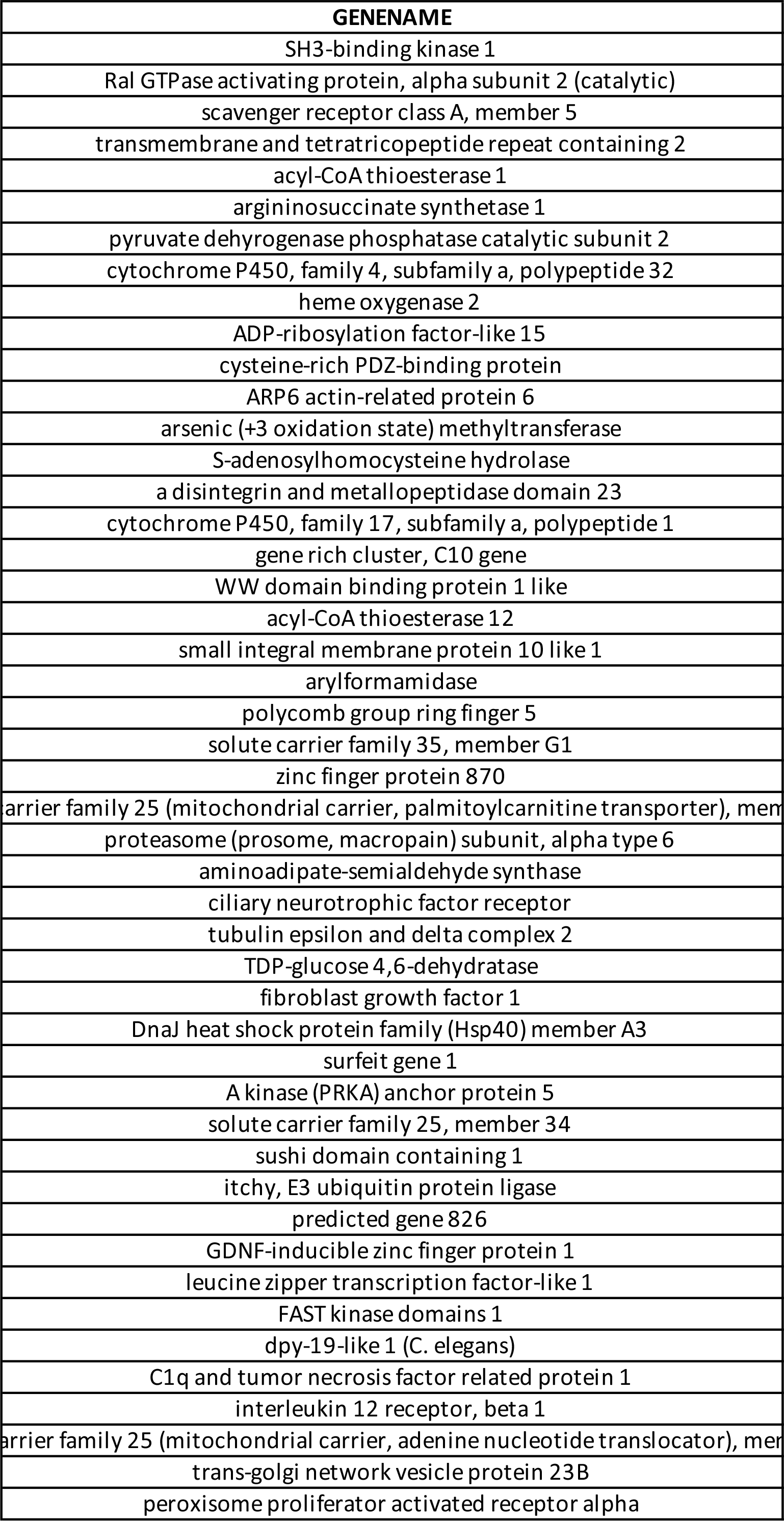
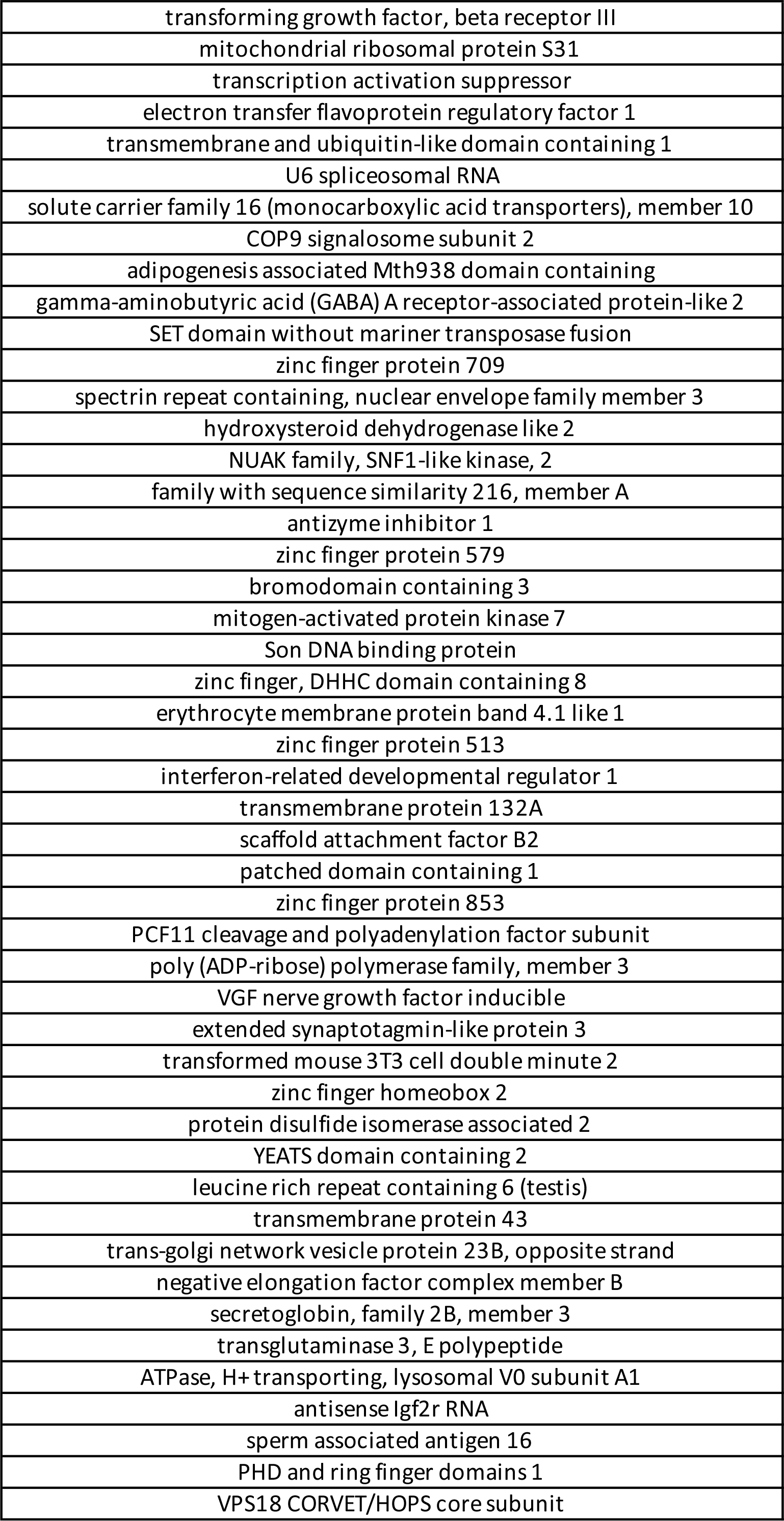
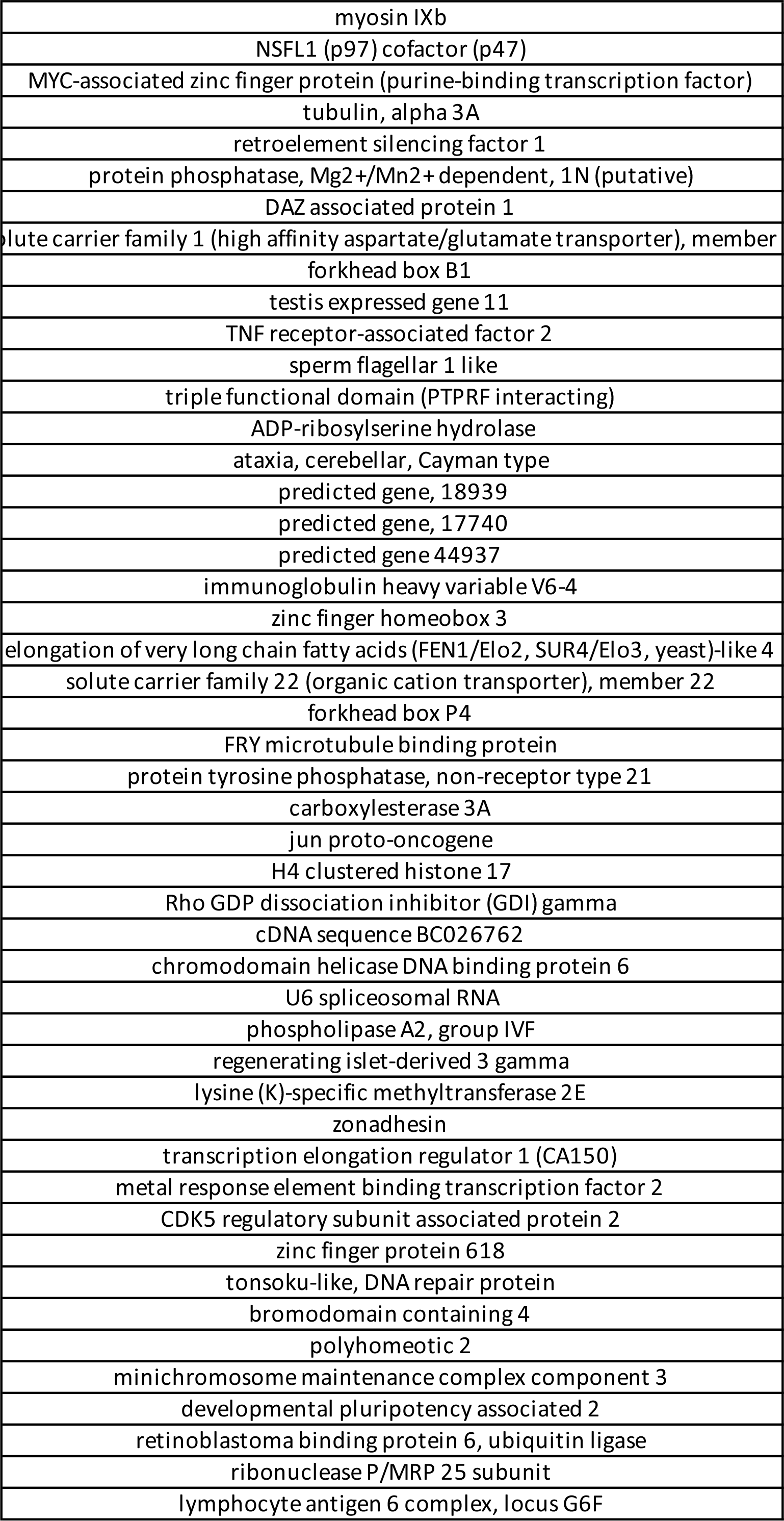
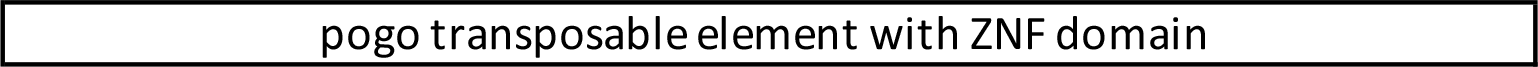
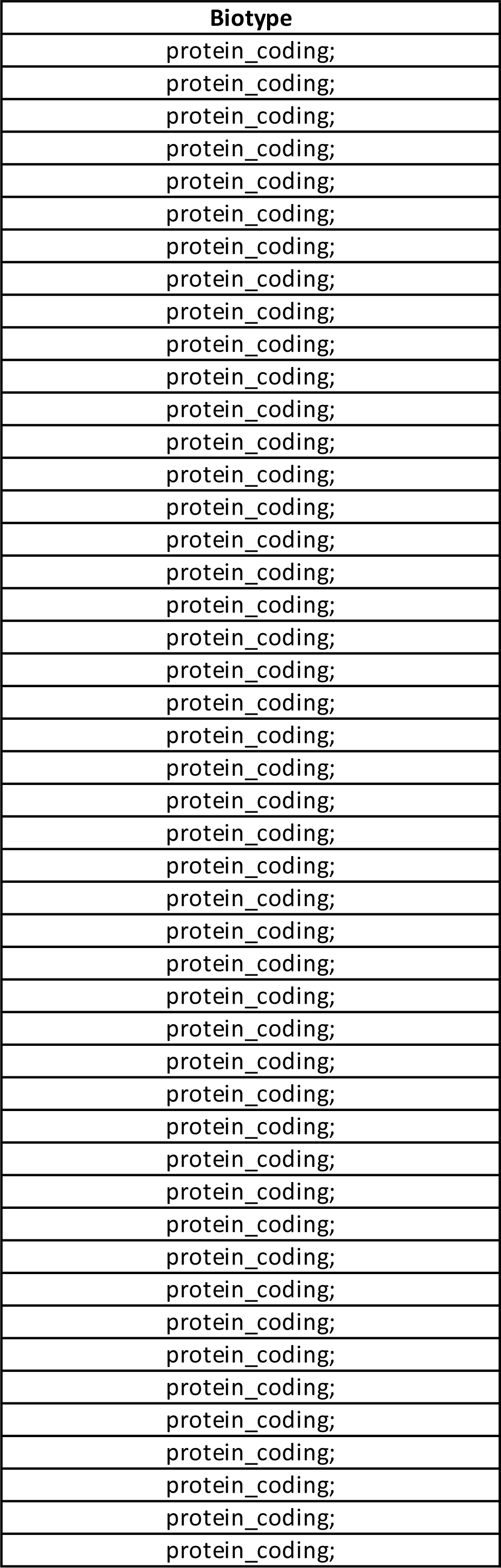
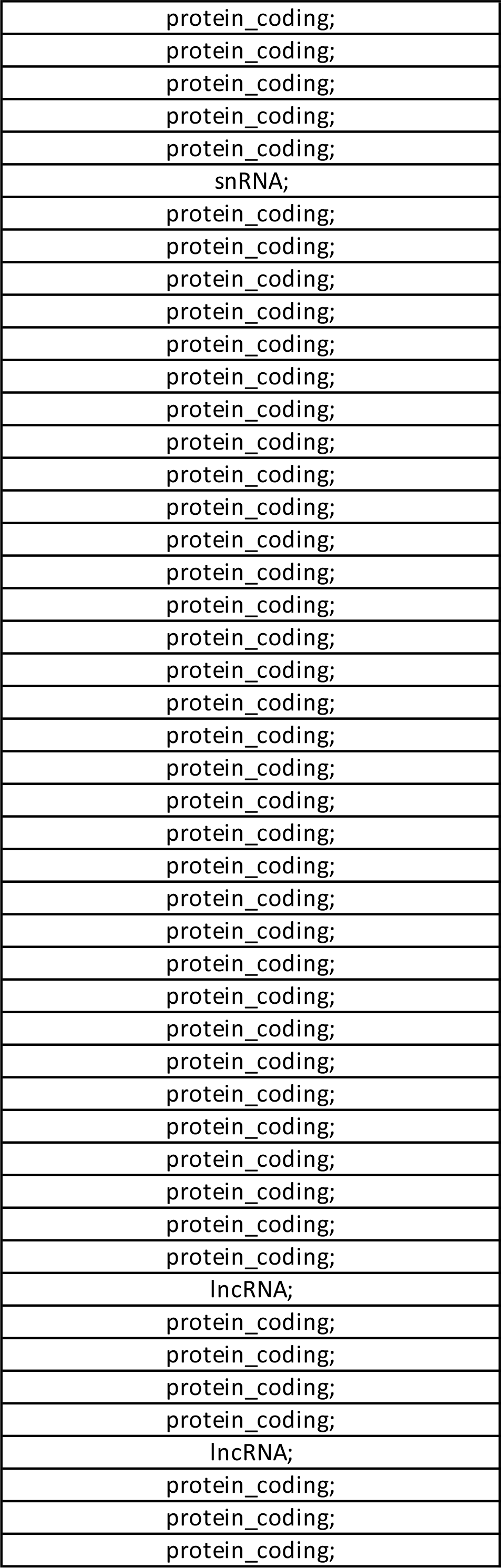
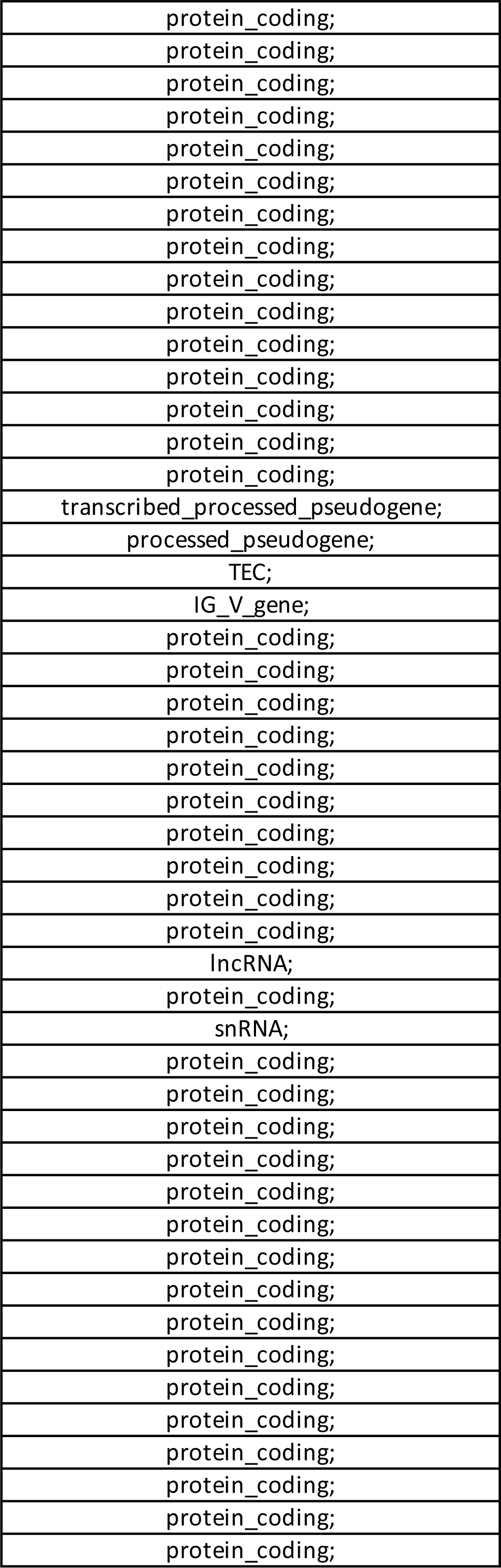
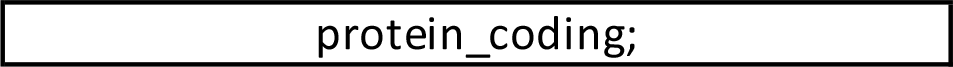

**Table S2:**
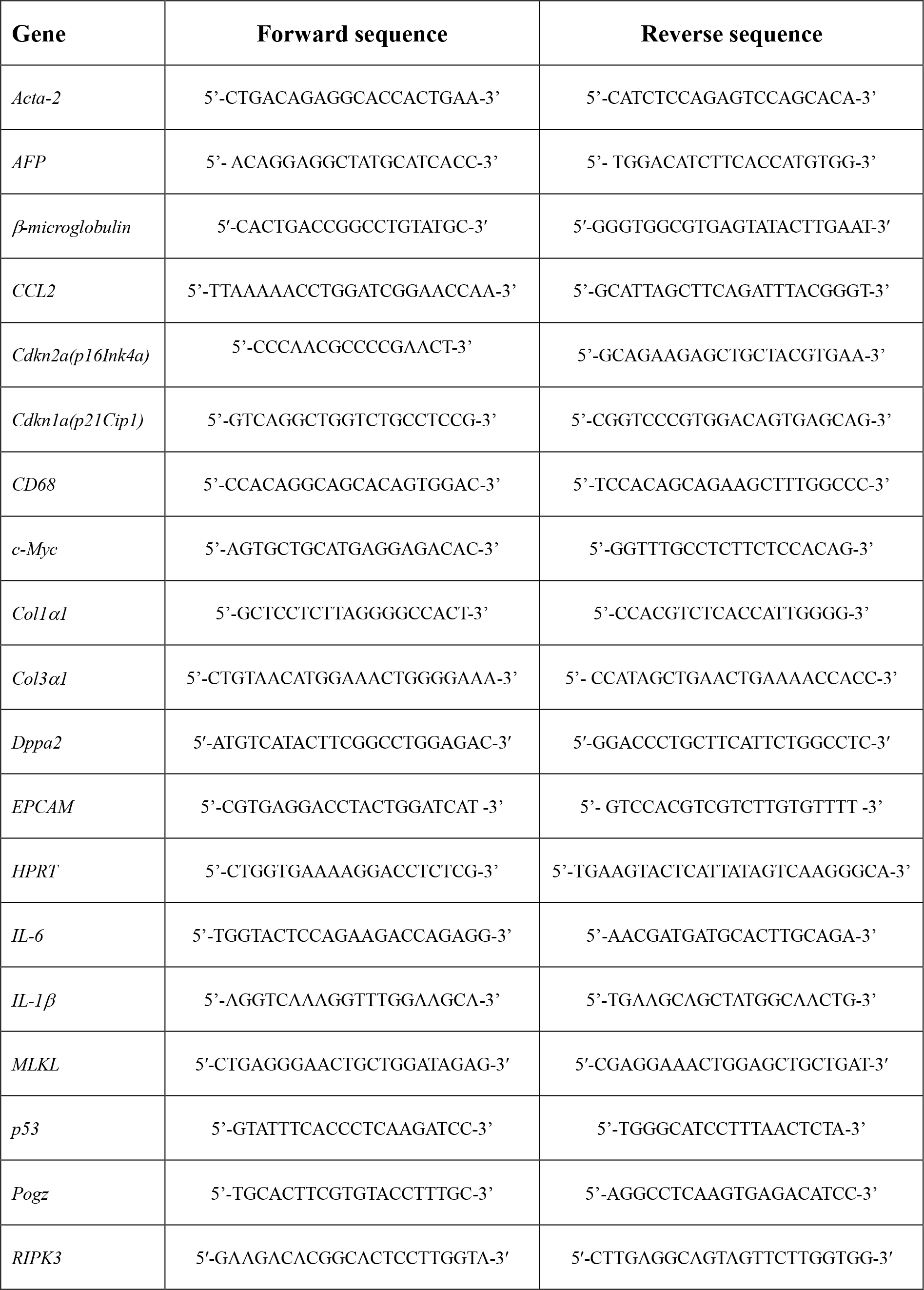
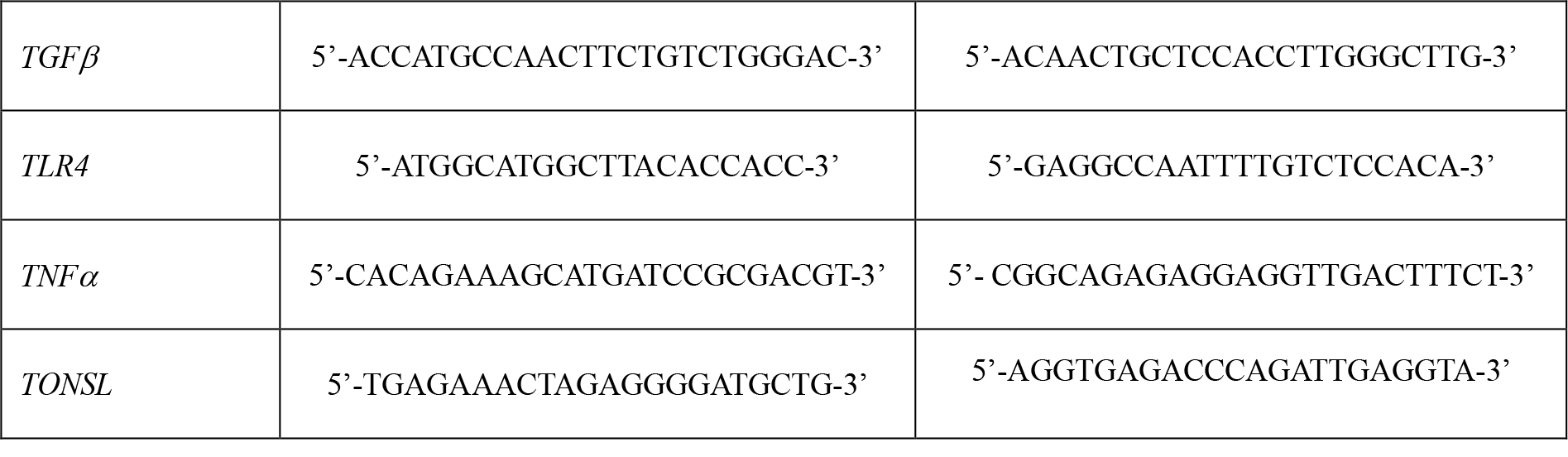
List of primers used for RT-PCR.

## Notes

### Competing Interest Statement

The authors have declared no competing interest.

