## Supplementary figures and images for "Blocking necroptosis reduces inflammation and tumor incidence in a mouse model of diet-induced hepatocellular carcinoma"

### Supplementary Figures

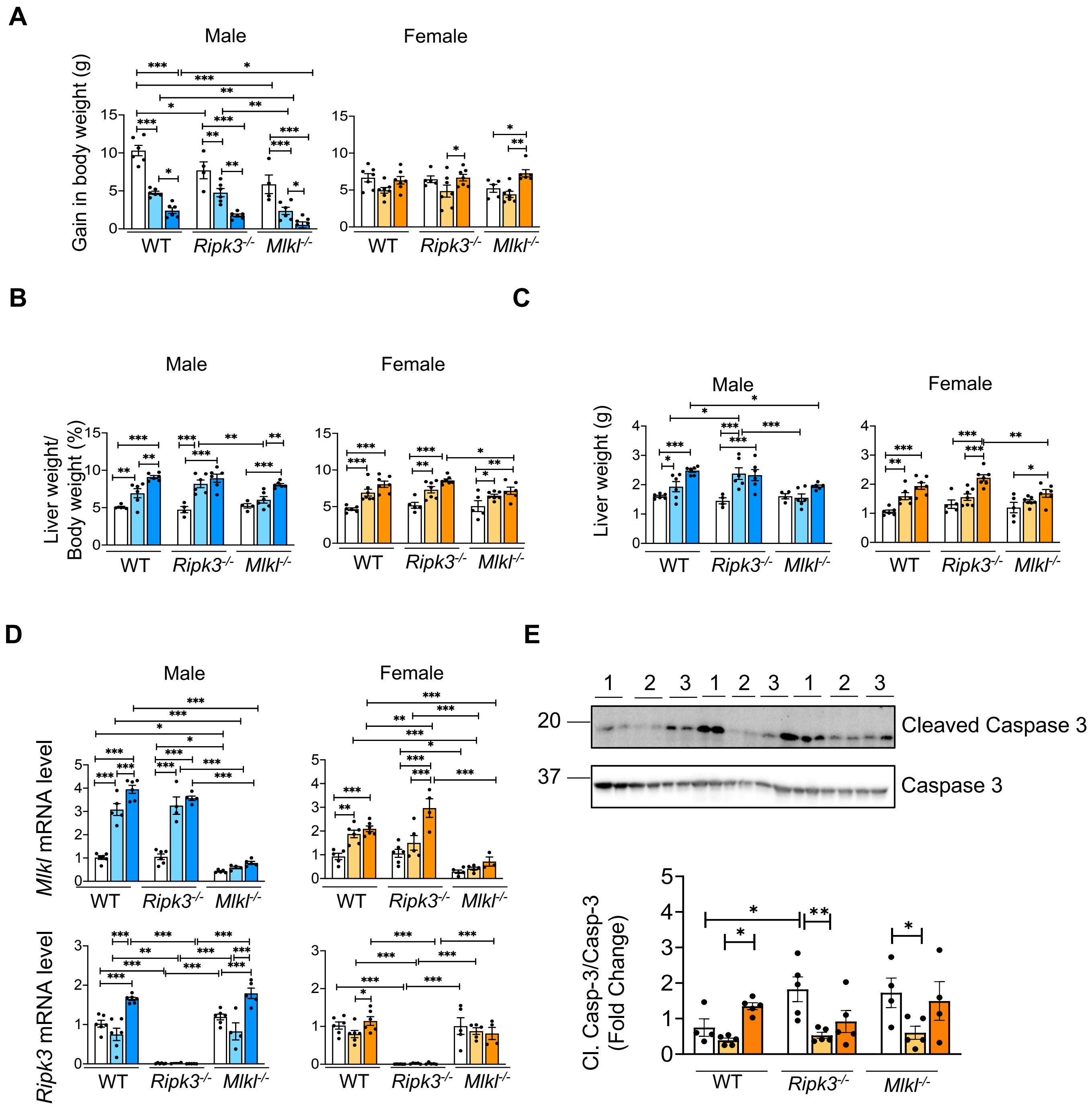

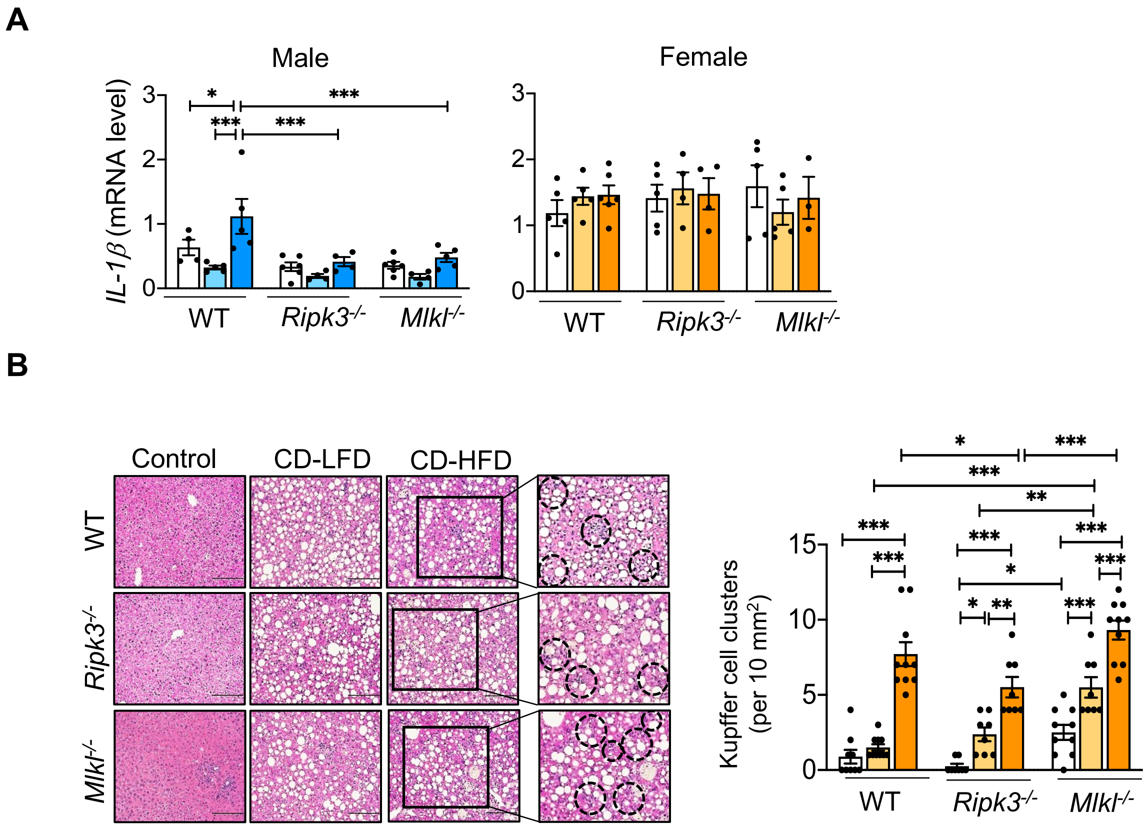

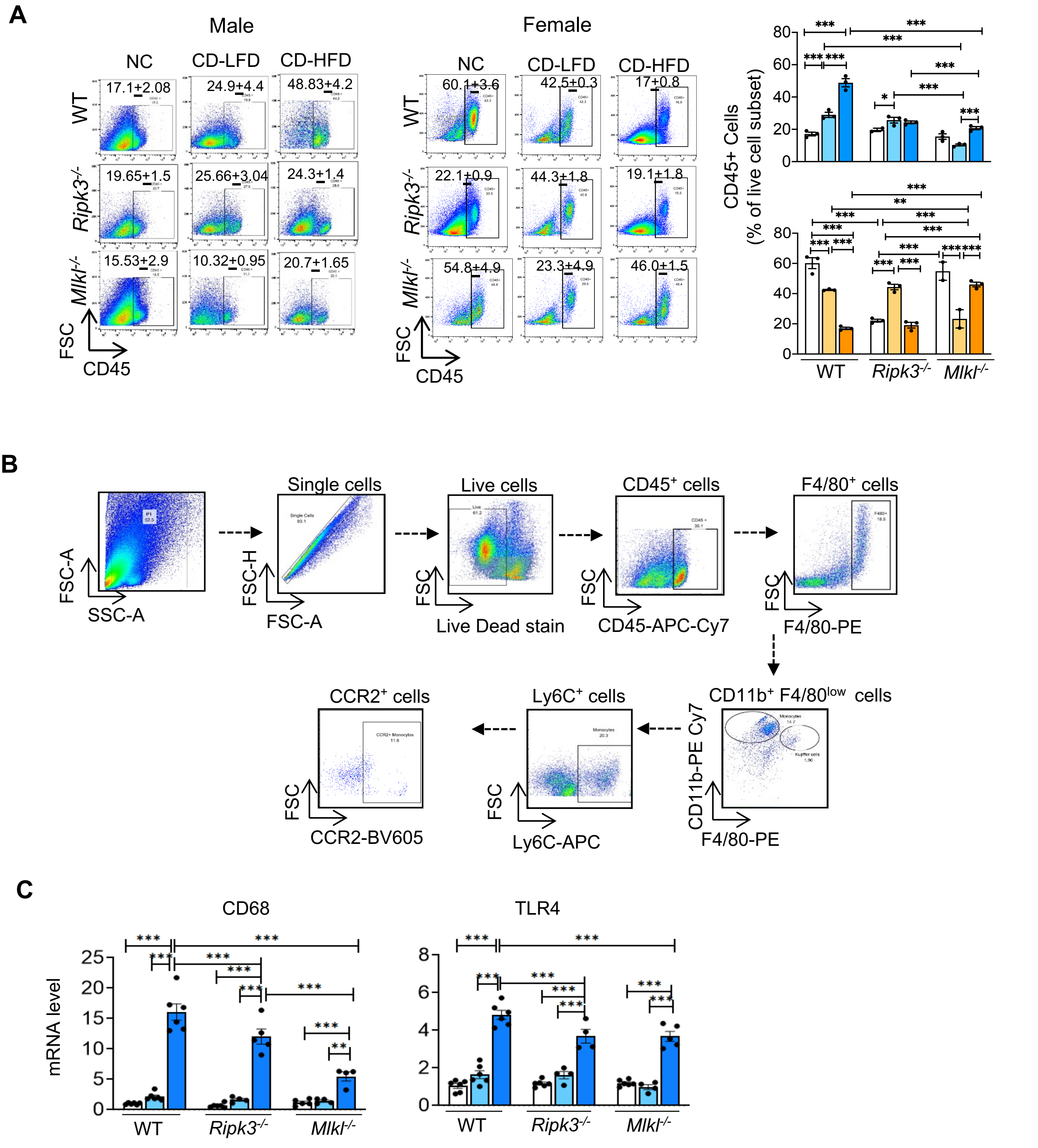

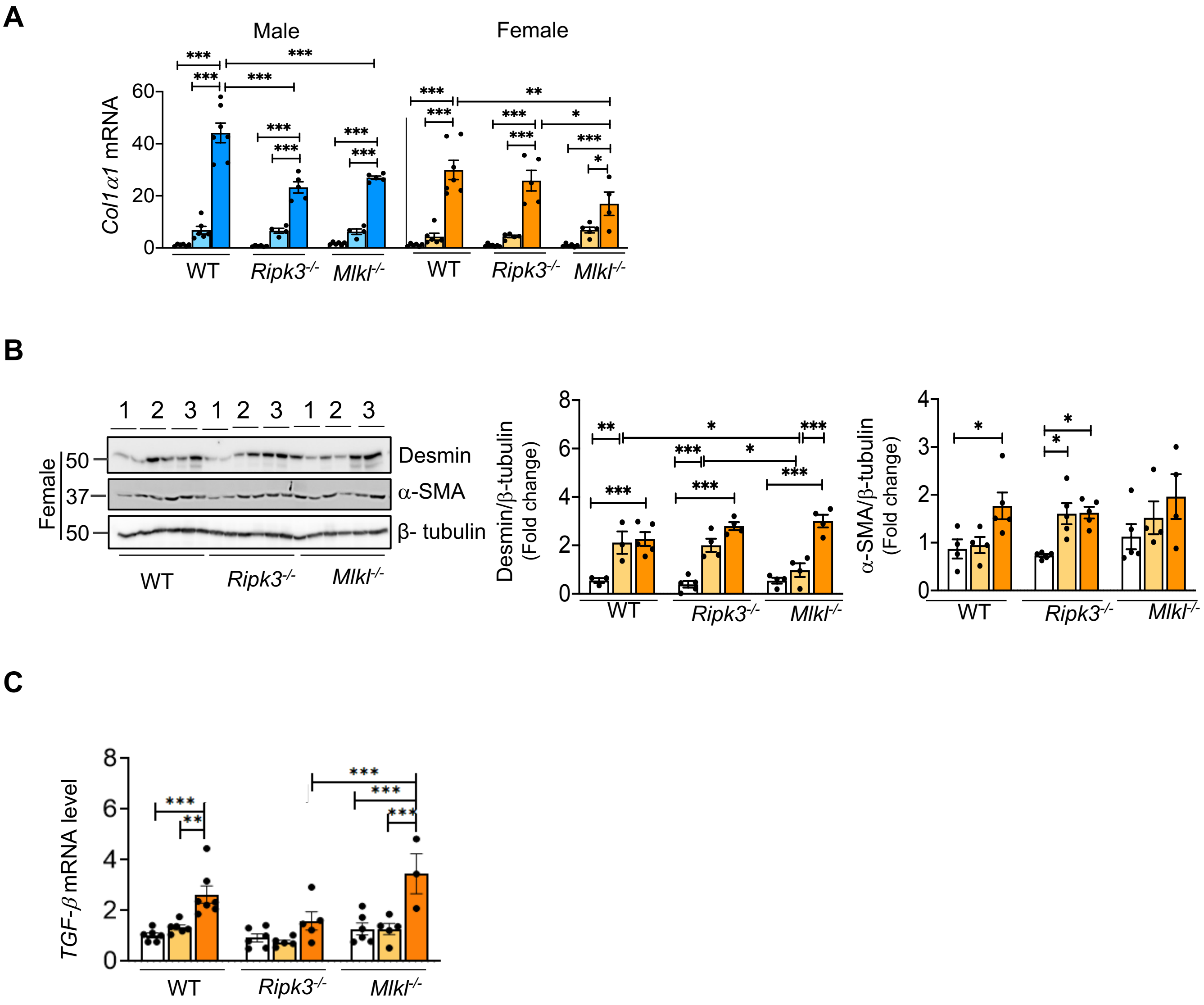

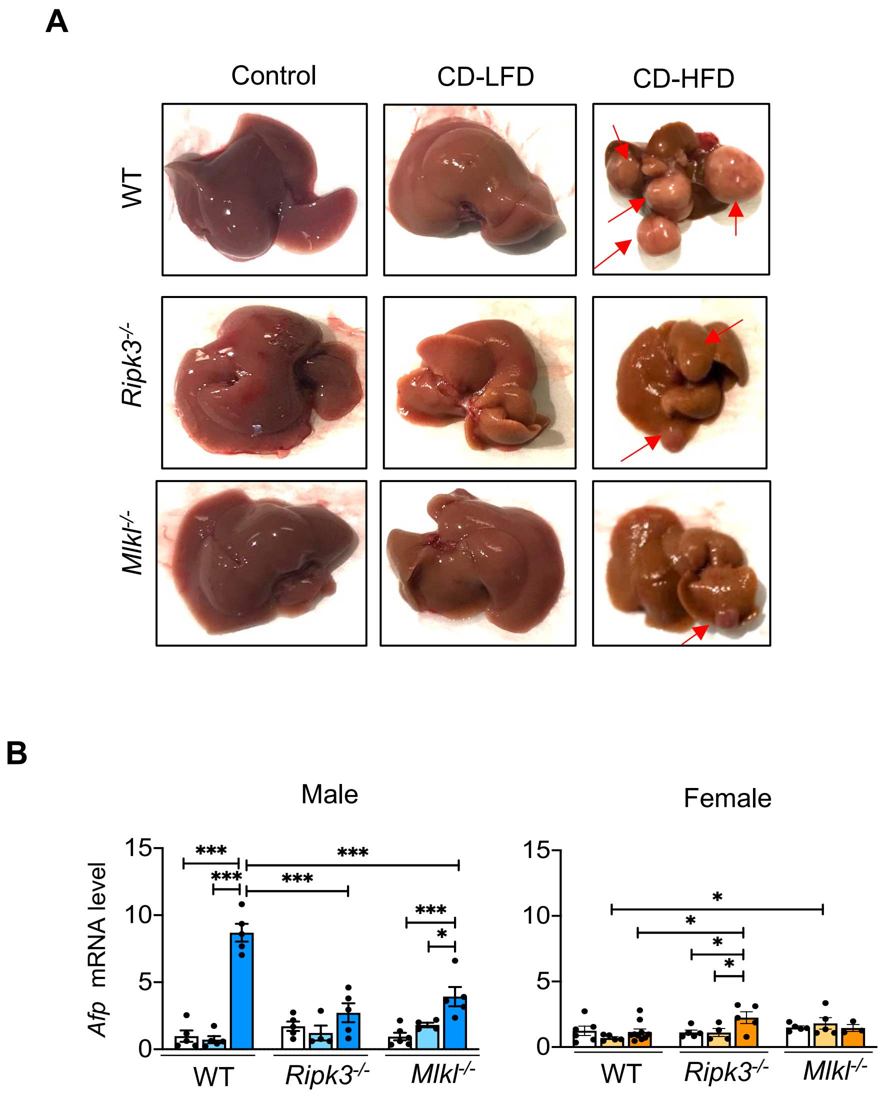

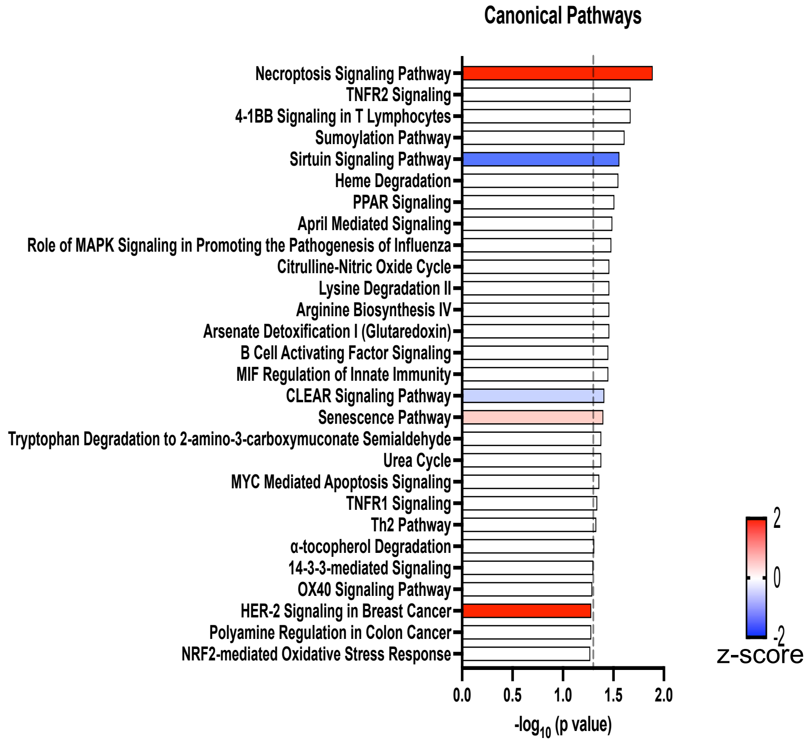

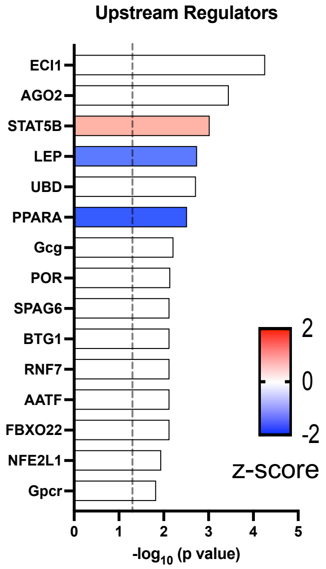

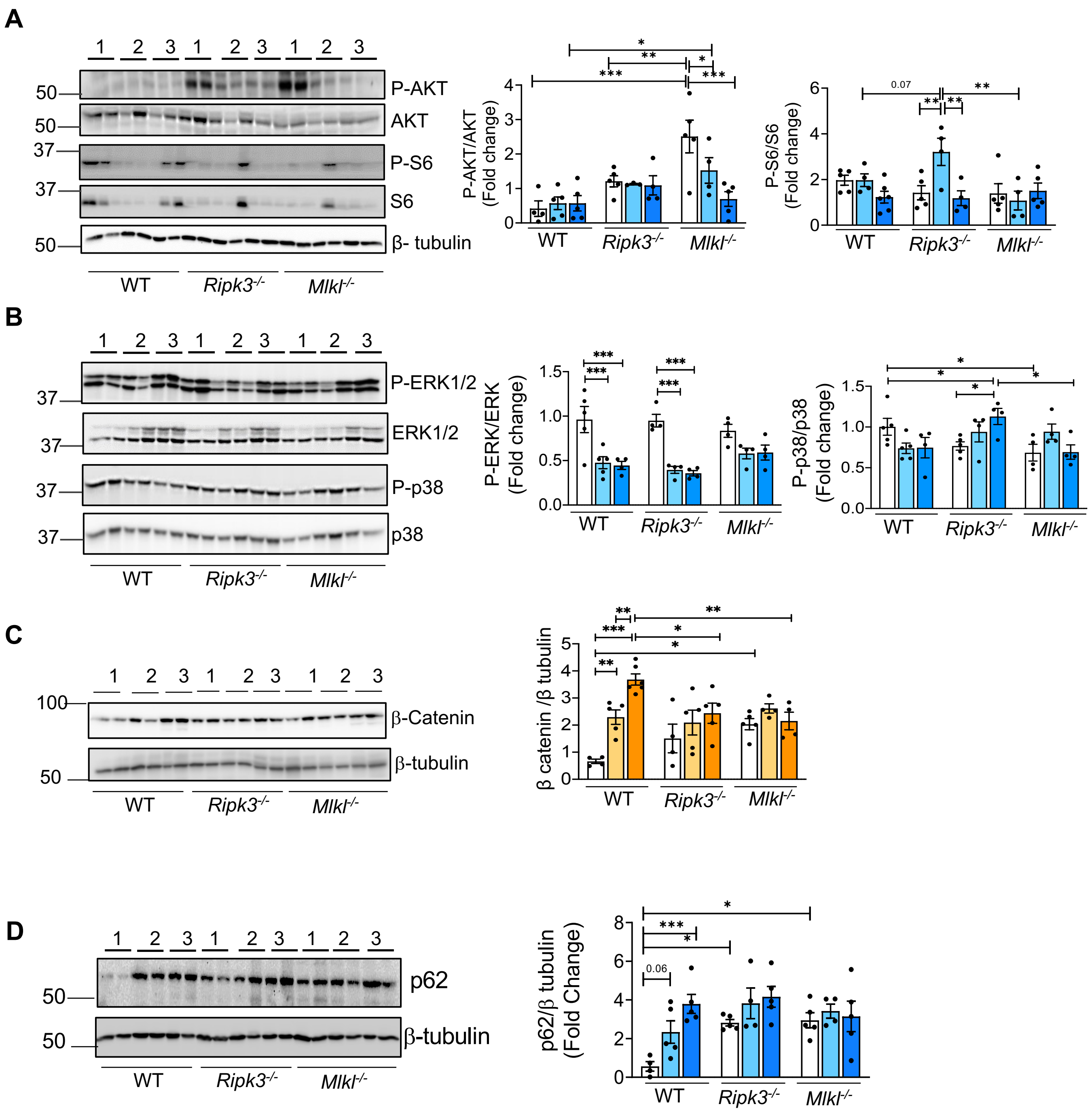
