## Supplementary material for "Blocking necroptosis reduces inflammation and tumor incidence in a mouse model of diet-induced hepatocellular carcinoma": Table S1

| SL.No | WT.C | WT.LFD | WT.HFD | Ripk3ko.C | Ripk3ko.LFD |
|---|---|---|---|---|---|
| 1 | 4.576 | 4.101 | 4.048 | 5.196 | 5.242 |
| 2 | 5.125 | 4.984 | 4.803 | 5.822 | 5.536 |
| 3 | 4.381 | 3.365 | 2.777 | 6.117 | 4.994 |
| 4 | 2.951 | 2.779 | 2.175 | 3.3 | 3.326 |
| 5 | 4.219 | 2.089 | 1.192 | 4.313 | 2.952 |
| 6 | 11.62 | 11.168 | 11.094 | 11.373 | 11.291 |
| 7 | 5.757 | 4.653 | 4.243 | 5.672 | 5.195 |
| 8 | 4.357 | 3.833 | 3.645 | 4.31 | 4.823 |
| 9 | 5.154 | 4.959 | 4.629 | 5.177 | 5.098 |
| 10 | 3.092 | 2.699 | 1.862 | 3.374 | 2.807 |
| 11 | 4.558 | 4.37 | 4.122 | 4.684 | 4.718 |
| 12 | 3.328 | 2.712 | 2.483 | 3.577 | 2.952 |
| 13 | 5.62 | 5.192 | 4.973 | 5.836 | 5.818 |
| 14 | 10.54 | 9.552 | 9.501 | 10.668 | 9.887 |
| 15 | 3.368 | 2.484 | 2.267 | 3.366 | 2.632 |
| 16 | 2.235 | 1.287 | 1.098 | 4.068 | 2.681 |
| 17 | 2.386 | 1.729 | 1.161 | 2.509 | 2.116 |
| 18 | 8.084 | 7.964 | 7.605 | 8.36 | 8.074 |
| 19 | 6.736 | 6.43 | 6.181 | 6.485 | 6.593 |
| 20 | 6.107 | 6.03 | 6.019 | 6.274 | 6.421 |
| 21 | 5.683 | 3.489 | 3.449 | 5.226 | 3.754 |
| 22 | 4.305 | 3.926 | 3.56 | 4.266 | 4.285 |
| 23 | 4.714 | 3.808 | 3.409 | 5.065 | 3.828 |
| 24 | 2.259 | 2.173 | 1.063 | 2.65 | 2.263 |
| 25 | 2.246 | 1.189 | 0.647 | 2.305 | 1.418 |
| 26 | 6.366 | 6.226 | 6.099 | 6.495 | 6.492 |
| 27 | 8.133 | 7.591 | 7.368 | 8.404 | 7.657 |
| 28 | 1.702 | 1.06 | 0.949 | 1.785 | 2.513 |
| 29 | 6.599 | 6.028 | 5.523 | 7.011 | 6.443 |
| 30 | 4.895 | 4.558 | 3.934 | 4.885 | 4.783 |
| 31 | 6.467 | 5.341 | 5.211 | 6.587 | 5.63 |
| 32 | 6.724 | 6.454 | 6.238 | 6.73 | 6.467 |
| 33 | 5.221 | 4.598 | 4.545 | 5.32 | 4.813 |
| 34 | 0.868 | 0.235 | -0.268 | 0.866 | 0.851 |
| 35 | 2.743 | 2.544 | 2.209 | 3.329 | 3.257 |
| 36 | 2.945 | 2.52 | 2.241 | 3.518 | 2.757 |
| 37 | 5.803 | 5.478 | 5.252 | 6.048 | 5.573 |
| 38 | 1.179 | 0.821 | 0.559 | 2.135 | 1.983 |
| 39 | 5.675 | 5.002 | 4.885 | 5.713 | 5.019 |
| 40 | 3.257 | 2.711 | 2.623 | 3.174 | 2.786 |
| 41 | 4.694 | 4.319 | 4.048 | 4.814 | 4.538 |
| 42 | 6.08 | 5.977 | 5.617 | 6.238 | 6.2 |
| 43 | 2.697 | 1.595 | 1.17 | 2.51 | 1.888 |
| 44 | 1.311 | 0.377 | -0.376 | 2.42 | 0.873 |
| 45 | 7.583 | 6.85 | 6.691 | 7.485 | 6.981 |
| 46 | 4.397 | 4.257 | 3.774 | 4.406 | 4.348 |
| 47 | 6.217 | 5.652 | 5.535 | 6.489 | 6.041 |

| | | | | | |
|---|---|---|---|---|---|
| 48 | 3.236 | 2.16 | 2.083 | 2.967 | 2.971 |
| 49 | 5.083 | 4.592 | 4.307 | 5.028 | 4.793 |
| 50 | 4.443 | 4.132 | 3.706 | 4.243 | 4.163 |
| 51 | 4.977 | 4.688 | 4.288 | 4.887 | 4.836 |
| 52 | 3.296 | 3.119 | 2.802 | 3.643 | 3.307 |
| 53 | -1.104 | -2.357 | -3.112 | -0.837 | -1.913 |
| 54 | 6.358 | 6.009 | 5.625 | 6.836 | 6.539 |
| 55 | 5.78 | 5.669 | 5.374 | 5.791 | 5.677 |
| 56 | 3.844 | 3.249 | 2.988 | 4 | 3.762 |
| 57 | 5.481 | 5.176 | 4.798 | 5.551 | 5.554 |
| 58 | 2.202 | 1.784 | 1.015 | 2.536 | 2.002 |
| 59 | 1.919 | 1.418 | 0.826 | 2.063 | 1.678 |
| 60 | 2.698 | 2.634 | 1.898 | 3.229 | 2.838 |
| 61 | 6.094 | 5.844 | 5.474 | 6.24 | 6.009 |
| 62 | 3.922 | 3.758 | 3.726 | 4.473 | 4.199 |
| 63 | 1.064 | 0.387 | 0.236 | 1.099 | 1.123 |
| 64 | 6.256 | 5.839 | 5.553 | 6.428 | 5.921 |
| 65 | 3.395 | 3.734 | 4.651 | 3.085 | 3.696 |
| 66 | 6.089 | 6.278 | 6.349 | 5.666 | 6.244 |
| 67 | 2.194 | 2.502 | 3.115 | 2.222 | 1.882 |
| 68 | 7.86 | 8.346 | 8.535 | 7.848 | 8.134 |
| 69 | 3.484 | 3.703 | 3.984 | 3.311 | 3.674 |
| 70 | 2.461 | 3.848 | 3.863 | 1.286 | 2.883 |
| 71 | 3.88 | 4.243 | 4.411 | 3.839 | 3.973 |
| 72 | 4.292 | 5.144 | 5.346 | 4.299 | 4.65 |
| 73 | 1.384 | 2.317 | 3.422 | 1.282 | 1.867 |
| 74 | 6.602 | 6.956 | 7.408 | 6.639 | 6.77 |
| 75 | -3.752 | -2.847 | -1.217 | -3.796 | -3.71 |
| 76 | -3.797 | -2.958 | -1.994 | -3.796 | -3.772 |
| 77 | 5.062 | 5.286 | 5.559 | 5.106 | 5.198 |
| 78 | 5.356 | 5.893 | 6.049 | 5.332 | 5.837 |
| 79 | -3.797 | -3.334 | -2.803 | -3.796 | -3.81 |
| 80 | -3.406 | -2.425 | -1.213 | -3.136 | -3.4 |
| 81 | 5.165 | 5.648 | 5.817 | 5.22 | 5.182 |
| 82 | 4.329 | 4.978 | 4.993 | 4.009 | 4.73 |
| 83 | -3.797 | -3.719 | -3.478 | -3.796 | -3.81 |
| 84 | 3.902 | 4.252 | 4.371 | 3.603 | 4 |
| 85 | -3.797 | -3.334 | -2.9 | -3.796 | -3.433 |
| 86 | 3.648 | 5.056 | 5.096 | 3.718 | 4.862 |
| 87 | -3.797 | -3.251 | -2.685 | -3.796 | -3.81 |
| 88 | 5.206 | 5.259 | 5.415 | 4.875 | 5.082 |
| 89 | -3.797 | -3.213 | -3.207 | -3.48 | -3.71 |
| 90 | -3.797 | -3.325 | -3.306 | -3.796 | -3.772 |
| 91 | 5.938 | 5.938 | 6.417 | 5.689 | 5.76 |
| 92 | -0.74 | 1.466 | 1.965 | 0.148 | 1.076 |
| 93 | -3.769 | -3.689 | -3.125 | -3.796 | -3.81 |
| 94 | 5.053 | 5.41 | 5.475 | 5.215 | 4.984 |
| 95 | 4.608 | 4.668 | 4.898 | 4.452 | 4.476 |

| | | | | | |
|---|---|---|---|---|---|
| 96 | 5.414 | 6.103 | 6.67 | 5.051 | 5.955 |
| 97 | 6.167 | 6.41 | 6.447 | 6.062 | 6.398 |
| 98 | 6.4 | 6.695 | 6.98 | 6.315 | 6.613 |
| 99 | -3.797 | -3.228 | -3.163 | -3.796 | -3.81 |
| 100 | 5.307 | 5.686 | 5.938 | 4.788 | 5.507 |
| 101 | -3.599 | -2.95 | -2.442 | -3.65 | -3.772 |
| 102 | 5.835 | 6.198 | 6.353 | 5.762 | 6.056 |
| 103 | -3.056 | -2.756 | -2.307 | -3.468 | -3.81 |
| 104 | -3.797 | -3.719 | -3.517 | -3.796 | -3.81 |
| 105 | -3.797 | -3.717 | -2.959 | -3.796 | -3.81 |
| 106 | 4.129 | 4.566 | 4.741 | 4.189 | 4.409 |
| 107 | -3.797 | -3.213 | -3.131 | -3.796 | -3.81 |
| 108 | 4.783 | 5.22 | 5.328 | 4.426 | 4.631 |
| 109 | 4.472 | 4.593 | 4.653 | 4.268 | 4.576 |
| 110 | -3.797 | -3.329 | -2.931 | -3.796 | -3.81 |
| 111 | -3.797 | -3.615 | -3.517 | -3.796 | -3.81 |
| 112 | -3.797 | -3.565 | -3.026 | -3.65 | -3.81 |
| 113 | -3.797 | -3.538 | -3.163 | -3.796 | -3.81 |
| 114 | -3.797 | -3.314 | -3.067 | -3.796 | -3.81 |
| 115 | 4.585 | 4.604 | 4.852 | 4.404 | 4.601 |
| 116 | -3.517 | -3.228 | -3.128 | -3.796 | -3.81 |
| 117 | -3.43 | -3.213 | -2.735 | -3.796 | -3.772 |
| 118 | 5.356 | 5.694 | 5.961 | 5.155 | 5.583 |
| 119 | 1.08 | 2.102 | 2.214 | 1.146 | 1.928 |
| 120 | 4.658 | 4.691 | 4.738 | 4.338 | 4.27 |
| 121 | 11.05 | 11.257 | 11.312 | 10.838 | 10.842 |
| 122 | 5.72 | 6.529 | 7.033 | 5.111 | 5.803 |
| 123 | -3.752 | -2.368 | -1.274 | -2.642 | -3.772 |
| 124 | -3.797 | -3.096 | -2.811 | -3.796 | -3.772 |
| 125 | -3.797 | -3.562 | -3.3 | -3.796 | -3.81 |
| 126 | 4.638 | 4.923 | 5.128 | 4.284 | 4.552 |
| 127 | -3.797 | -3.293 | -3.064 | -3.65 | -3.772 |
| 128 | -3.517 | -2.584 | -1.991 | -3.796 | -3.81 |
| 129 | -3.797 | -3.334 | -2.962 | -3.796 | -3.81 |
| 130 | 5.427 | 6.089 | 6.297 | 5.456 | 5.924 |
| 131 | -3.74 | -1.787 | -0.809 | -3.796 | -3.433 |
| 132 | 4.184 | 4.563 | 4.814 | 4.37 | 4.205 |
| 133 | 3.216 | 3.556 | 3.565 | 3.423 | 3.251 |
| 134 | 2.823 | 3.203 | 3.3 | 2.458 | 2.834 |
| 135 | 0.439 | 1.623 | 2.017 | -0.415 | 1.331 |
| 136 | 1.187 | 1.741 | 2.171 | 0.687 | 1.35 |
| 137 | 6.251 | 6.593 | 6.801 | 6.18 | 6.516 |
| 138 | 5.071 | 5.562 | 5.79 | 5.25 | 5.524 |
| 139 | 2.23 | 2.835 | 3.431 | 2.02 | 2.619 |
| 140 | -3.797 | -3.329 | -3.271 | -3.796 | -3.772 |
| 141 | 6.21 | 6.439 | 6.694 | 6.318 | 6.337 |
| 142 | -3.797 | -2.785 | -2.311 | -3.796 | -3.81 |
| 143 | -3.797 | -3.345 | -2.735 | -3.65 | -3.81 |

| | | | | | |
|---|---|---|---|---|---|
| 144 | 4.558 | 4.736 | 4.834 | 4.589 | 4.477 |

| Ripk3ko.HFD | MLKLko.C | MLKLko.LFD | MLKLko.HFD | gene_id |
|---|---|---|---|---|
| 5.403 | 5.172 | 5.296 | 4.69 | ENSMUSG00000042978 |
| 5.502 | 5.647 | 5.503 | 5.457 | ENSMUSG00000037110 |
| 5.302 | 5.812 | 5.066 | 4.586 | ENSMUSG00000022032 |
| 3.257 | 3.292 | 3.552 | 3.057 | ENSMUSG00000036019 |
| 3.538 | 3.696 | 4.036 | 2.06 | ENSMUSG00000072949 |
| 11.689 | 11.436 | 11.514 | 11.685 | ENSMUSG00000076441 |
| 5.35 | 5.042 | 5.359 | 4.963 | ENSMUSG00000048371 |
| 5.565 | 3.99 | 5.044 | 3.919 | ENSMUSG00000063929 |
| 5.281 | 5.078 | 5.201 | 5.367 | ENSMUSG00000004070 |
| 2.898 | 2.987 | 3.359 | 2.671 | ENSMUSG00000042348 |
| 4.551 | 4.492 | 4.682 | 4.376 | ENSMUSG00000024146 |
| 3.096 | 3.348 | 3.33 | 2.649 | ENSMUSG00000019948 |
| 5.73 | 5.334 | 5.507 | 5.481 | ENSMUSG00000003559 |
| 10.349 | 10.243 | 10.105 | 10.134 | ENSMUSG00000027597 |
| 3.108 | 3.535 | 3.335 | 3.688 | ENSMUSG00000025964 |
| 2.009 | 3.106 | 3.388 | 1.971 | ENSMUSG00000003555 |
| 1.736 | 2.216 | 2.344 | 1.656 | ENSMUSG00000072772 |
| 8.344 | 8.396 | 8.198 | 8.15 | ENSMUSG00000047731 |
| 6.655 | 6.446 | 6.813 | 6.452 | ENSMUSG00000021620 |
| 6.212 | 6.065 | 6.316 | 6.1 | ENSMUSG00000072704 |
| 4.215 | 4.648 | 4.77 | 3.813 | ENSMUSG00000017718 |
| 4.089 | 4.27 | 4.204 | 3.796 | ENSMUSG00000024805 |
| 4.026 | 4.684 | 4.465 | 4.304 | ENSMUSG00000044026 |
| 1.953 | 2.995 | 2.53 | 1.714 | ENSMUSG00000095325 |
| 1.312 | 2.352 | 2.473 | 1.459 | ENSMUSG00000021265 |
| 6.296 | 6.444 | 6.451 | 6.248 | ENSMUSG00000021024 |
| 8.196 | 8.155 | 7.906 | 8.078 | ENSMUSG00000029695 |
| 1.651 | 1.908 | 2.569 | 1.579 | ENSMUSG00000028444 |
| 5.97 | 6.979 | 6.946 | 5.591 | ENSMUSG00000024118 |
| 4.487 | 4.901 | 4.81 | 4.17 | ENSMUSG00000022130 |
| 6.161 | 5.736 | 6.093 | 5.514 | ENSMUSG00000036585 |
| 6.506 | 6.73 | 6.545 | 6.584 | ENSMUSG00000004069 |
| 4.816 | 5.184 | 4.995 | 4.576 | ENSMUSG00000015790 |
| 0.694 | 0.597 | 1.15 | -0.122 | ENSMUSG00000021057 |
| 3.068 | 3.112 | 4.031 | 2.803 | ENSMUSG00000040740 |
| 2.745 | 3.607 | 3.156 | 2.501 | ENSMUSG00000038578 |
| 5.616 | 5.809 | 5.633 | 5.804 | ENSMUSG00000027598 |
| 1.349 | 1.525 | 2.853 | 1.059 | ENSMUSG00000074623 |
| 5.495 | 5.733 | 5.348 | 5.349 | ENSMUSG00000027439 |
| 3.066 | 3.44 | 3.025 | 3.048 | ENSMUSG00000025245 |
| 4.377 | 4.669 | 4.609 | 4.13 | ENSMUSG00000027086 |
| 5.973 | 6.757 | 6.401 | 6.037 | ENSMUSG00000043067 |
| 2.017 | 2.633 | 1.716 | 2.948 | ENSMUSG00000017446 |
| 0.3 | 2.498 | 1.637 | 0.154 | ENSMUSG00000000791 |
| 7.265 | 7.03 | 7.128 | 7.342 | ENSMUSG00000015112 |
| 4.335 | 4.429 | 4.356 | 4.475 | ENSMUSG00000014177 |
| 6.262 | 6.251 | 6.022 | 6.33 | ENSMUSG00000022383 |

| | | | | |
|---|---|---|---|---|
| 2.461 | 2.9 | 2.51 | 3.009 | ENSMUSG00000029287 |
| 4.495 | 5.18 | 4.847 | 4.43 | ENSMUSG00000031533 |
| 4.09 | 4.579 | 4.384 | 4.311 | ENSMUSG00000040651 |
| 4.992 | 4.689 | 4.871 | 4.474 | ENSMUSG00000040370 |
| 3.135 | 3.314 | 3.349 | 3.017 | ENSMUSG00000028958 |
| -1.66 | -1.623 | -0.531 | -1.478 | ENSMUSG00000119317 |
| 6.356 | 6.128 | 6.296 | 6.12 | ENSMUSG00000019838 |
| 5.626 | 5.938 | 5.781 | 5.674 | ENSMUSG00000027206 |
| 3.454 | 3.452 | 3.67 | 3.012 | ENSMUSG00000035642 |
| 5.152 | 5.388 | 5.306 | 5.084 | ENSMUSG00000031950 |
| 1.514 | 2.554 | 2.453 | 1.115 | ENSMUSG00000034639 |
| 1.551 | 2.362 | 1.711 | 1.638 | ENSMUSG00000056019 |
| 2.705 | 2.95 | 2.953 | 2.301 | ENSMUSG00000054150 |
| 6.211 | 5.95 | 6.011 | 5.929 | ENSMUSG00000028383 |
| 4.019 | 3.779 | 4.167 | 3.887 | ENSMUSG00000009772 |
| 0.891 | 1.141 | 0.964 | 1.058 | ENSMUSG00000029463 |
| 6.154 | 6.551 | 6.005 | 6.165 | ENSMUSG00000037458 |
| 4.062 | 2.937 | 3.576 | 3.932 | ENSMUSG00000051550 |
| 6.044 | 5.633 | 6.037 | 5.77 | ENSMUSG00000026918 |
| 2.721 | 1.941 | 2.182 | 2.812 | ENSMUSG00000001034 |
| 8.386 | 8.071 | 8.132 | 8.384 | ENSMUSG00000022961 |
| 3.69 | 3.194 | 3.399 | 3.706 | ENSMUSG00000060166 |
| 3.635 | 1.75 | 2.431 | 3.655 | ENSMUSG00000027624 |
| 4.076 | 4.205 | 3.92 | 3.592 | ENSMUSG00000043059 |
| 4.644 | 4.294 | 4.865 | 4.826 | ENSMUSG00000001627 |
| 2.781 | 1.191 | 1.879 | 3.152 | ENSMUSG00000024736 |
| 6.843 | 6.77 | 6.85 | 6.692 | ENSMUSG00000042625 |
| -1.609 | -3.716 | -3.466 | -3.196 | ENSMUSG00000041552 |
| -2.143 | -3.663 | -3.81 | -2.2 | ENSMUSG00000093910 |
| 5.149 | 5.048 | 5.237 | 5.044 | ENSMUSG00000041328 |
| 5.711 | 5.684 | 5.535 | 5.763 | ENSMUSG00000023249 |
| -3.296 | -3.788 | -3.81 | -3.68 | ENSMUSG00000037428 |
| -2.259 | -3.067 | -3.346 | -2.871 | ENSMUSG00000037681 |
| 5.44 | 5.2 | 5.424 | 5.657 | ENSMUSG00000020184 |
| 4.686 | 3.972 | 4.501 | 4.382 | ENSMUSG00000040721 |
| -3.768 | -3.788 | -3.81 | -3.81 | ENSMUSG00000024184 |
| 4.161 | 3.902 | 3.842 | 4.06 | ENSMUSG00000041215 |
| -3.699 | -3.788 | -3.81 | -3.597 | ENSMUSG00000022375 |
| 4.917 | 3.971 | 4.354 | 5.046 | ENSMUSG00000030095 |
| -3.273 | -3.788 | -3.666 | -3.81 | ENSMUSG00000086677 |
| 5.091 | 5.211 | 5.12 | 5.177 | ENSMUSG00000013465 |
| -3.724 | -3.788 | -3.81 | -3.81 | ENSMUSG00000078754 |
| -3.768 | -3.788 | -3.666 | -3.81 | ENSMUSG00000027401 |
| 6.052 | 5.843 | 5.599 | 6.292 | ENSMUSG00000019302 |
| 0.994 | -0.387 | 0.268 | 0.965 | ENSMUSG00000078247 |
| -3.81 | -3.788 | -3.81 | -3.81 | ENSMUSG00000053153 |
| 5.155 | 5.395 | 5.151 | 5.267 | ENSMUSG00000038611 |
| 4.72 | 4.548 | 4.316 | 4.413 | ENSMUSG00000034216 |

| | | | | |
|---|---|---|---|---|
| 6.42 | 5.248 | 5.725 | 6.566 | ENSMUSG00000004677 |
| 6.207 | 6.288 | 6.121 | 6.089 | ENSMUSG00000027455 |
| 6.554 | 6.299 | 6.599 | 6.632 | ENSMUSG00000030678 |
| -3.81 | -3.657 | -3.272 | -3.81 | ENSMUSG00000067702 |
| 5.51 | 4.555 | 4.735 | 5.212 | ENSMUSG00000032712 |
| -3.757 | -3.788 | -3.272 | -3.652 | ENSMUSG00000030402 |
| 6.033 | 5.909 | 5.956 | 6.01 | ENSMUSG00000069565 |
| -2.976 | -3.788 | -3.347 | -3.68 | ENSMUSG00000005357 |
| -3.81 | -3.788 | -3.81 | -3.81 | ENSMUSG00000059246 |
| -3.81 | -3.788 | -3.81 | -3.81 | ENSMUSG00000009670 |
| 4.067 | 4.275 | 4.402 | 4.265 | ENSMUSG00000026942 |
| -3.724 | -3.703 | -3.416 | -3.68 | ENSMUSG00000073795 |
| 5.104 | 4.715 | 4.565 | 5.126 | ENSMUSG00000022263 |
| 4.376 | 4.324 | 4.231 | 4.216 | ENSMUSG00000042558 |
| -3.719 | -3.788 | -3.81 | -3.597 | ENSMUSG00000034958 |
| -3.81 | -3.788 | -3.81 | -3.81 | ENSMUSG00000114147 |
| -3.81 | -3.788 | -3.81 | -3.81 | ENSMUSG00000114316 |
| -3.81 | -3.788 | -3.81 | -3.81 | ENSMUSG00000109434 |
| -3.768 | -3.788 | -3.81 | -3.57 | ENSMUSG00000094174 |
| 4.209 | 4.411 | 4.284 | 4.209 | ENSMUSG00000038872 |
| -3.64 | -3.788 | -3.81 | -3.81 | ENSMUSG00000032262 |
| -3.81 | -3.788 | -3.558 | -3.81 | ENSMUSG00000022366 |
| 5.632 | 5.46 | 5.461 | 5.627 | ENSMUSG00000023991 |
| 1.717 | 1.219 | 1.289 | 1.968 | ENSMUSG00000056602 |
| 4.385 | 4.076 | 4.195 | 4.414 | ENSMUSG00000021009 |
| 11.119 | 10.495 | 10.89 | 11.247 | ENSMUSG00000069922 |
| 6.529 | 5.067 | 5.503 | 6.255 | ENSMUSG00000052684 |
| -2.689 | -3.657 | -2.693 | -3.358 | ENSMUSG00000069306 |
| -3.704 | -3.788 | -3.666 | -3.68 | ENSMUSG00000073433 |
| -3.81 | -3.788 | -3.81 | -3.81 | ENSMUSG00000108413 |
| 4.739 | 4.505 | 4.605 | 4.595 | ENSMUSG00000057133 |
| -3.768 | -3.788 | -3.81 | -3.68 | ENSMUSG00000119598 |
| -2.913 | -3.207 | -3.466 | -3.607 | ENSMUSG00000046971 |
| -3.757 | -3.629 | -3.81 | -3.81 | ENSMUSG00000030017 |
| 5.818 | 5.621 | 5.593 | 5.857 | ENSMUSG00000029004 |
| -2.512 | -3.788 | -3.298 | -1.655 | ENSMUSG00000079173 |
| 4.411 | 4.475 | 4.185 | 4.355 | ENSMUSG00000024498 |
| 3.191 | 3.378 | 3.274 | 2.961 | ENSMUSG00000029267 |
| 2.928 | 2.604 | 2.598 | 2.974 | ENSMUSG00000039298 |
| 1.075 | -0.185 | 0.507 | 1.136 | ENSMUSG00000028358 |
| 1.35 | 1.169 | 1.281 | 1.301 | ENSMUSG00000059323 |
| 6.352 | 6.315 | 6.221 | 6.196 | ENSMUSG00000024002 |
| 5.466 | 5.282 | 5.309 | 5.421 | ENSMUSG00000028796 |
| 2.744 | 2.332 | 2.136 | 3.014 | ENSMUSG00000041859 |
| -3.81 | -3.788 | -3.81 | -3.81 | ENSMUSG00000072419 |
| 6.208 | 6.21 | 6.247 | 6.05 | ENSMUSG00000030779 |
| -2.804 | -3.788 | -3.558 | -3.617 | ENSMUSG00000062309 |
| -3.81 | -3.788 | -3.81 | -3.81 | ENSMUSG00000034923 |

| | | | | |
|---|---|---|---|---|
| 4.412 | 4.471 | 4.478 | 4.154 | ENSMUSG00000038902 |

| sample_1 | sample_2 | value_1 | value_2 | lgFold |
|---|---|---|---|---|
| MR:D012 | WT:D012 | 0.103 | -0.675 | -0.778 |
| MR:D012 | WT:D012 | -0.064 | -0.461 | -0.397 |
| MR:D012 | WT:D012 | -0.54 | -1.726 | -1.186 |
| MR:D012 | WT:D012 | -0.009 | -0.538 | -0.529 |
| MR:D012 | WT:D012 | -0.357 | -1.638 | -1.281 |
| MR:D012 | WT:D012 | 0.119 | -0.195 | -0.314 |
| MR:D012 | WT:D012 | -0.144 | -0.679 | -0.535 |
| MR:D012 | WT:D012 | 0.475 | -0.382 | -0.857 |
| MR:D012 | WT:D012 | 0.093 | -0.219 | -0.312 |
| MR:D012 | WT:D012 | -0.189 | -0.607 | -0.417 |
| MR:D012 | WT:D012 | -0.006 | -0.25 | -0.244 |
| MR:D012 | WT:D012 | -0.17 | -0.545 | -0.376 |
| MR:D012 | WT:D012 | -0.079 | -0.349 | -0.269 |
| MR:D012 | WT:D012 | -0.202 | -0.584 | -0.383 |
| MR:D012 | WT:D012 | -0.174 | -0.643 | -0.468 |
| MR:D012 | WT:D012 | -0.702 | -1.541 | -0.84 |
| MR:D012 | WT:D012 | -0.327 | -0.645 | -0.318 |
| MR:D012 | WT:D012 | -0.081 | -0.334 | -0.252 |
| MR:D012 | WT:D012 | 0.031 | -0.185 | -0.216 |
| MR:D012 | WT:D012 | 0.061 | -0.115 | -0.175 |
| MR:D012 | WT:D012 | -0.526 | -1.069 | -0.543 |
| MR:D012 | WT:D012 | -0.157 | -0.376 | -0.218 |
| MR:D012 | WT:D012 | -0.432 | -0.774 | -0.343 |
| MR:D012 | WT:D012 | -0.427 | -0.737 | -0.31 |
| MR:D012 | WT:D012 | -0.56 | -0.952 | -0.392 |
| MR:D012 | WT:D012 | -0.068 | -0.201 | -0.134 |
| MR:D012 | WT:D012 | -0.152 | -0.449 | -0.297 |
| MR:D012 | WT:D012 | -0.054 | -0.666 | -0.612 |
| MR:D012 | WT:D012 | -0.491 | -0.801 | -0.31 |
| MR:D012 | WT:D012 | -0.225 | -0.461 | -0.237 |
| MR:D012 | WT:D012 | -0.187 | -0.625 | -0.438 |
| MR:D012 | WT:D012 | -0.115 | -0.235 | -0.121 |
| MR:D012 | WT:D012 | -0.238 | -0.426 | -0.188 |
| MR:D012 | WT:D012 | -0.074 | -0.593 | -0.519 |
| MR:D012 | WT:D012 | -0.077 | -0.596 | -0.519 |
| MR:D012 | WT:D012 | -0.367 | -0.691 | -0.324 |
| MR:D012 | WT:D012 | -0.131 | -0.32 | -0.189 |
| MR:D012 | WT:D012 | -0.208 | -0.849 | -0.642 |
| MR:D012 | WT:D012 | -0.208 | -0.449 | -0.241 |
| MR:D012 | WT:D012 | -0.155 | -0.377 | -0.222 |
| MR:D012 | WT:D012 | -0.188 | -0.372 | -0.184 |
| MR:D012 | WT:D012 | -0.241 | -0.441 | -0.199 |
| MR:D012 | WT:D012 | -0.243 | -0.704 | -0.461 |
| MR:D012 | WT:D012 | -1.036 | -1.534 | -0.498 |
| MR:D012 | WT:D012 | -0.07 | -0.339 | -0.269 |
| MR:D012 | WT:D012 | -0.029 | -0.264 | -0.235 |
| MR:D012 | WT:D012 | -0.104 | -0.438 | -0.334 |

| | | | | |
|---|---|---|---|---|
| MR:D012 | WT:D012 | -0.216 | -0.559 | -0.342 |
| MR:D012 | WT:D012 | -0.301 | -0.437 | -0.135 |
| MR:D012 | WT:D012 | -0.186 | -0.341 | -0.155 |
| MR:D012 | WT:D012 | -0.072 | -0.249 | -0.177 |
| MR:D012 | WT:D012 | -0.143 | -0.328 | -0.185 |
| MR:D012 | WT:D012 | -0.317 | -1.041 | -0.723 |
| MR:D012 | WT:D012 | -0.18 | -0.443 | -0.263 |
| MR:D012 | WT:D012 | -0.122 | -0.219 | -0.097 |
| MR:D012 | WT:D012 | -0.211 | -0.459 | -0.248 |
| MR:D012 | WT:D012 | -0.201 | -0.342 | -0.141 |
| MR:D012 | WT:D012 | -0.513 | -0.755 | -0.242 |
| MR:D012 | WT:D012 | -0.344 | -0.669 | -0.325 |
| MR:D012 | WT:D012 | -0.189 | -0.512 | -0.323 |
| MR:D012 | WT:D012 | -0.078 | -0.283 | -0.205 |
| MR:D012 | WT:D012 | -0.004 | -0.25 | -0.246 |
| MR:D012 | WT:D012 | -0.066 | -0.513 | -0.447 |
| MR:D012 | WT:D012 | -0.247 | -0.445 | -0.198 |
| MR:D012 | WT:D012 | 0.451 | 0.732 | 0.28 |
| MR:D012 | WT:D012 | 0.117 | 0.292 | 0.176 |
| MR:D012 | WT:D012 | 0.28 | 0.542 | 0.262 |
| MR:D012 | WT:D012 | 0.188 | 0.326 | 0.139 |
| MR:D012 | WT:D012 | 0.202 | 0.344 | 0.142 |
| MR:D012 | WT:D012 | 0.76 | 1.164 | 0.404 |
| MR:D012 | WT:D012 | 0.02 | 0.207 | 0.187 |
| MR:D012 | WT:D012 | 0.34 | 0.548 | 0.208 |
| MR:D012 | WT:D012 | 0.841 | 1.123 | 0.282 |
| MR:D012 | WT:D012 | 0.105 | 0.329 | 0.224 |
| MR:D012 | WT:D012 | 0.569 | 1.125 | 0.556 |
| MR:D012 | WT:D012 | 0.358 | 0.9 | 0.541 |
| MR:D012 | WT:D012 | 0.078 | 0.216 | 0.138 |
| MR:D012 | WT:D012 | 0.12 | 0.298 | 0.178 |
| MR:D012 | WT:D012 | 0.214 | 0.502 | 0.288 |
| MR:D012 | WT:D012 | 0.185 | 1.067 | 0.881 |
| MR:D012 | WT:D012 | 0.215 | 0.35 | 0.135 |
| MR:D012 | WT:D012 | 0.247 | 0.498 | 0.251 |
| MR:D012 | WT:D012 | 0.013 | 0.143 | 0.13 |
| MR:D012 | WT:D012 | 0.167 | 0.327 | 0.16 |
| MR:D012 | WT:D012 | 0.075 | 0.464 | 0.389 |
| MR:D012 | WT:D012 | 0.5 | 0.723 | 0.223 |
| MR:D012 | WT:D012 | 0.123 | 0.551 | 0.428 |
| MR:D012 | WT:D012 | 0.031 | 0.161 | 0.13 |
| MR:D012 | WT:D012 | -0.011 | 0.416 | 0.427 |
| MR:D012 | WT:D012 | 0.002 | 0.32 | 0.317 |
| MR:D012 | WT:D012 | 0.159 | 0.315 | 0.156 |
| MR:D012 | WT:D012 | 0.524 | 1.111 | 0.587 |
| MR:D012 | WT:D012 | -0.007 | 0.173 | 0.18 |
| MR:D012 | WT:D012 | -0.039 | 0.144 | 0.184 |
| MR:D012 | WT:D012 | 0.1 | 0.195 | 0.095 |

| | | | | |
|---|---|---|---|---|
| MR:D012 | WT:D012 | 0.596 | 0.755 | 0.159 |
| MR:D012 | WT:D012 | 0.019 | 0.143 | 0.124 |
| MR:D012 | WT:D012 | 0.189 | 0.311 | 0.122 |
| MR:D012 | WT:D012 | -0.011 | 0.339 | 0.35 |
| MR:D012 | WT:D012 | 0.325 | 0.594 | 0.269 |
| MR:D012 | WT:D012 | 0.058 | 0.668 | 0.61 |
| MR:D012 | WT:D012 | 0.074 | 0.27 | 0.195 |
| MR:D012 | WT:D012 | 0.102 | 0.777 | 0.674 |
| MR:D012 | WT:D012 | -0.004 | 0.126 | 0.131 |
| MR:D012 | WT:D012 | -0.003 | 0.196 | 0.199 |
| MR:D012 | WT:D012 | 0.076 | 0.254 | 0.178 |
| MR:D012 | WT:D012 | 0.052 | 0.468 | 0.416 |
| MR:D012 | WT:D012 | 0.213 | 0.44 | 0.227 |
| MR:D012 | WT:D012 | 0.026 | 0.166 | 0.14 |
| MR:D012 | WT:D012 | 0.084 | 0.471 | 0.388 |
| MR:D012 | WT:D012 | -0.01 | 0.144 | 0.154 |
| MR:D012 | WT:D012 | -0.01 | 0.329 | 0.339 |
| MR:D012 | WT:D012 | -0.006 | 0.305 | 0.311 |
| MR:D012 | WT:D012 | 0.087 | 0.449 | 0.362 |
| MR:D012 | WT:D012 | -0.09 | 0.173 | 0.263 |
| MR:D012 | WT:D012 | 0.033 | 0.474 | 0.442 |
| MR:D012 | WT:D012 | -0.059 | 0.483 | 0.542 |
| MR:D012 | WT:D012 | 0.123 | 0.319 | 0.197 |
| MR:D012 | WT:D012 | 0.279 | 0.596 | 0.316 |
| MR:D012 | WT:D012 | 0.016 | 0.258 | 0.242 |
| MR:D012 | WT:D012 | 0.141 | 0.346 | 0.206 |
| MR:D012 | WT:D012 | 0.584 | 0.969 | 0.385 |
| MR:D012 | WT:D012 | 0.35 | 1.231 | 0.882 |
| MR:D012 | WT:D012 | 0.062 | 0.549 | 0.487 |
| MR:D012 | WT:D012 | -0.008 | 0.244 | 0.251 |
| MR:D012 | WT:D012 | 0.084 | 0.355 | 0.271 |
| MR:D012 | WT:D012 | 0.033 | 0.411 | 0.378 |
| MR:D012 | WT:D012 | 0.127 | 0.952 | 0.826 |
| MR:D012 | WT:D012 | 0 | 0.418 | 0.419 |
| MR:D012 | WT:D012 | 0.152 | 0.42 | 0.267 |
| MR:D012 | WT:D012 | 0.739 | 1.622 | 0.884 |
| MR:D012 | WT:D012 | 0.022 | 0.241 | 0.219 |
| MR:D012 | WT:D012 | -0.101 | 0.121 | 0.223 |
| MR:D012 | WT:D012 | 0.147 | 0.404 | 0.257 |
| MR:D012 | WT:D012 | 0.491 | 1.039 | 0.548 |
| MR:D012 | WT:D012 | 0.158 | 0.577 | 0.419 |
| MR:D012 | WT:D012 | 0.059 | 0.275 | 0.216 |
| MR:D012 | WT:D012 | 0.13 | 0.286 | 0.156 |
| MR:D012 | WT:D012 | 0.313 | 0.651 | 0.338 |
| MR:D012 | WT:D012 | -0.009 | 0.387 | 0.396 |
| MR:D012 | WT:D012 | -0.051 | 0.2 | 0.251 |
| MR:D012 | WT:D012 | 0.202 | 0.924 | 0.722 |
| MR:D012 | WT:D012 | -0.015 | 0.495 | 0.511 |

| | | | | |
|---|---|---|---|---|
| MR:D012 | WT:D012 | -0.075 | 0.151 | 0.226 |

| test_stat | p_value | q_value | SYMBOL |
|---|---|---|---|
| -5.265 | 4.59E-06 | 0.000825 | Sbk1 |
| -4.674 | 3.09E-05 | 0.00255 | Ralgapa2 |
| -4.625 | 3.61E-05 | 0.00263 | Scara5 |
| -4.611 | 3.78E-05 | 0.000983 | Tmtc2 |
| -4.355 | 8.48E-05 | 0.00202 | Acot1 |
| -4.334 | 9.05E-05 | 0.00475 | Ass1 |
| -4.153 | 0.000159 | 0.00643 | Pdp2 |
| -4.071 | 0.000204 | 0.00747 | Cyp4a32 |
| -3.77 | 0.000508 | 0.0123 | Hmox2 |
| -3.742 | 0.000552 | 0.00868 | Arl15 |
| -3.722 | 0.000587 | 0.0133 | Cript |
| -3.663 | 0.000697 | 0.0149 | Actr6 |
| -3.534 | 0.00102 | 0.0183 | As3mt |
| -3.53 | 0.00103 | 0.0185 | Ahcy |
| -3.518 | 0.00107 | 0.0142 | Adam23 |
| -3.511 | 0.00109 | 0.0144 | Cyp17a1 |
| -3.433 | 0.00136 | 0.017 | Grcc10 |
| -3.427 | 0.00138 | 0.0215 | Wbp1l |
| -3.373 | 0.00161 | 0.0192 | Acot12 |
| -3.306 | 0.00195 | 0.0258 | Smim10l1 |
| -3.286 | 0.00207 | 0.0227 | Afmid |
| -3.243 | 0.00234 | 0.0285 | Pcgf5 |
| -3.167 | 0.00288 | 0.0285 | Slc35g1 |
| -3.123 | 0.00325 | 0.0311 | Zfp870 |
| -3.115 | 0.00332 | 0.0314 | Slc25a29 |
| -3.113 | 0.00334 | 0.0315 | Psma6 |
| -3.11 | 0.00337 | 0.0359 | Aass |
| -3.094 | 0.00352 | 0.0327 | Cntfr |
| -3.081 | 0.00365 | 0.0334 | Tedc2 |
| -3.026 | 0.00423 | 0.0405 | Tgds |
| -3.006 | 0.00447 | 0.0418 | Fgf1 |
| -3.006 | 0.00448 | 0.0418 | Dnaja3 |
| -2.993 | 0.00464 | 0.0422 | Surf1 |
| -2.964 | 0.00501 | 0.0415 | Akap5 |
| -2.866 | 0.00648 | 0.049 | Slc25a34 |
| -2.84 | 0.00695 | 0.0515 | Susd1 |
| -2.819 | 0.00735 | 0.0549 | Itch |
| -2.813 | 0.00745 | 0.0543 | Gm826 |
| -2.764 | 0.00846 | 0.0592 | Gzf1 |
| -2.743 | 0.00895 | 0.0608 | Lztfl1 |
| -2.734 | 0.00915 | 0.0617 | Fastkd1 |
| -2.676 | 0.0106 | 0.0688 | Dpy19l1 |
| -2.655 | 0.0112 | 0.0706 | C1qtnf1 |
| -2.612 | 0.0125 | 0.0767 | Il12rb1 |
| -2.578 | 0.0136 | 0.0776 | Slc25a13 |
| -2.569 | 0.0139 | 0.0787 | Tvp23b |
| -2.563 | 0.0141 | 0.0794 | Ppara |

| | | | |
|---|---|---|---|
| -2.542 | 0.0148 | 0.0826 | Tgfbr3 |
| -2.453 | 0.0184 | 0.101 | Mrps31 |
| -2.434 | 0.0193 | 0.104 | Tasor |
| -2.433 | 0.0193 | 0.0987 | Etfrf1 |
| -2.431 | 0.0195 | 0.099 | Tmub1 |
| -2.43 | 0.0195 | 0.104 | U6 |
| -2.403 | 0.0208 | 0.103 | Slc16a10 |
| -2.392 | 0.0213 | 0.111 | Cops2 |
| -2.386 | 0.0217 | 0.106 | Aamdc |
| -2.366 | 0.0227 | 0.109 | Gabarapl2 |
| -2.364 | 0.0228 | 0.115 | Setmar |
| -2.35 | 0.0236 | 0.112 | Zfp709 |
| -2.311 | 0.0259 | 0.12 | Syne3 |
| -2.291 | 0.0271 | 0.123 | Hsdl2 |
| -2.282 | 0.0276 | 0.125 | Nuak2 |
| -2.221 | 0.0319 | 0.137 | Fam216a |
| -2.212 | 0.0325 | 0.139 | Azin1 |
| 2.186 | 0.0345 | 0.145 | Zfp579 |
| 2.21 | 0.0327 | 0.146 | Brd3 |
| 2.219 | 0.032 | 0.138 | Mapk7 |
| 2.241 | 0.0305 | 0.134 | Son |
| 2.255 | 0.0295 | 0.136 | Zdhhc8 |
| 2.282 | 0.0277 | 0.131 | Epb41l1 |
| 2.288 | 0.0273 | 0.13 | Zfp513 |
| 2.293 | 0.027 | 0.123 | Ifrd1 |
| 2.309 | 0.026 | 0.12 | Tmem132a |
| 2.322 | 0.0252 | 0.117 | Safb2 |
| 2.326 | 0.025 | 0.123 | Ptchd1 |
| 2.331 | 0.0246 | 0.121 | Zfp853 |
| 2.334 | 0.0245 | 0.115 | Pcf11 |
| 2.336 | 0.0244 | 0.121 | Parp3 |
| 2.336 | 0.0244 | 0.121 | Vgf |
| 2.37 | 0.0225 | 0.108 | Esyt3 |
| 2.396 | 0.0211 | 0.104 | Mdm2 |
| 2.407 | 0.0206 | 0.108 | Zfhx2 |
| 2.408 | 0.0206 | 0.108 | Pdia2 |
| 2.415 | 0.0202 | 0.107 | Yeats2 |
| 2.417 | 0.0201 | 0.106 | Lrrc6 |
| 2.423 | 0.0198 | 0.106 | Tmem43 |
| 2.459 | 0.0182 | 0.0997 | Tvp23bos |
| 2.489 | 0.0169 | 0.0902 | Nelfb |
| 2.501 | 0.0164 | 0.0932 | Scgb2b3 |
| 2.523 | 0.0155 | 0.0895 | Tgm3 |
| 2.533 | 0.0151 | 0.0877 | Atp6v0a1 |
| 2.539 | 0.0149 | 0.0869 | Airn |
| 2.549 | 0.0146 | 0.0854 | Spag16 |
| 2.551 | 0.0145 | 0.0813 | Phrf1 |
| 2.597 | 0.0129 | 0.0785 | Vps18 |

| | | | |
|---|---|---|---|
| 2.634 | 0.0118 | 0.0735 | Myo9b |
| 2.636 | 0.0117 | 0.0734 | Nsfl1c |
| 2.646 | 0.0114 | 0.0703 | Maz |
| 2.673 | 0.0107 | 0.0673 | Tuba3a |
| 2.702 | 0.00993 | 0.0659 | Resf1 |
| 2.74 | 0.00902 | 0.0611 | Ppm1n |
| 2.761 | 0.00853 | 0.0595 | Dazap1 |
| 2.761 | 0.00853 | 0.0592 | Slc1a6 |
| 2.784 | 0.00805 | 0.0568 | Foxb1 |
| 2.792 | 0.00788 | 0.0564 | Tex11 |
| 2.797 | 0.00778 | 0.0564 | Traf2 |
| 2.828 | 0.00717 | 0.0544 | Spef1l |
| 2.859 | 0.00661 | 0.0496 | Trio |
| 2.873 | 0.00638 | 0.0485 | Adprs |
| 2.879 | 0.00627 | 0.0505 | Atcay |
| 2.888 | 0.00613 | 0.0474 | Gm18939 |
| 2.888 | 0.00612 | 0.0474 | Gm17740 |
| 2.894 | 0.00602 | 0.0471 | Gm44937 |
| 2.897 | 0.00598 | 0.0494 | Ighv6-4 |
| 2.932 | 0.00545 | 0.0437 | Zfhx3 |
| 2.959 | 0.00508 | 0.042 | Elovl4 |
| 2.96 | 0.00506 | 0.0418 | Slc22a22 |
| 2.983 | 0.00475 | 0.0426 | Foxp4 |
| 3.039 | 0.00409 | 0.0362 | Fry |
| 3.045 | 0.00402 | 0.0359 | Ptpn21 |
| 3.061 | 0.00386 | 0.0383 | Ces3a |
| 3.064 | 0.00382 | 0.0346 | Jun |
| 3.083 | 0.00363 | 0.0373 | H4c17 |
| 3.093 | 0.00353 | 0.0366 | Arhgdig |
| 3.094 | 0.00352 | 0.0327 | BC026762 |
| 3.098 | 0.00349 | 0.0364 | Chd6 |
| 3.142 | 0.00309 | 0.0339 | U6 |
| 3.174 | 0.00282 | 0.0282 | Pla2g4f |
| 3.25 | 0.00228 | 0.0243 | Reg3g |
| 3.29 | 0.00204 | 0.0226 | Kmt2e |
| 3.328 | 0.00183 | 0.0249 | Zan |
| 3.359 | 0.00168 | 0.0198 | Tcerg1 |
| 3.378 | 0.0016 | 0.0191 | Mtf2 |
| 3.429 | 0.00138 | 0.0171 | Cdk5rap2 |
| 3.438 | 0.00134 | 0.0168 | Zfp618 |
| 3.448 | 0.0013 | 0.0212 | Tonsl |
| 3.462 | 0.00125 | 0.016 | Brd4 |
| 3.497 | 0.00113 | 0.0148 | Phc2 |
| 3.516 | 0.00107 | 0.0142 | Mcm3 |
| 3.54 | 0.001 | 0.0136 | Dppa2 |
| 3.58 | 0.000891 | 0.0168 | Rbbp6 |
| 3.638 | 0.000752 | 0.0111 | Rpp25 |
| 3.676 | 0.000671 | 0.0101 | Ly6g6f |

| | | | |
|---|---|---|---|
| 4.664 | 3.20E-05 | 0.00085 | Pogz |

| GENENAME |
|---|
| SH3-binding kinase 1 |
| Ral GTPase activating protein, alpha subunit 2 (catalytic) |
| scavenger receptor class A, member 5 |
| transmembrane and tetratricopeptide repeat containing 2 |
| acyl-CoA thioesterase 1 |
| argininosuccinate synthetase 1 |
| pyruvate dehyrogenase phosphatase catalytic subunit 2 |
| cytochrome P450, family 4, subfamily a, polypeptide 32 |
| heme oxygenase 2 |
| ADP-ribosylation factor-like 15 |
| cysteine-rich PDZ-binding protein |
| ARP6 actin-related protein 6 |
| arsenic (+3 oxidation state) methyltransferase |
| S-adenosylhomocysteine hydrolase |
| a disintegrin and metallopeptidase domain 23 |
| cytochrome P450, family 17, subfamily a, polypeptide 1 |
| gene rich cluster, C10 gene |
| WW domain binding protein 1 like |
| acyl-CoA thioesterase 12 |
| small integral membrane protein 10 like 1 |
| arylformamidase |
| polycomb group ring finger 5 |
| solute carrier family 35, member G1 |
| zinc finger protein 870 |
| arrier family 25 (mitochondrial carrier, palmitoylcarnitine transporter), mem |
| proteasome (prosome, macropain) subunit, alpha type 6 |
| aminoadipate-semialdehyde synthase |
| ciliary neurotrophic factor receptor |
| tubulin epsilon and delta complex 2 |
| TDP-glucose 4,6-dehydratase |
| fibroblast growth factor 1 |
| DnaJ heat shock protein family (Hsp40) member A3 |
| surfeit gene 1 |
| A kinase (PRKA) anchor protein 5 |
| solute carrier family 25, member 34 |
| sushi domain containing 1 |
| itchy, E3 ubiquitin protein ligase |
| predicted gene 826 |
| GDNF-inducible zinc finger protein 1 |
| leucine zipper transcription factor-like 1 |
| FAST kinase domains 1 |
| dpy-19-like 1 (C. elegans) |
| C1q and tumor necrosis factor related protein 1 |
| interleukin 12 receptor, beta 1 |
| arrier family 25 (mitochondrial carrier, adenine nucleotide translocator), mer |
| trans-golgi network vesicle protein 23B |
| peroxisome proliferator activated receptor alpha |

| |
|---|
| transforming growth factor, beta receptor III |
| mitochondrial ribosomal protein S31 |
| transcription activation suppressor |
| electron transfer flavoprotein regulatory factor 1 |
| transmembrane and ubiquitin-like domain containing 1 |
| U6 spliceosomal RNA |
| solute carrier family 16 (monocarboxylic acid transporters), member 10 |
| COP9 signalosome subunit 2 |
| adipogenesis associated Mth938 domain containing |
| gamma-aminobutyric acid (GABA) A receptor-associated protein-like 2 |
| SET domain without mariner transposase fusion |
| zinc finger protein 709 |
| spectrin repeat containing, nuclear envelope family member 3 |
| hydroxysteroid dehydrogenase like 2 |
| NUAK family, SNF1-like kinase, 2 |
| family with sequence similarity 216, member A |
| antizyme inhibitor 1 |
| zinc finger protein 579 |
| bromodomain containing 3 |
| mitogen-activated protein kinase 7 |
| Son DNA binding protein |
| zinc finger, DHHC domain containing 8 |
| erythrocyte membrane protein band 4.1 like 1 |
| zinc finger protein 513 |
| interferon-related developmental regulator 1 |
| transmembrane protein 132A |
| scaffold attachment factor B2 |
| patched domain containing 1 |
| zinc finger protein 853 |
| PCF11 cleavage and polyadenylation factor subunit |
| poly (ADP-ribose) polymerase family, member 3 |
| VGF nerve growth factor inducible |
| extended synaptotagmin-like protein 3 |
| transformed mouse 3T3 cell double minute 2 |
| zinc finger homeobox 2 |
| protein disulfide isomerase associated 2 |
| YEATS domain containing 2 |
| leucine rich repeat containing 6 (testis) |
| transmembrane protein 43 |
| trans-golgi network vesicle protein 23B, opposite strand |
| negative elongation factor complex member B |
| secretoglobin, family 2B, member 3 |
| transglutaminase 3, E polypeptide |
| ATPase, H+ transporting, lysosomal V0 subunit A1 |
| antisense Igf2r RNA |
| sperm associated antigen 16 |
| PHD and ring finger domains 1 |
| VPS18 CORVET/HOPS core subunit |

| |
|---|
| myosin IXb |
| NSFL1 (p97) cofactor (p47) |
| MYC-associated zinc finger protein (purine-binding transcription factor) |
| tubulin, alpha 3A |
| retroelement silencing factor 1 |
| protein phosphatase, Mg2+/Mn2+ dependent, 1N (putative) |
| DAZ associated protein 1 |
| olute carrier family 1 (high affinity aspartate/glutamate transporter), member |
| forkhead box B1 |
| testis expressed gene 11 |
| TNF receptor-associated factor 2 |
| sperm flagellar 1 like |
| triple functional domain (PTPRF interacting) |
| ADP-ribosylserine hydrolase |
| ataxia, cerebellar, Cayman type |
| predicted gene, 18939 |
| predicted gene, 17740 |
| predicted gene 44937 |
| immunoglobulin heavy variable V6-4 |
| zinc finger homeobox 3 |
| elongation of very long chain fatty acids (FEN1/Elo2, SUR4/Elo3, yeast)-like 4 |
| solute carrier family 22 (organic cation transporter), member 22 |
| forkhead box P4 |
| FRY microtubule binding protein |
| protein tyrosine phosphatase, non-receptor type 21 |
| carboxylesterase 3A |
| jun proto-oncogene |
| H4 clustered histone 17 |
| Rho GDP dissociation inhibitor (GDI) gamma |
| cDNA sequence BC026762 |
| chromodomain helicase DNA binding protein 6 |
| U6 spliceosomal RNA |
| phospholipase A2, group IVF |
| regenerating islet-derived 3 gamma |
| lysine (K)-specific methyltransferase 2E |
| zonadhesin |
| transcription elongation regulator 1 (CA150) |
| metal response element binding transcription factor 2 |
| CDK5 regulatory subunit associated protein 2 |
| zinc finger protein 618 |
| tonsoku-like, DNA repair protein |
| bromodomain containing 4 |
| polyhomeotic 2 |
| minichromosome maintenance complex component 3 |
| developmental pluripotency associated 2 |
| retinoblastoma binding protein 6, ubiquitin ligase |
| ribonuclease P/MRP 25 subunit |
| lymphocyte antigen 6 complex, locus G6F |

| pogo transposable element with ZNF domain |
| --- |

| Biotype |
| --- |
| protein_coding; |
| protein_coding; |
| protein_coding; |
| protein_coding; |
| protein_coding; |
| protein_coding; |
| protein_coding; |
| protein_coding; |
| protein_coding; |
| protein_coding; |
| protein_coding; |
| protein_coding; |
| protein_coding; |
| protein_coding; |
| protein_coding; |
| protein_coding; |
| protein_coding; |
| protein_coding; |
| protein_coding; |
| protein_coding; |
| protein_coding; |
| protein_coding; |
| protein_coding; |
| protein_coding; |
| protein_coding; |
| protein_coding; |
| protein_coding; |
| protein_coding; |
| protein_coding; |
| protein_coding; |
| protein_coding; |
| protein_coding; |
| protein_coding; |
| protein_coding; |
| protein_coding; |
| protein_coding; |
| protein_coding; |
| protein_coding; |
| protein_coding; |
| protein_coding; |
| protein_coding; |
| protein_coding; |
| protein_coding; |
| protein_coding; |
| protein_coding; |
| protein_coding; |
| protein_coding; |

| |
|---|
| protein_coding; |
| protein_coding; |
| protein_coding; |
| protein_coding; |
| protein_coding; |
| snRNA; |
| protein_coding; |
| protein_coding; |
| protein_coding; |
| protein_coding; |
| protein_coding; |
| protein_coding; |
| protein_coding; |
| protein_coding; |
| protein_coding; |
| protein_coding; |
| protein_coding; |
| protein_coding; |
| protein_coding; |
| protein_coding; |
| protein_coding; |
| protein_coding; |
| protein_coding; |
| protein_coding; |
| protein_coding; |
| protein_coding; |
| protein_coding; |
| protein_coding; |
| protein_coding; |
| protein_coding; |
| protein_coding; |
| protein_coding; |
| protein_coding; |
| protein_coding; |
| protein_coding; |
| protein_coding; |
| protein_coding; |
| protein_coding; |
| protein_coding; |
| lncRNA; |
| protein_coding; |
| protein_coding; |
| protein_coding; |
| protein_coding; |
| lncRNA; |
| protein_coding; |
| protein_coding; |
| protein_coding; |

| |
|---|
| protein_coding; |
| protein_coding; |
| protein_coding; |
| protein_coding; |
| protein_coding; |
| protein_coding; |
| protein_coding; |
| protein_coding; |
| protein_coding; |
| protein_coding; |
| protein_coding; |
| protein_coding; |
| protein_coding; |
| protein_coding; |
| protein_coding; |
| transcribed_processed_pseudogene; |
| processed_pseudogene; |
| TEC; |
| IG_V_gene; |
| protein_coding; |
| protein_coding; |
| protein_coding; |
| protein_coding; |
| protein_coding; |
| protein_coding; |
| protein_coding; |
| protein_coding; |
| protein_coding; |
| protein_coding; |
| lncRNA; |
| protein_coding; |
| snRNA; |
| protein_coding; |
| protein_coding; |
| protein_coding; |
| protein_coding; |
| protein_coding; |
| protein_coding; |
| protein_coding; |
| protein_coding; |
| protein_coding; |
| protein_coding; |
| protein_coding; |
| protein_coding; |
| protein_coding; |
| protein_coding; |
| protein_coding; |
| protein_coding; |

| |
|---|
| protein_coding; |
