## Supplementary material for "Blocking necroptosis reduces inflammation and tumor incidence in a mouse model of diet-induced hepatocellular carcinoma": Table S2

**Table S2: List of primers used for RT-PCR.**

| Gene | Forward sequence | Reverse sequence |
|---|---|---|
| *Acta-2* | 5'-CTGACAGAGGCACCACTGAA-3' | 5'-CATCTCCAGAGTCCAGCACA-3' |
| *AFP* | 5'- ACAGGAGGCTATGCATCACC-3' | 5'- TGGACATCTTCACCATGTGG-3' |
| *β-microglobulin* | 5′-CACTGACCGGCCTGTATGC-3′ | 5′-GGGTGGCGTGAGTATACTTGAAT-3′ |
| *CCL2* | 5'-TTAAAAACCTGGATCGGAACCAA-3' | 5'-GCATTAGCTTCAGATTTACGGGT-3' |
| *Cdkn2a(p16Ink4a)* | 5'-CCCAACGCCCCGAACT-3' | 5'-GCAGAAGAGCTGCTACGTGAA-3' |
| *Cdkn1a(p21Cip1)* | 5'-GTCAGGCTGGTCTGCCTCCG-3' | 5'-CGGTCCCGTGGACAGTGAGCAG-3' |
| *CD68* | 5'-CCACAGGCAGCACAGTGGAC-3' | 5'-TCCACAGCAGAAGCTTTGGCCC-3' |
| *c-Myc* | 5'-AGTGCTGCATGAGGAGACAC-3' | 5'-GGTTTGCCTCTTCTCCACAG-3' |
| *Col1α1* | 5'-GCTCCTCTTAGGGGCCACT-3' | 5'-CCACGTCTCACCATTGGGG-3' |
| *Col3α1* | 5'-CTGTAACATGGAAACTGGGGAAA-3' | 5'- CCATAGCTGAACTGAAAACCACC-3' |
| *Dppa2* | 5′-ATGTCATACTTCGGCCTGGAGAC-3′ | 5′-GGACCCTGCTTCATTCTGGCCTC-3′ |
| *EPCAM* | 5'-CGTGAGGACCTACTGGATCAT -3' | 5'- GTCCACGTCGTCTTGTGTTTT -3' |
| *HPRT* | 5'-CTGGTGAAAAGGACCTCTCG-3' | 5'-TGAAGTACTCATTATAGTCAAGGGCA-3' |
| *IL-6* | 5'-TGGTACTCCAGAAGACCAGAGG-3' | 5'-AACGATGATGCACTTGCAGA-3' |
| *IL-1β* | 5'-AGGTCAAAGGTTTGGAAGCA-3' | 5'-TGAAGCAGCTATGGCAACTG-3' |
| *MLKL* | 5′-CTGAGGGAACTGCTGGATAGAG-3′ | 5′-CGAGGAAACTGGAGCTGCTGAT-3′ |
| *p53* | 5'-GTATTTCACCCTCAAGATCC-3' | 5'-TGGGCATCCTTTAACTCTA-3' |
| *Pogz* | 5'-TGCACTTCGTGTACCTTTGC-3' | 5'-AGGCCTCAAGTGAGACATCC-3' |
| *RIPK3* | 5′-GAAGACACGGCACTCCTTGGTA-3′ | 5′-CTTGAGGCAGTAGTTCTTGGTGG-3′ |

| | | |
|---|---|---|
| *TGFβ* | 5'-ACCATGCCAACTTCTGTCTGGGAC-3' | 5'-ACAACTGCTCCACCTTGGGCTTG-3' |
| *TLR4* | 5'-ATGGCATGGCTTACACCACC-3' | 5'-GAGGCCAATTTTGTCTCCACA-3' |
| *TNFα* | 5'-CACAGAAAGCATGATCCGCGACGT-3' | 5'- CGGCAGAGAGGAGGTTGACTTTCT-3' |
| *TONSL* | 5'-TGAGAAACTAGAGGGGATGCTG-3' | 5'-AGGTGAGACCCAGATTGAGGTA-3' |
